# FASN Inactivation-Induced Progranulin (GRN) Expression Promotes Lysosome-Dependent Cell Death to Suppress Leukemogenesis

**DOI:** 10.1101/2025.05.06.652401

**Authors:** Meng Su, Zhiyi Lv, Yating Lu, Xinshu Xie, Ailin Zou, Hanqi Liu, Jie Ouyang, Qinglin Li, Xuezhen Ma, Yexin Yang, Kang Liu, Yaohui He, Shuqian Xu, Ji Li, Lili Chen, Xin Lu, Jie Yang, Caiming Wu, Lu Liu, Lei Zhang, Yue Sheng, Yong Huang, Yang Mei

## Abstract

Cancer cells rely on lipogenesis in addition to exogenous lipid uptake, and fatty acid synthase (FASN) is aberrantly overexpressed in myeloid leukemia. However, the precise role of FASN in leukemogenesis *in vivo* remains elusive. Here, we demonstrated that FASN is essential for leukemogenesis. Ablation of FASN impeded leukemic cell growth, survival, clonogenicity *in vitro*, and the leukemic burden *in vivo*. Conditional knockout of FASN barely affected hematopoiesis but significantly attenuated the leukemic progression of MLL-AF9 transplants. By screening a library of platensimycin derivatives, we identified compound MS-C19 as a potent FASN inhibitor. Pharmacological targeting of FASN by MS-C19 suppressed leukemic cell growth and clonogenicity in clinical AML blasts. Mechanistically, both MS-C19 treatment and FASN deficiency triggered the activation of lysosomal and inflammatory gene expression. Loss of FASN led to lysosomal membrane permeabilization and subsequent lysosome-associated cell death, but not lysosome biogenesis. We further identified that *GRN*, a lysosomal and neuroinflammatory gene, was potently transcribed by TFEB upon FASN loss. Depletion of *GRN* significantly reversed the inhibitory effects caused by FASN knockdown. Our work demonstrates that FASN is a therapeutic target for myeloid leukemia, and inactivation of FASN by the lead compound MS-C19 provides an alternative approach for leukemic intervention.

**Key Points:**

1. Genetic ablation of FASN induces apoptosis, impedes leukemia cell growth, and improves the survival of leukemic xenografts.
2. Targeting FASN by the compound MS-C19, a derivative of the natural microbe antibiotic platensimycin, reduces the cell survival of leukemic cell lines, the clinical leukemic blasts, and MLL-AF9 transformed murine leukemic cells.
3. FASN inhibition induces lysosome-dependent cell death that relies on Progranulin (*GRN*) expression.

## Introduction

Recent evidence has highlighted the vital roles of dysregulated cellular metabolism in initiating and promoting leukemic progression, of which fatty acid synthesis became intriguing ^1–4^. In addition to taking up exogenous lipids, cancer cells generate fatty acids from alternative carbon sources through de novo lipogenesis ^5,6^, and fatty acid synthase (FASN) is one core enzyme that engages in cyclical reactions to produce palmitate. FASN expression is upregulated in solid cancers and provides survival advantages for cancer cells ^7–13^. Clinical studies also linked fatty acid metabolism with prognosis, immunosurveillance, and tumor microenvironments in acute lymphoblastic leukemia (ALL) and acute myeloid leukemia (AML) ^14–16^. Inhibiting fatty acid metabolism and homeostasis has emerged as a promising anti-leukemia strategy ^2–4,17–20^, whereas controversial findings also existed ^21^. FASN was previously identified as a poor prognostic factor for ALL^14^, and depletion of FASN-sensitized all-trans retinoic acid (ATRA)-induced acute promyelocytic leukemia (APL) cell differentiation *in vitro* ^17^. However, whether FASN is essential and targetable in leukemogenesis *in vitro* and *in vivo* remains largely unknown.

Several FASN inhibitors have been developed and entered preclinical trials. However, despite decades of efforts in targeting FASN, only Orlistat has received FDA approval as an anti-obesity medication ^22^. None of the FASN inhibitors have been formally used for cancer treatment, particularly hematologic malignancies ^5,6^. Platensimycin (PTM) is a novel natural antibiotic produced by *Streptomyces platensis*, it was shown to have selective inhibition of mammalian FASN in diabetes ^23^. Whether FASN inhibitors based on PTM could efficiently target leukemia cells is unclear.

Here, we demonstrate that aberrantly overexpressed FASN is associated with poor survival of AML patients. Genetically depletion of FASN largely impeded the leukemic cell growth and clonogenicity. Although ablation of *Fasn in vivo* is largely dispensable for normal hematopoiesis, its deficiency significantly impairs the leukemogenesis induced by oncogene expression. We screened a PTM derivatives compound library and identified MS-C19 as a potential FASN inhibitor. Pharmacological inhibition of FASN by MS-C19 is sufficient to induce apoptosis and promote cell differentiation of leukemic cells. We further revealed that FASN inhibition led to TFEB-orchestrated lysosomal gene activation but contributed minimally to the lysosome biogenesis. FASN deficiency disrupted the lysosome membrane permeabilization and triggered lysosome-dependent cell death. Blocking lysosomal activity either by bafilomycin A1 treatment or *GRN* silencing reverts the apoptosis caused by FASN inhibition. Our work demonstrates that targeting FASN is an effective avenue for therapeutically killing leukemic cells. Also, it provides insights into the development of novel FASN inhibitors from natural products.

## Results

### FASN is aberrantly overexpressed in AML patients

*FASN* mRNA levels were significantly elevated in AML patients with different karyotypes (Fig.1A) ^24^. We validated the finding using an independent AML cohort (Fig.1B). Using the online Kaplan-Meier plotter tool (KMPlot), we uncovered that high FASN expression is associated with a poor overall (P=0.081) and event-free (P=0.019) survival of untreated AML patients (Fig.1C). This was further supported by analyzing the TCGA dataset, in which FASN expression inversely correlated with patient overall survival (P=0.04) (Fig.1D). These results suggest that FASN is aberrantly overexpressed in AML patients and negatively correlates with patient survival.

**Fig.1.**
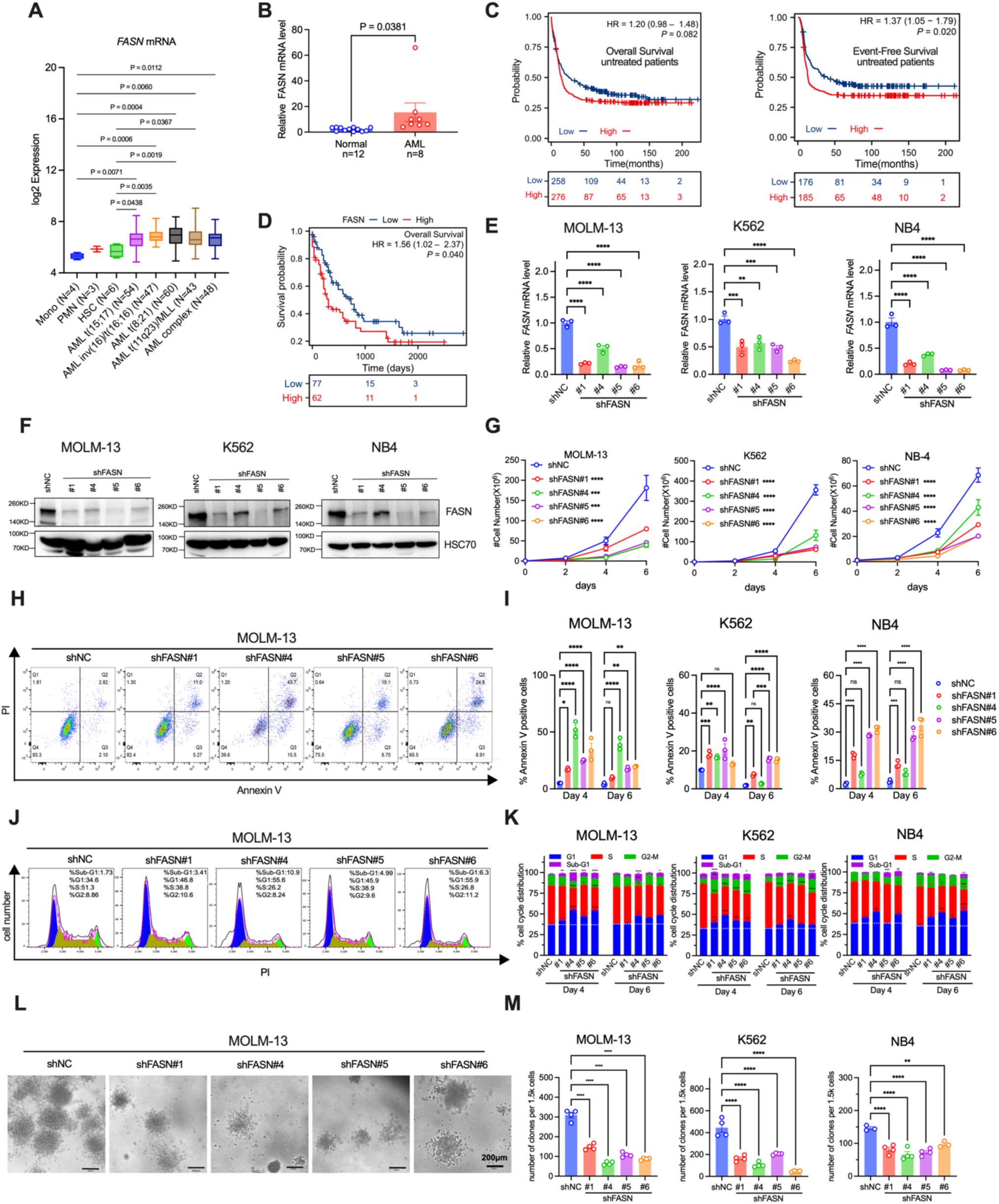
Increased FASN expression is associated with a poor diagnosis in AML patients and knockdown of FASN impairs leukemic cell growth *in vitro*. (**A**) FASN expression in AML bone marrow blasts compared to HSC (Lin-CD34+ CD38-CD90+ CD45RA-), PMN (Polymorphonuclear cells), or Mono (Monocytes, CD14+ CD16-) from healthy donors. AML complex, AML with complex aberrant karyotype. Statistical analysis was performed by one-way ANOVA with Tukey’s corrections. (**B**) qPCR detecting *FASN* mRNA levels in de novo AML cohorts using 18s rRNA as the internal control. An unpaired student t-test was performed with two-tailed. (**C**) Kaplan-Meier survival curves of untreated AML patients with high or low FASN expression were generated using public online KMPlotter. (**D**) Similar to C, except the data were scored from the TCGA database. (**E-F**) FASN mRNA (E) or protein (F) levels assessed d by qPCR or western blotting from indicated cells transduced with negative control (shNC) or constructs expressing FASN shRNA. (**G**) Growth curve of indicated leukemic cell lines with either intact FASN expression or FASN depletion by shRNAs. (**H**-**I**) Apoptotic cells in G were assayed by annexin V staining. Representative flow cytometric plots were shown in H, and apoptotic cell percentages were quantitated in I. (**J**-**K**) Cell cycle profiling in control and FASN-depleted cells. Representative flow cytometric plots and cell cycle distributions were further shown in J and K, respectively. (**L**-**M**) Clonogenicity analysis of indicated cells. Representative colony images were shown in L, and colony counts were quantitated in M. All experiments were performed in biological triplicates, except for quadruplicates in M as indicated. The statistical analysis was performed by one-way ANOVA with Tukey’s corrections, except two-way ANOVA with Tukey’s corrections in I and K. *, P<0.05. **, P<0.01. ***, P<0.001. ****, P<0.0001.

### Impaired FASN expression impeded the growth and survival of human leukemic cell lines

To examine whether FASN is required for leukemic cell growth and survival *in vitro*, we knocked down FASN using FASN-specific shRNA in human leukemic cell lines (Fig.1E-F, and fig. S1A-B). Depletion of FASN significantly attenuated the cell growth of AML cells and induced cell apoptosis (Fig.1G-I). The cell cycle progressions were also interrupted by FASN deficiency (Fig.1J-K). Consistently, the colony-forming ability of all the tested leukemic cells with reduced FASN expression was decreased (Fig.1L-M).

We also evaluated the impact of FASN knockout on leukemic cell growth and survival using CRISPR-Cas9 genome editing. Three independent single guide RNAs (sgRNA) directly targeting the human FASN genome were introduced (fig. S2A), and single cell-derived *FASN^-/-^* clones were selected from MOLM-13 cells expressing either single sgRNA or their combinations. We verified the deletion of *FASN* in single KO clones (Fig.2A and fig. S2B-S2C). Accordingly, *FASN^-/-^* cells exhibited almost completely blocked cell growth, paralleled by accumulated apoptosis, reduced S-phase entry, and compromised colony formation abilities (Fig.2B-E).

**Fig.2.**
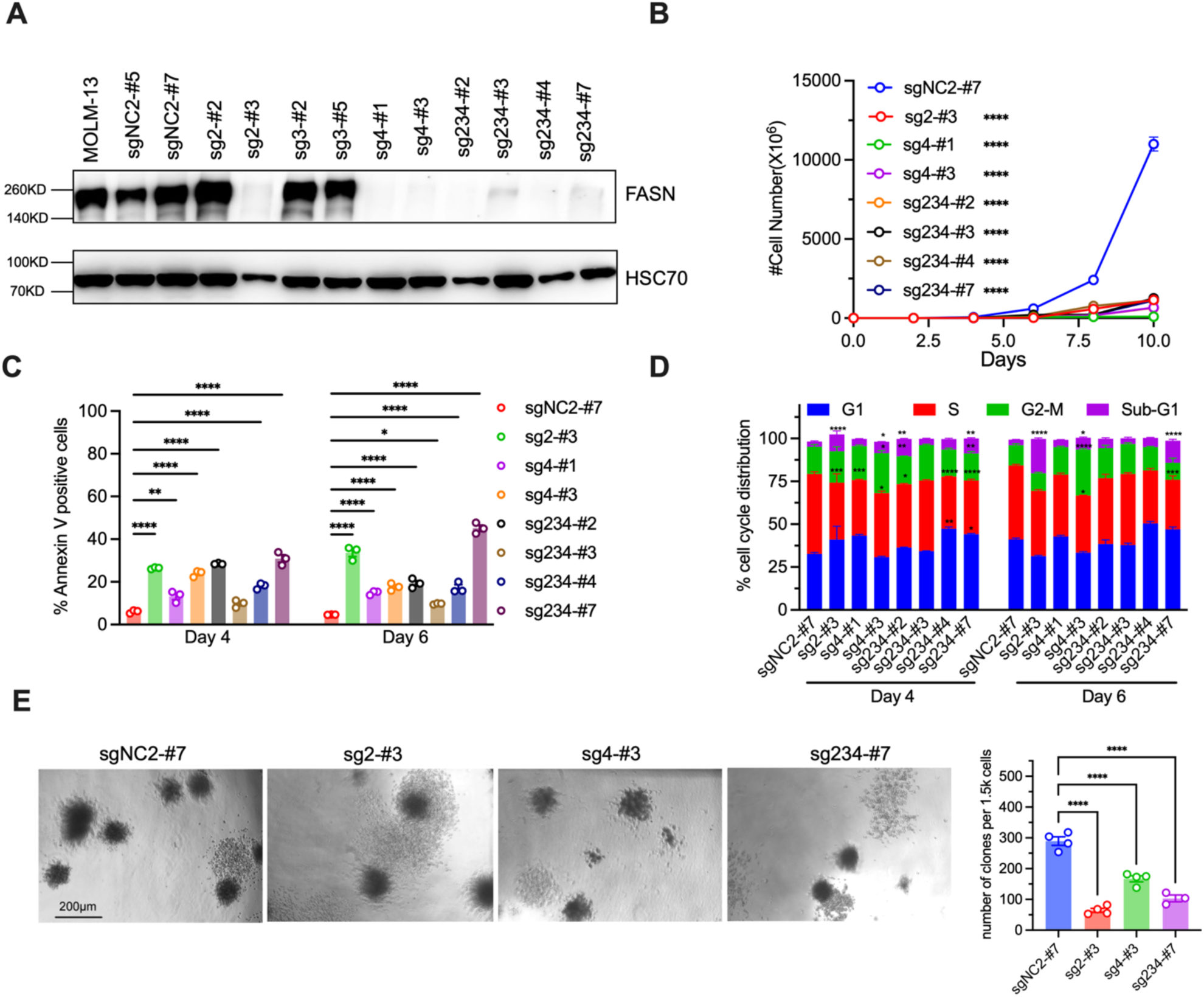
FASN depletion by CRISPR-Cas9 suppresses leukemia cell growth *in vitro*. (**A**) FASN-knockout MOLM-13 cells isolated from single-colony were verified by western blotting. (**B**-**D**) The cell growth curves, apoptosis, and cell cycle analysis of indicated cells in A were shown in B, C, and D, respectively. (**E**) Colony formation assays were performed using indicated cells. Representative images were shown on the left and average colony numbers were shown on the right. All experiments were conducted in biological triplicates, except quadruplicates in E. The statistical analysis was performed by one-way ANOVA in B and E, and two-way ANOVA in C and D with Tukey’s corrections for all. *, P<0.05. **, P<0.01. ***, P<0.001. ****, P<0.0001.

Altogether, our data demonstrated that impaired FASN expression largely impeded cell growth, cell cycle progression, cell survival, and clonogenicity of human leukemic cells *in vitro*.

### FASN is required for leukemogenesis *in vivo*

To further elucidate the roles of FASN in leukemogenesis *in vivo*, we transplanted either *FASN* knockdown or control MOLM-13 cells into immunodeficient mice. Loss of *FASN* significantly prolonged survival of recipients bearing xenografts (Fig.3A). By monitoring the leukemic burden using non-invasive imaging of bioluminescence, we observed markedly inhibited progression of human leukemic cells harboring luciferase *in vivo* upon FASN depletion, accompanied by significantly reduced engraftment of human CD45^+^ cells in the BM and spleen of recipients (Fig.3B-C). Consistently, FASN KO in MOLM-13 cells delayed the mortality in xenograft-bearing mice (Fig.3D).

**Fig.3.**
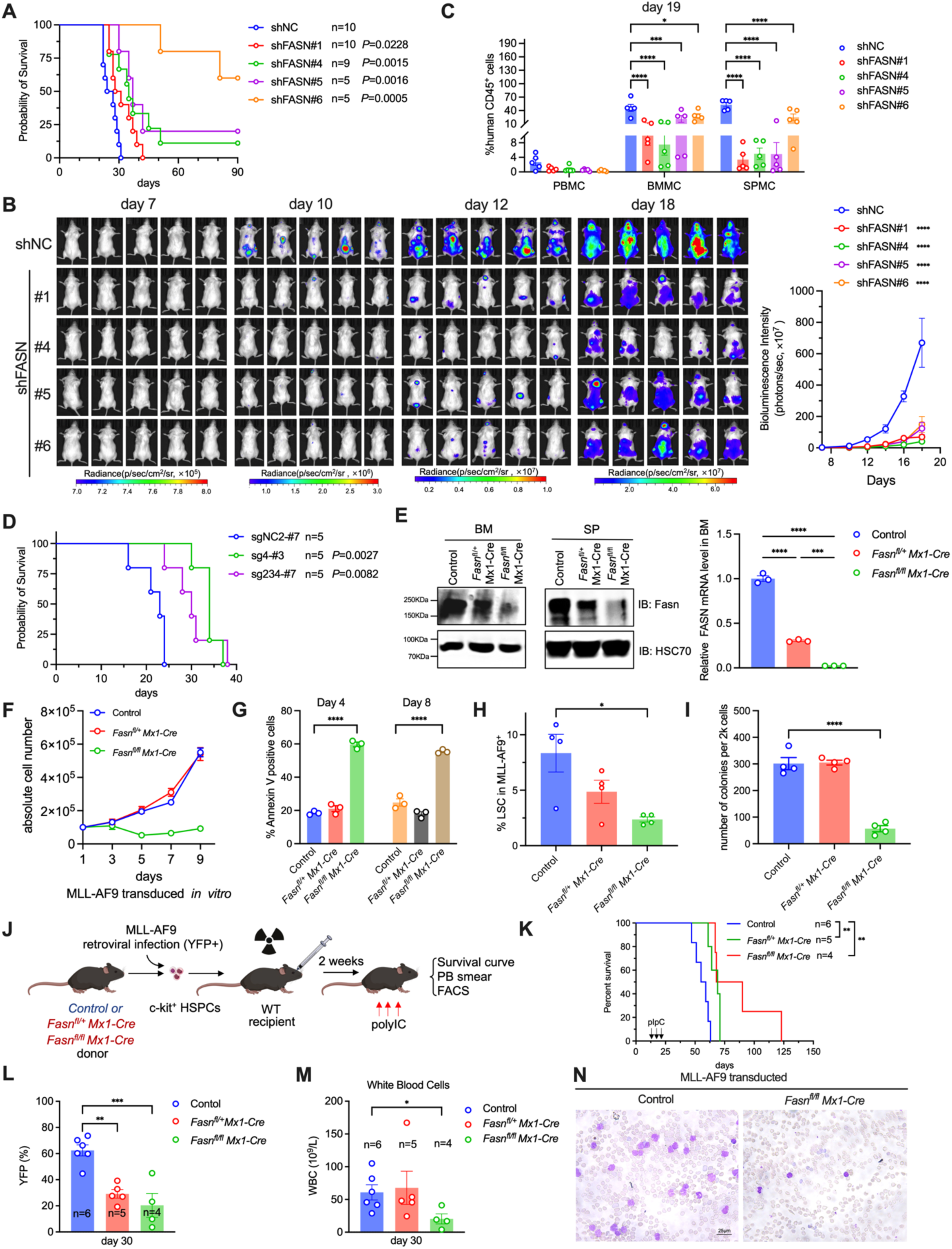
FASN deficiency inhibits leukemic progress *in vivo*. (**A**) Kaplan-Meier survival curves for immune-deficient NSGS mice transplanted with control or FASN KD MOLM-13 cells. (**B**) IVIS imaging deciphering the *in vivo* leukemic progression of indicated cells transplanted into immunodeficient NKG mice. (**C**) The same mice from B were sacrificed and the leukemic blast cells were assessed by hCD45 staining followed by flow cytometric analysis. (**D**) Survival curves of NSGS transplants receiving control or FASN null MOLM-13 cells. (**E**) Western blotting (upper) and qPCR (lower) confirmation of Fasn depletion in bone marrow and spleen from indicated mice. (**F-I**) The c-kit^+^ BM cells from control or Fasn-deficient mice were retrovirally transduced to express MLL-AF9. Two weeks post-infection, the cell growth curve (F), apoptosis (G), LSC frequency (H), and clonogenicity (I) were examined. The experiments were performed in triplicates, except quadruplicates in I. (**J**) Schematic diagram illustrating the establishment of MLL-AF9 induced control or Fasn deficient leukemic mouse models. (**K**) Kaplan-Meier curves for survival of animals after polyIC injections (6µg/g, three doses, every other day). (**L**) Peripheral YFP-positive leukemic blasts were determined by flow cytometric assay 30 days post-transplantation. (**M**) Peripheral white blood cell counts from indicated transplants. (**N**) Leukemic blast cells were visualized by peripheral blood smear with Giemsa staining. Two-way ANOVA in B and C, One-way ANOVA in E, G, H, I, and L with Tukey’s corrections in all; Unpaired student t-test in M; Mantel-Cox (Log-Rank) statistical analysis in A, D, and K. *, P<0.05. **, P<0.01. ***, P<0.001. ****, P<0.0001.

Next, we tested if FASN deficiency prevented the occurrence or progression of leukemia induced by ectopically expressed MLL-AF9 in murine hematopoietic cells *in vivo*. Conditional *Fasn* knockout mice were crossed with Mx1-Cre to generate *Fasn*^fl/+^ Mx1-Cre and *Fasn*^fl/fl^ Mx1-Cre mice. We confirmed that three consecutive doses of polyIC induced partial and complete removal of Fasn expression in *Fasn*^fl/+^ Mx1-Cre and *Fasn*^fl/fl^ Mx1-Cre mice, respectively (Fig.3E). We next retrovirally introduced the MLL-AF9 oncogene into c-kit^+^ HPSCs after polyIC administration, followed by continues *in vitro* culture to achieve YFP^+^ cells (fig. S3A). Compared with transformed control cells, Fasn null cells expressing MLL-AF9 showed lower growth rate, increased apoptosis and differentiation, and impaired cell proliferation, leukemic stem cell (LSCs) frequency, and colony formation (Fig.3F-I, and fig. S3B-C). The MLL-AF9 oncogene was also retrovirally introduced into c-kit^+^ HPSCs with intact Fasn expression, followed by transplantation into recipient mice (fig. S3D). After 2 weeks post-transplantation, the recipients were challenged with polyIC to induce the genomic excision of *Fasn* (Fig.3J). We observed significant delayed mouse mortality, paralleled with decreased YFP^+^ leukemic cells and white blood cell counts in PB of Fasn deficient transplants (Fig.3K-N). To eliminate the influence dictated by potential variability in viral transduction efficacy, we transplanted the *in vitro* established MLL-AF9 transformed YFP^+^ cells with wild-type supportive BM cells into the recipients (fig. S3D). The transplants reconstituted by *Fasn* heterozygous and homozygous cells expressing MLL-AF9 had delayed disease progression compared with recipients receiving transduced control cells (fig. S3E). Accordingly, loss of Fasn reduced the leukemic burden as evidenced by the significantly decreased leukemic blasts and colony-forming units in the PB, as well as reduced frequency and numbers of LSCs in the BM and SP of *Fasn^fl/fl^ Mx1-Cre* transplants (fig. S3F-J). Altogether, these results indicate that FASN is required for leukemogenesis *in vivo*.

### Fasn is dispensable for normal murine hematopoiesis and HSC maintenance

Next, we evaluated the *in vivo* function of Fasn for normal hematopoiesis. The complete blood cell counts, spleen size, and cellularity of BM and spleen as determined one month after Fasn deletion gave no apparent difference in all the mice regardless of Fasn expression (Fig.4A-C). Maturate hematopoietic cells developed normally in Fasn deficient mice (Fig.4D-E, and fig. S4A). In addition, the lack of Fasn neither influenced the maintenance of HSC (SLAM-LSK) (Fig.4F-G, and fig. S4B) and long-term HSC (LT-HSC) (fig. S4C-E) nor the compositions of lineage-committed progenitors (CMP, GMP, MEP, and CLP) in mice (Fig.4H-I, and fig. S4F). When a competitive repopulation assay was performed to investigate whether loss of *Fasn* impaired the reconstitution ability of HSPCs *in vivo*, we found all the donor-derived cells exhibited similar percentages in the PB of transplants (4-16 weeks) (Fig.4J). Fasn-deficient HSCs had comparable repopulation abilities in PB, BM, and spleen for all three lineages (Fig.4K-M). Chimerism analyses of BM HSCs from the recipient mice revealed that Fasn depletion had minimal effects on the maintenance of HSCs and committed progenitors *in vivo* (Fig.4N-O). The competitive reconstitution assay was also performed using cells with intact Fasn before polyIC treatments, and genomic excision was induced by three consecutive polyIC administrations after 4 weeks. As expected, the levels of donor-cell reconstitution barely differed between control or Fasn-deficient cells up to 16 weeks (Fig.4P). There were also indistinguishable differences in BM HSC pool among all the recipients (Fig.4Q). Nonetheless, *Fasn* had been efficiently deleted in LSK cells after polyIC injections in the tested conditions (fig. S4G). These results collectively demonstrate that HSCs do not require Fasn for self-renew or differentiation under either steady-state or long-term reconstitution conditions, and Fasn inactivation is largely unharmful to normal hematopoiesis.

**Fig.4.**
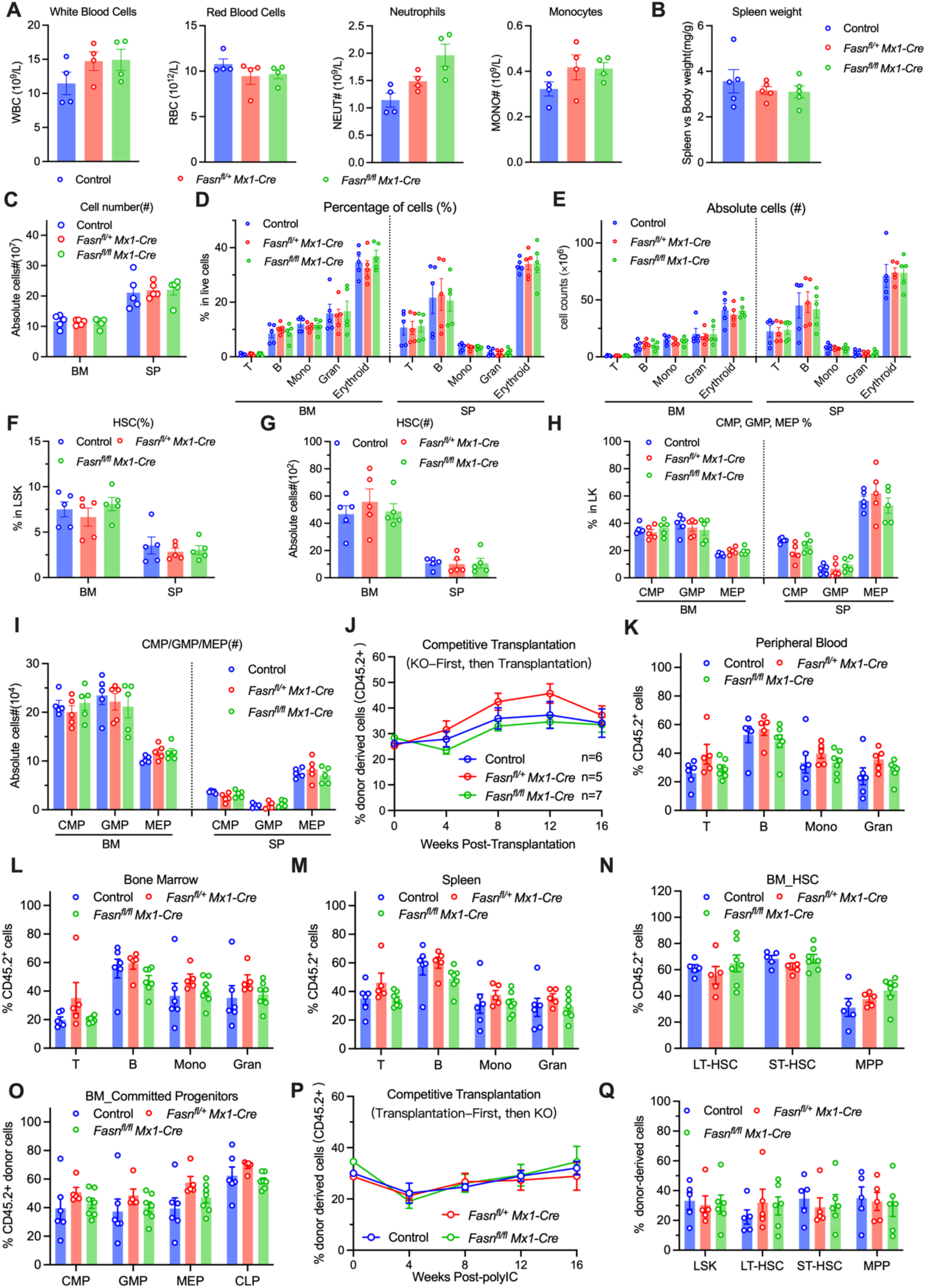
Conditional Fasn deletion does not disturb normal hematopoiesis and HSC maintenance in adult mice. (**A**) Complete blood counts of indicated mice 30 days post polyIC i.p. injection. (**B**) The spleen weights of mice in A were measured and were quantitated as the relative ratio to the body weight. (**C**) Absolute cells in bone marrow and spleen from mice in A. (**D**-**E**) Percentages and absolute cell numbers of different lineage-positive cells from indicated mice as determined by flow cytometric assay were shown in D and E, respectively. T, CD3e^+^; B, B220^+^; Mono, Ly6G^-^ CD11b^+^; Gran, Ly6G^+^CD11b^+^; Erythroid, TER119^+^. (**F**-**G**) Percentages and absolute cell numbers of HSC were examined in F and G respectively. HSC, CD150^+^CD48^-^Lin^-^Sca1^+^c-Kit^+^. (**H**-**I**) Same as in F and G, except GMP, CMP, and MEP were determined. GMP, CD34^+^CD16/32^+^Lin^-^c-Kit^+^; CMP, CD34^+^CD16/32^-^Lin^-^c-Kit^+^; MEP, CD34^-^CD16/32^-^Lin^-^c-Kit^+^. n=4 in each group for A, n=5 in each group for B-I. (**J**) Chimerism analyses of competitive bone marrow transplants. 1×10^6^ donor bone marrow mononuclear cells (CD45.2+, after polyIC treatment) with equal competitors (CD45.1^+^) were retro-orbitally injected into CD45.1^+^ recipient mice, peripheral blood bleeding was conducted at the indicated time window followed by flow cytometric analysis. (**K**-**M**) The compositions of donor-derived maturation cells in peripheral blood (PB), bone marrow (BM), and spleen (SP) from indicated transplants in J at week 16 were respectively assessed in K, L, and M. (**N-O**) Donor constitutions in bone marrow HSPCs (N) and committed progenitors (O) of transplants from J were analyzed at week 16. For J-O, Control, n=6; *Fasn^fl/+^ Mx1-Cre*, n=5; *Fasn^fl/fl^ Mx1-Cre*, n=7. LT-HSC, long-term HSCs, CD34^-^CD135^-^LSK; ST-HSC, short-term HSCs, CD34^+^CD135^-^LSK; MPP, multipotent progenitors, CD34^+^CD135^+^LSK; CLP, common lymphoid progenitors, CD127^+^Lin^-^Sca1^mid^c-kt^mid^. (**P**-**Q**) Similar as in J, except donor cells were intact before transplantation. Fasn deletion was then achieved by polyIC injection 4 weeks post-transplantation. Chimerism analyses of PB bleeding at indicated time points and bone marrow HSPCs at week 16 were shown in P and Q, respectively. Control, n=5; *Fasn^fl/+^ Mx1-Cre*, n=5; *Fasn^fl/fl^ Mx1-Cre*, n=6.

### Targeted screening of FASN inhibitors from Platensimycin (PTM) derivatives

We envision an approach to discover novel FASN inhibitors with anti-leukemia potential from a platensimycin triazole library, prepared via a Cu(I)-catalyzed click chemistry (fig. S5). We screened the focused library of 70 PTM derivatives, paralleled with PTM and Orlistat, against K562 cells (Fig.5A-B). Compound 19 showed profound inhibitory activity, which is more potent than Orlistat (Fig.5B). Unexpectedly, the parent PTM had little if any suppressive effect on leukemic cell growth (Fig.5B). The compound **19**, referred hereafter as MS-C19, exhibited substantially lower IC50 over Orlistat and TVB-3166 in K562 and MOLM-13 cells (Fig.5C). Accordingly, MOLM-13 and K562 cells responded dose-dependently to MS-C19 in the aspects of reduced cell growth, increased cell apoptosis, declined S phase or accumulated G2/M phase cells and compromised colony formation abilities, which were either barely or occasionally observed with PTM, Orlistat or TVB-3166 treatment (Fig.5D-G, and fig. S6A). Moreover, when treating the MLL-AF9 murine leukemic cells with Fasn inhibitors, we constantly observed attenuated cell growth and clonogenicity of cells treated with TVB-3166, Orlistat, and MS-C19 (Fig.5H-J). Strikingly, MS-C19 induced myeloid or erythroid differentiation of treated cells (Fig.5K-O). Inhibition of Fasn by TVB-3166 or MS-C19, but not Orlistat or PTM, also promoted the myeloid maturation of MLL-AF9 transformed murine leukemia cells (Fig.5P, and fig. S6B-C). MS-C19-induced myeloid differentiation was further confirmed by nitroblue tetrazolium (NBT) staining (fig. S6D-E). More specifically, we observed dose- and time-dependent reduction of FASN protein levels in MS-C19 treated MOLM-13 (Fig.5Q), which was absent in Orlistat or TVB-3166 treated cells (fig. S7A-B).

**Fig.5.**
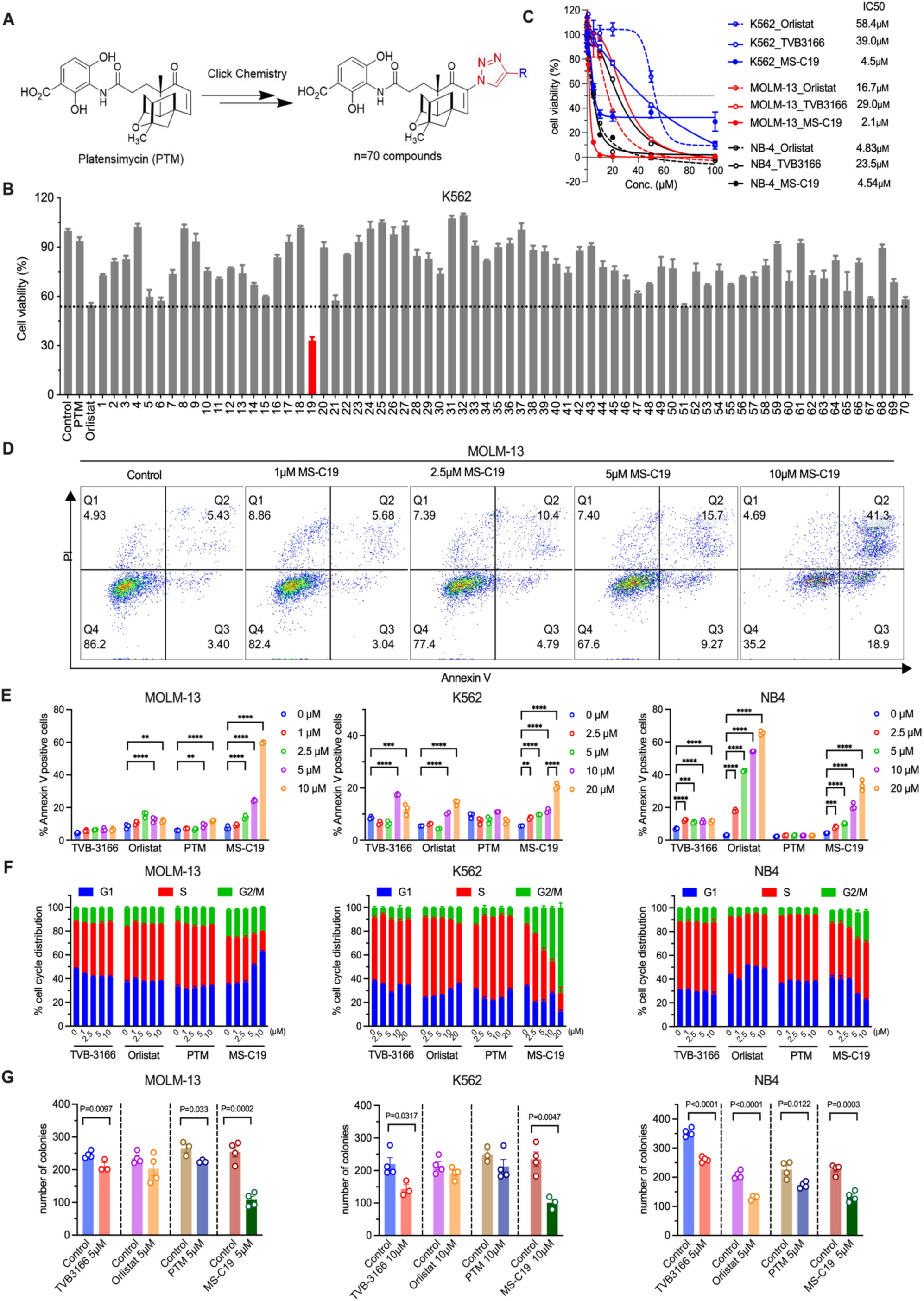
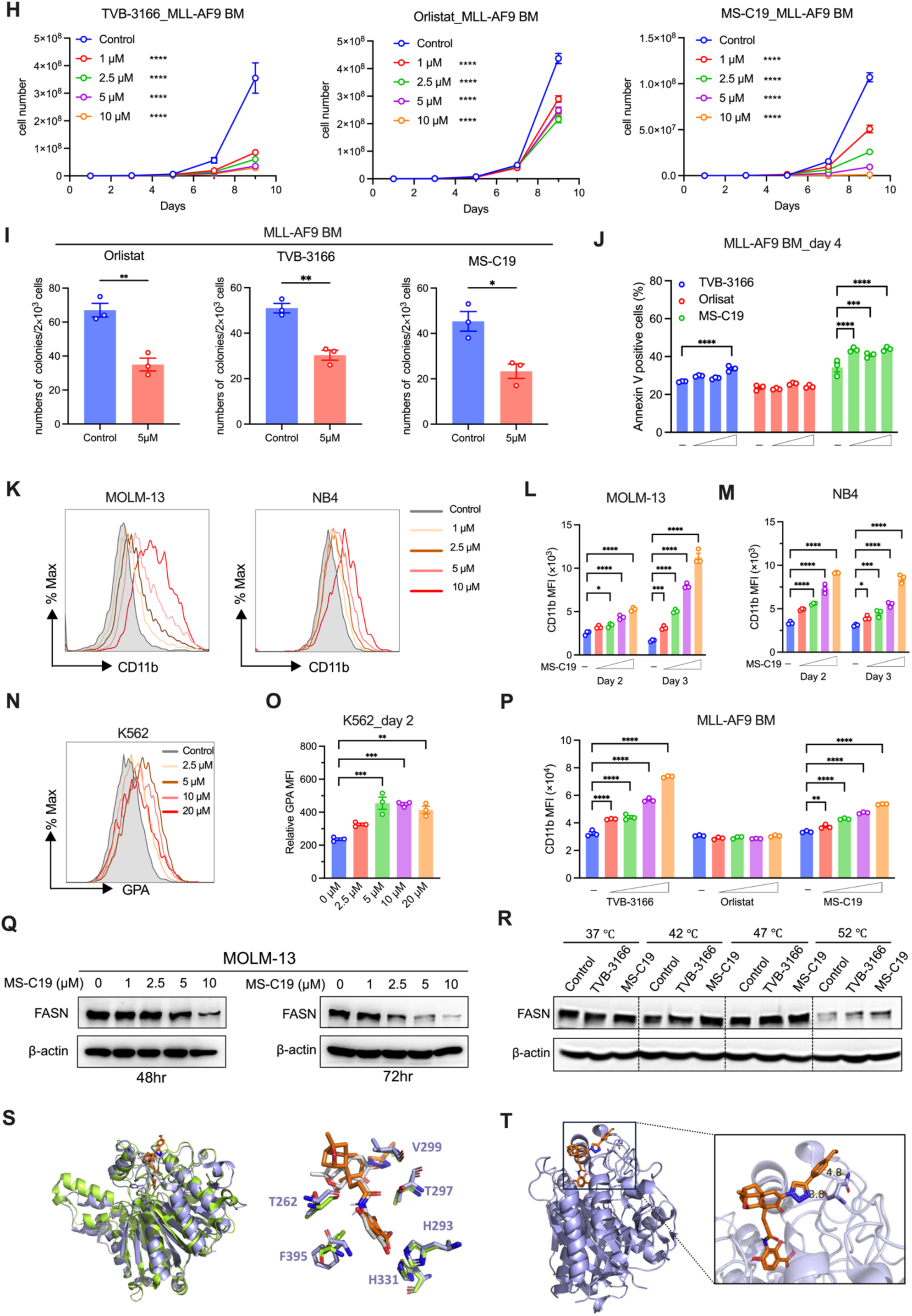
Synthesis and screening of a novel FASN inhibitor, MS-C19, from the PTM compound library. (**A**) The synthetic routines for a PTM derivatives library containing 70 compounds. (**B**) Primary screening of indicated compounds by treating K562 cells (20µM, 72hr) followed by measuring cell viability through mitochondrial MTT assay. (**C**) K562, MOLM-13, and NB4 cells were treated with varied concentrations of indicated FASN inhibitors for 72hr, the cell survival was determined by MTT assay, and IC50 values were determined for cytotoxicity of compounds. (**D-G**) Evaluations of apoptosis (D-E), cell cycle profiling (F), and colony formation (G) of leukemic cells treated with vehicle control or indicated FASN inhibitors. Representative flow cytometric plots for apoptotic assay in MOLM-13 challenged with MS-C19 for 48hr were shown in D. (**H**-**J**) Growth curve (H), colony formation (I), and apoptosis (J) of MLL-AF9-expressed BM cells treated with indicated FASN inhibitors. (**K**-**M**) Myeloid differentiation of MOLM-13 and NB4 treated by MS-C19 were assayed by CD11b staining. Representative flow cytometry histograms were overlayed and shown for day 3 in K. Mean fluorescence intensity (MFI) of CD11b was further quantitated in L and M. (**N**-**O**) Similar to H, except erythroid-specific differentiation marker GPA was assessed. (**P**) The MFI of CD11b in MLL-AF9 transduced BM cells treated with or without indicated FASN inhibitors were assayed by flow cytometric analysis. (**Q**) Western blotting analyses of FASN expression in MOLM-13 with or without MS-C19 treatment. β-actin, loading control. (**R**) Cellular thermal shift assay to examine the binding of TVB3166 or MS-C19 with endogenous FASN in MOLM-13 cells. The cells were treated with 10μM inhibitors at 37°C for 1 h followed by incubation at indicated temperature for 3 min. Western blotting analysis was then performed for FASN. (**S**) Structural alignment of ecFabF(C163Q)/PTM and FASN-KS domain/MS-C19. The model of FASN-KS domain/MS-C19 was generated by AutoDock Vina program. The structure of the FASN-KS domain is from PDB code: 3HHD. Conserved amino acid residues binding with MS-C19 in the FASN KS domain are labeled. (**T**) Predicted binding mode of additional ethylbenzene-triazole of MS-C19 in FASN-KS domain binding pocket. Experiments were conducted in biological triplicates, except quadruplicates in G. The statistical analysis was performed by two-way ANOVA in E, G, I, and J, and one-way ANOVA in L with Tukey’s corrections in all. *, P<0.05. **, P<0.01. ***, P<0.001. ****, P<0.0001

To determine whether MS-C19 directly binds to its potential target FASN, we employed a cellular thermal shift assay (CETSA). Expectedly, MS-C19 protected FASN from heat-induced protein denaturation or degradation (Fig.6R, and fig. S7C). Previously, the structure of ecFabF(C163Q) in complex with PTM has been revealed ^25^ (fig. S7D). The human FASN β-ketoacyl synthase (KS) domain exhibits high similarity to ecFabF, including the active pocket for PTM binding ^26^. We docked the PTM and MS-C19 into the active site. MS-C19 fits well with the KS domain in a similar orientation as PTM (Fig.5S, and fig. S7E). In particular, the carboxylates of the benzoic acids of both PTM and MS-C19 interact with the two His residues belonging to the His-His-Cys catalytic triad. Additionally, the ethylbenzene-triazole of MS-C19 extends further toward the surface of the binding pocket, where the ethylbenzene portion has the potential to form an amide-π stacking interaction with the side chain amide bond of FASN Asn 220 (Fig.5T). Moreover, the nitrogen atom of the triazole moiety may also form an additional hydrogen bond with the main chain carbonyl of Asn 220 (Fig.5T). These additional binding interactions between MS-C19 and FASN Asn 220 likely enhance the binding affinity of MS-C19 to FASN.

**Fig.6.**
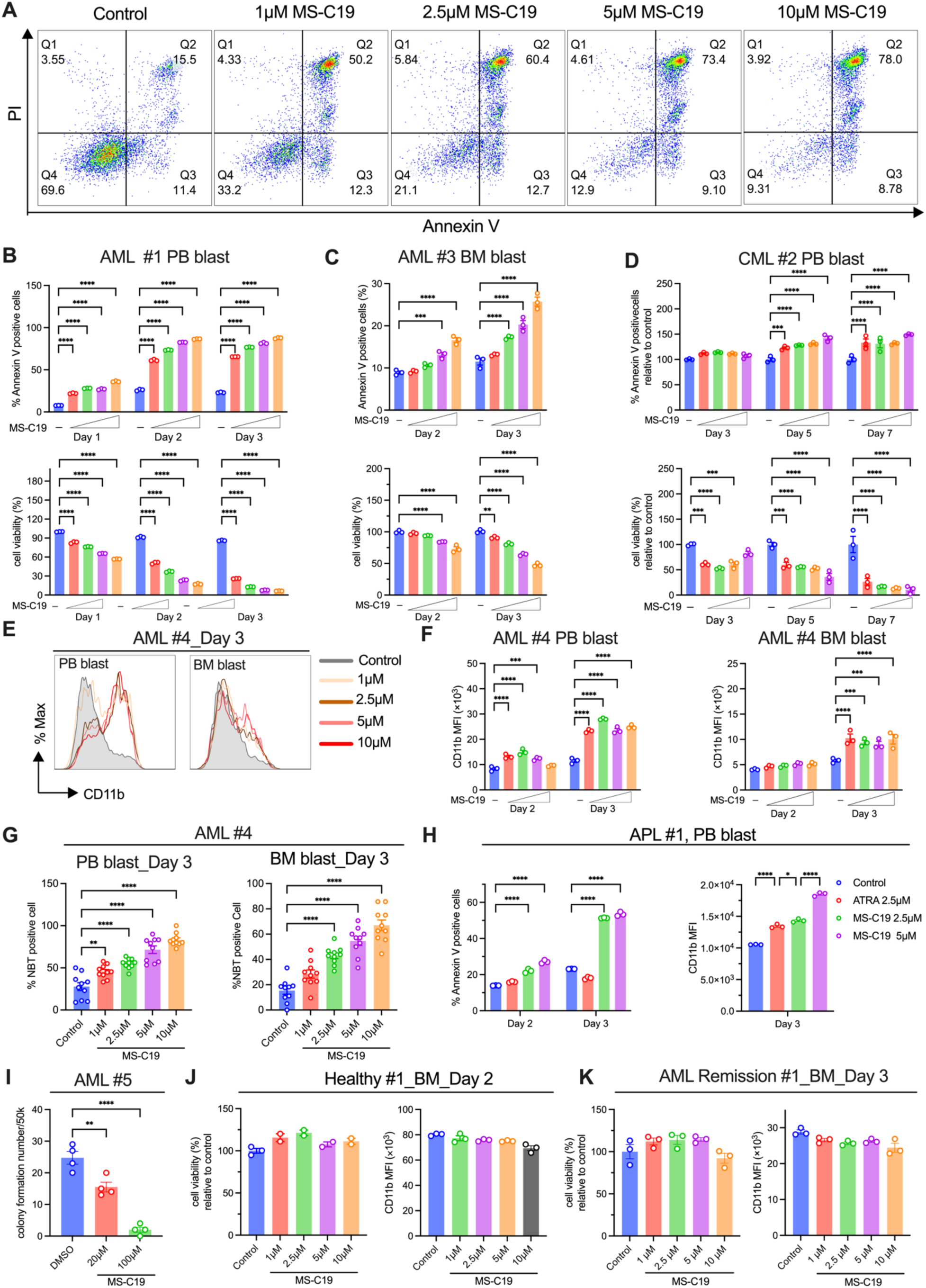
MS-C19 induces cell apoptosis and differentiation of leukemic blasts from clinical patients. (**A**-**B**) Peripheral leukemic blast cells from AML patients were treated with varied concentrations (1µM, 2.5µM, 5µM, 10µM) of MS-C19, cell apoptosis, and survival was examined by Annexin V staining and PI staining followed by flow cytometric assay. Representative dot plots on day 3 were shown in A. Quantification of Annexin V positive cells and cell viability was shown as the upper and lower panel in B respectively. (**C**) The same assay as in B, except bone marrow leukemic blast cells from AML patients were used. (**D**) Same as B except for peripheral leukemic blast cells from CML patients were tested with MS-C19 (2.5µM, 5µM, 10µM, 20µM). (**E**-**G**) Myeloid differentiation of either peripheral blasts or bone marrow blasts from AML patients was determined by CD11b expression or NBT staining. Representative flow cytometric histograms of CD11b staining were illustrated in E, and quantifications of mean fluorescence intensity of CD11b were shown in F. Portions of NBT positive cells were quantitated randomly from 10 fields under microscope and data was presented in G. (**H**) Apoptosis (left) and CD11b levels (right) determined in APL blasts treated with or without ATRA or MS-C19. (**I**) Assessment of colony formation ability of AML blasts. (**J-K**) Bone marrow mononuclear cells isolated from healthy donors (J) or AML patients under remission (K) were treated with or without MS-C19 at indicated concentrations for 48 or 72 hours, and cell viability and differentiation were determined by flow cytometric analysis. All experiments were conducted in triplicates. The statistical analysis was performed by two-way ANOVA in B, C, D, F, and H (left), and one-way ANOVA in G, H (right), and I with Tukey’s corrections for all. *, P<0.05. **, P<0.01. ***, P<0.001. ****, P<0.0001.

Collectively, the PTM derivative MS-C19 suppressed leukemic cell growth, survival, and clonogenicity, and promoted cell differentiation in vitro by targeting FASN. MS-C19 was superior to the commercial FASN inhibitors Orlistat and TVB-3166, demonstrating greater potency against leukemia *in vitro*.

### MS-C19 induced cell death and differentiation in clinical leukemic cells

We next decided to evaluate the efficacy of MS-C19 on human clinical samples. Leukemic blasts in PB or BM from primary AML and CML patients were obtained and cultured with vehicle control or MS-C19. We observed increased apoptotic cells and declined cell viability in MS-C19-treated leukemic blasts compared to vehicle-treated controls (Fig.6A-D, and fig. S8A-B). As expected, MS-C19 accelerated the myeloid differentiation of AML blasts (Fig. 6E-G, and fig. S8C-D). MS-C19 and Orlistat also induced robust apoptosis and triggered myeloid differentiation of clinical BM- and PB-derived AML cells (fig. S8E-G). In contrast to the ATRA-triggered differentiation in leukemic cell lines, ATRA failed to induce any cell apoptosis or myeloid differentiation in AML patient-derived leukemic blasts (fig. S8H-I), which is in line with the clinical observation that most favorable efficacy of ATRA treatment is for PML-RARA positive APL. MS-C19 also dampened the survival of APL blasts, and efficiently induced the differentiation of cells (Fig. 6H). More importantly, MS-C19 significantly reduced the clonogenicity of AML blast cells (Fig. 6I). By contrast, after being treated with MS-C19, no apparent cell death or differentiation were observed in the BM cells of individuals who were either healthy or in clinical remission (Fig. 6J-K). Overall, our data showed that targeting FASN by MS-C19 exerts anti-leukemia effects.

### FASN inhibition augmented lysosomal and inflammatory gene activation

To gain in-depth insights into the mechanisms underlying the role of FASN in leukemia, we performed bulk RNA-sequencing analysis of FASN-depleted or control MOLM-13 in addition to MS-C19 treated or untreated cells (fig. S9A-F). We identified 134 commonly upregulated genes in the presence of both FASN interference and MS-C19 treatment (Fig. 7A, upper panel). GO and KEGG analysis further clustered them into several signalosomes, in which the lysosomal activity and inflammation-related pathways appeared repeatedly (Fig. 7A-C). Accordingly, 56 commonly downregulated DGEs were revealed to be associated with mitotic cell cycle transition, nuclear division, and centromeric region of chromosome (Fig. 7D-E), which likely accounts for the decreased S phase in cells with either FASN depletion or MS-C19 insults (Fig. 1K and 5F). By quantitative PCR, we validated that *PLD3*, *LYZ*, *S100A8*, *HLA-DRA*, *GRN*, *TIMP1*, *CXCL8*, *TREM2*, *HMOX1*, and *CST3* were significantly induced, while *MTBP*, *TOP2A*, *FANCM*, *KIF14*, *SMC4* were profoundly compromised in FASN knockdown or MS-C19 treated cells (Fig. 7F and fig. S9G).

**Fig.7.**
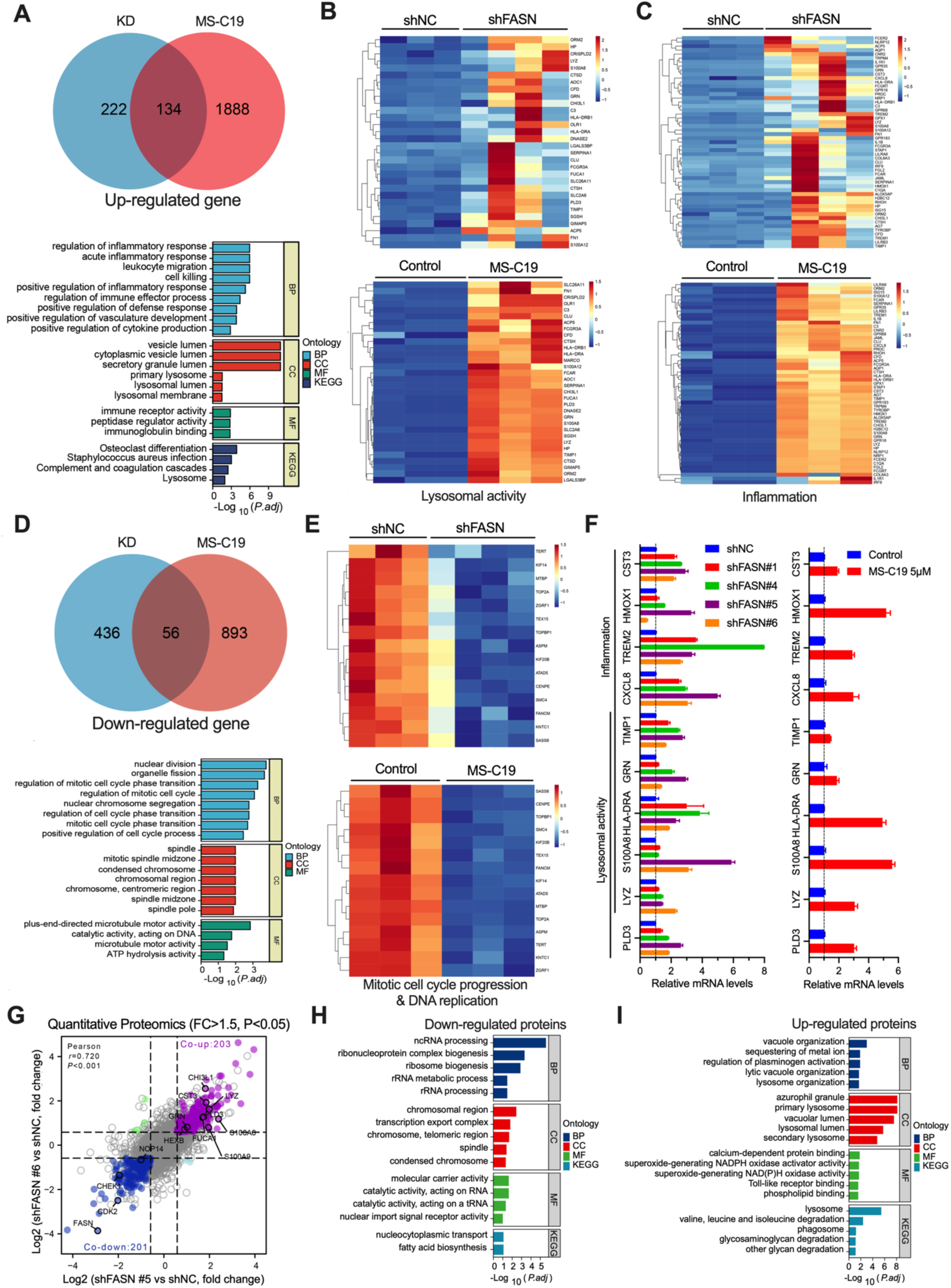
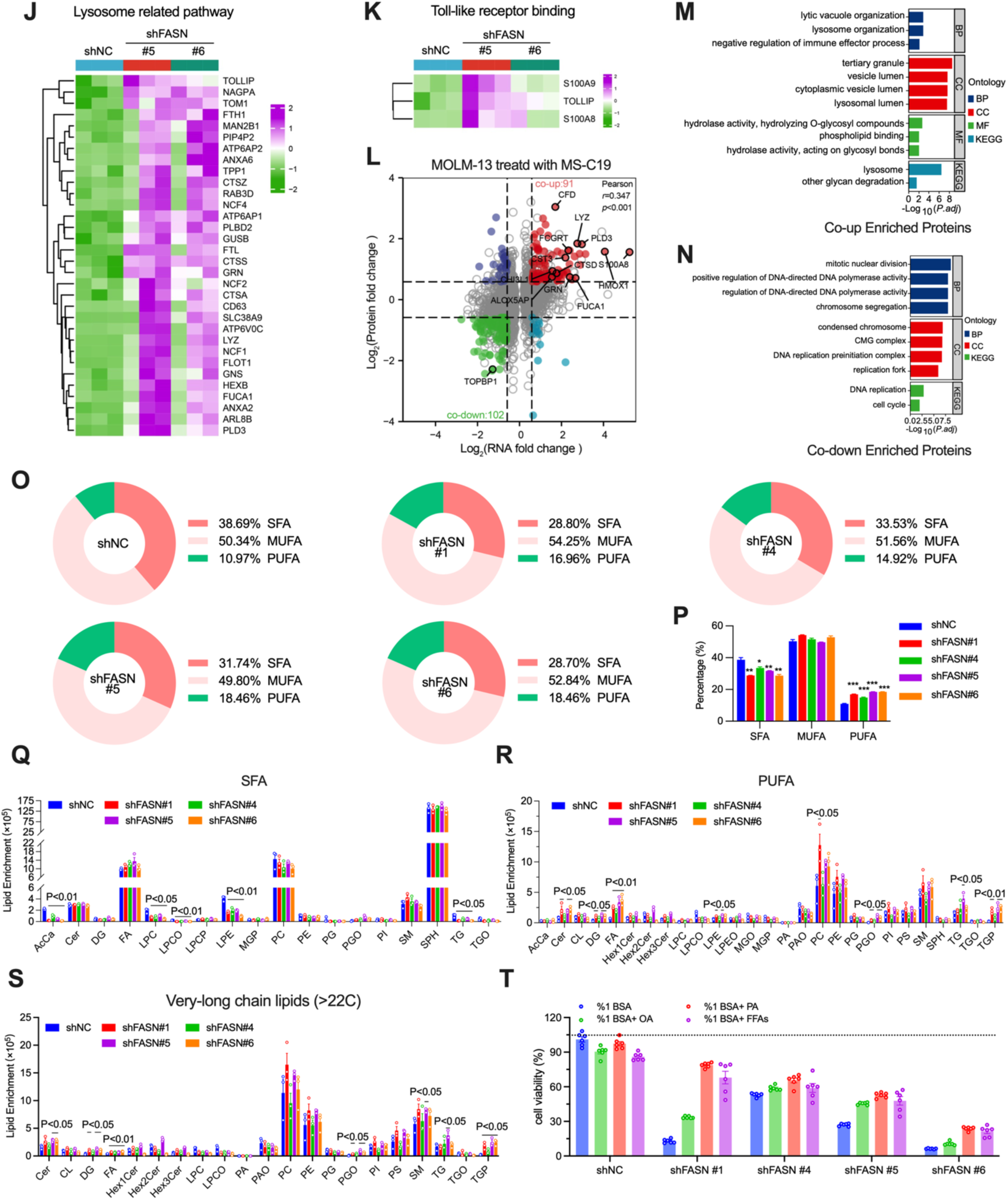
Multi-omics analysis reveals aberrant lysosomal gene activation and increased uptake of polyunsaturated fatty acids in FASN-depleted cells. (**A**) Venn plots showing the upregulated genes shared by both FASN depletion and MS-C19 treatment (upper). GO and KEGG pathway analysis of 134 overlapping genes were further shown (lower). (**B**-**C**) Deferentially expressed genes involved in the lysosomal activity and inflammation pathway were shown in hierarchically clustered heatmaps in B and C, respectively. (**D**-**E**) Similar to A and B, commonly down-regulated genes and mitotic spindle-related pathways were analyzed and illustrated. (**F**) Evaluation of indicated gene expression levels by qPCR in both FASN KD (left) and MS-C19 treated (right) cells. (**G**) Pearson correlation analysis of globally altered proteins in FASN KD cells determined by label-free quantitative proteomics. (**H-I**) Pathway enrichment analyses of differentially altered proteins identified in G. (**J**-**K**) Clustering heatmaps illustrating protein expression patterns in lysosome-related (J) and Toll-like receptor binding (K) pathways. (**L**) Integrated analysis of quantitative proteomics and transcriptomics in MOLM-13 cells untreated or treated with MS-C19 at 5µM for 48hr. Commonly up-regulated in red and down-regulated genes in green (fold change>1.5, p<0.05) were illustrated. (**M**-**N**) GO-term and KEGG pathway analysis of upregulated (M) and downregulated (N) genes identified in L. (**O**-**P**) Global lipidomic analyses of MOLM-13 cells depleted of FASN. Lipid distribution profiling was shown as pie charts in O, and was further quantitated in P. (**Q-S**) Determination of SFA (Q), PUFA (R), and very-long-chain lipids (S) in control or FASN KD cells. AcCa, Acylcarnitine; Cer, Ceramide; CL, Cardiolipins; DG, Diradylglycerols; FA, Fatty Acyls; Hex1Cer, Monohexosylceramide; Hex2Cer, Dihexosylceramide; Hex3Cer, Trihexosylceramide; LPC, Lysophosphatidylcholine; LPCO, Monoalkylglycerophosphocholines; LPCP, LPC-Plasmalogen; LPE, Lysophosphatidylethanolamine; MGP, Monopalmitin; PA, Phosphatidic acid; PAO, 1-alkyl-2-acylglycerophosphates; PC, Phosphatidylcholine; PE, Phosphatidyl ethanolamine; PG, Diacylglycerophosphoglycerols; PGO, 1-O-alkyl-2-acylglycerophosphoglycerols; PI, Phosphatidylinositol; PS, Phosphatidylserine; LPEO, Monoalkylglycerophosphoethanolamines; MGO, Monoalkylglycerols; SM, Sphingomyelin; SPH, Sphingosine; TG, Triradylglycerols; TGO, Alkyldiacylglycerols; TGP, Triglyceride Phosphate. (**T**) Cell viability of control or FASN KD cells treated with or without oleic acid (OA), palmitic acid (PA), or their mixture (FFA, OA: PA=1:2). The statistical analysis was performed using unpaired student t-test in P and Q between shFASN and shNC. *, P<0.05. **, P<0.01. ***, P<0.001. ****, P<0.0001.

By quantitative proteomics analysis, we revealed that while the declined proteins were associated with ribosome biogenesis, rRNA processing, and chromosome and spindles (Fig. 7H), the aberrantly upregulated proteins were mainly enriched in the lysosome and lytic vacuole organization-related pathways (Fig. 7I-J). FASN-depleted cells also exhibited augmented Toll-like receptor binding signaling with accumulated S100A8/S100A9 protein expression (Fig. 7K). We further examined the protein alterations triggered by MS-C19 and performed the integrative analysis with transcriptomic data (Fig. 7L). 91 co-upregulated proteins were enriched for lysosome or lytic vacuole organization (Fig. 7M), and 102 co-downregulated proteins were engaged in mitotic nuclear division, chromosome segregation, and DNA replication (Fig. 7N). These data suggest the essential role of FASN in cellular lysosomal and inflammatory gene expression and mitotic cell cycle progression.

### FASN maintains lipid homeostasis by modulating PUFAs and VLCFAs

By performing the global lipidomic analyses in FASN-deficient MOLM-13, we observed that the lipidic homeostasis was disrupted in FASN KD cells (Fig. 7O-P). Specifically, although the monounsaturated fatty acids (MUFA) remained unchanged (Fig. 7P, and fig. S10A), the saturated fatty acids (SFA) mostly provided by cellular FA biosynthesis were decreased in FASN KD cells (Fig. 7Q and fig. S10B-C). On the contrary, the polyunsaturated fatty acids (PUFAs), mainly from food uptake, were significantly enriched upon FASN loss (Fig. 7R and fig. S10D-E), suggesting a complementary usage of exogenous lipids in FASN-depleted cells. In addition, the very-long-chain fatty acids containing lipids (VLCFAs) (i.e., DG, FA, PGO, TGP) were increasingly synthesized (Fig. 7S). An exogenous supply of either oleic acid (OA), a MUFA, palmitic acid (PA), an SFA, or free fatty acid (FFA, OA: PA=1:2) significantly improved the survival of FASN-depleted cells (Fig. 7T). However, they did not fully revert the FASN-loss-induced cell death, suggesting that FASN regulates multiple signaling pathways in addition to lipid homeostasis.

### FASN deficiency leads to lysosome-dependent cell death (LDCD) other than lysosome biogenesis

Reduction of FASN expression by ATRA stimulation was previously reported to augment lysosome biogenesis through enhanced TFEB nuclear translocation triggered by mTOR inactivation ^17,27^. We observed that lysosomal genes were actively transcribed upon FASN loss with TFEB signature enriched (fig. S11A-D). Importantly, by treating the cells with bafilomycin A1, a lysosomal acidification inhibitor, we observed a significant reversal of apoptosis under both FASN depletion and MS-C19 treated conditions (Fig. 8A-B). However, when the cellular lysosomes were examined by transmission electron microscope (TEM) scanning and lysosomal associated membrane protein 1 (LAMP1) staining, unexpectedly, both the lysosome numbers and LAMP1 fluorescence intensity remained unchanged upon FASN loss (Fig. 8C-D, and fig. S12A-B). Since Fasn KO adipocytes exhibited impaired lysosome maturation and membrane dynamics^28^, lowering FASN in leukemic cells may destroy lysosomal membrane permeabilization (LMP), which in turn results in the translocation or leakage of intra-lysosomal components such as cathepsins^29^, to the cytoplasm to induce lysosomal-dependent cell death (LDCD). In support of this hypothesis, Galectin-3 puncta formation, a marker of LMP ^30^, was accumulated in FASN-depleted MOLM-13 cells (Fig. 8E and fig. S12C). We thus conclude that FASN inactivation promoted TFEB-governed lysosomal gene transcription but contributed barely to lysosome biogenesis. FASN safeguarded the permeabilization of the lysosomal membrane, and loss of FASN led to compromised LMP and subsequent LDCD.

**Fig.8.**
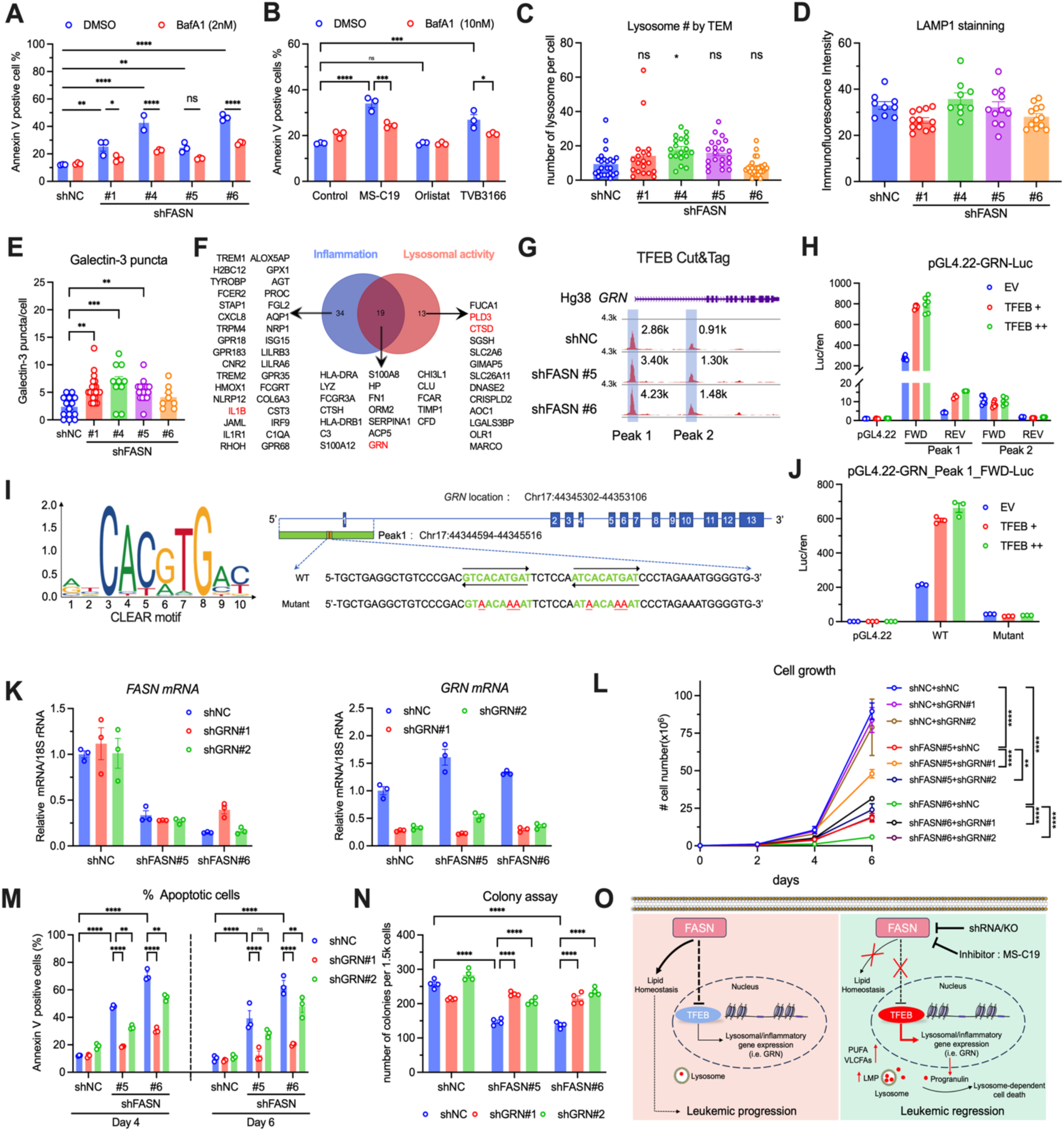
FASN inactivation-induced leukemic cell death relies on GRN expression. **(A-B)** MOLM-13 with FASN depletion (A) or inhibition (B) was treated with a lysosome inhibitor, Bafilomycin A1 (BafA1), for 48hr. Percentages of apoptotic cells were determined by annexin V staining. The cells in B were pretreated with FASN inhibitors for 12 hours prior to BafA1 treatments. (**C**) Quantifications of lysosome numbers visualized by SEM. (**D**) Immunofluorescent intensity quantifications of LAMP1 staining. (**E**) Quantification of Galectin-3 puncta per cell in the indicated conditions determined by fluorescence microscopy. (**F**) Venn diagram showing overlapping genes from indicated pathways. (**G**) Genome browser tracks showing TFEB occupancy at the *GRN* locus in control and FASN-depleted cells from CUT&Tag. (**H**) Luciferase activity in *GRN*-luciferase reporter constructs induced by ectopic TFEB expression performed in 293T cells. (**I**) Schematic illustrating the consensus CLEAR motif recognized by TFEB (left), and the potential motifs identified in GRN promoter region (right). The wild-type sequence (WT) is annotated in green, while the mutated nucleotides (Mutant) are indicated in red. (**J**) Similar luciferase activity assays as in H were performed using 293T cells co-transfected with either the empty vector or TFEB plus luciferase reporters containing either intact (WT) or mutated (Mutant) *GRN*-luciferase. (**K**) *FASN* (right) and *GRN* (left) mRNA levels in MOLM-13 cells infected with control or FASN shRNAs plus with or without GRN depletion as determined by quantitative PCR. (**L**-**N**) The growth curve (L), apoptotic cell rates (M), and clonogenicity (N) of indicated cells in K were determined. (**O**) Schematic graphs depicting the mechanism of FASN inactivation-induced leukemic regression. Inhibition or loss of FASN not only triggers the TFEB transcriptional activation to enhance the lysosomal gene such as *GRN* expression, but also disrupts the lipid homeostasis, leading to augmented PUFA uptake, and VLCFAs synthesis. FASN deficiency compromises the lysosomal membrane permeabilization (LMP) and promotes the lysosome-dependent cell death (LDCD) that relies on Progranulin. Two-way ANOVA in A, B, L, M, and N, one-way ANOVA in E with Tukey’s corrections in all. *, P<0.05. **, P<0.01. ***, P<0.001. ****, P<0.0001.

### Progranulin depletion reverts FASN inactivation-induced cell death

To search for a bona fide downstream target that mediates FASN depletion- or inhibition-induced cell death, we focused on the co-upregulated inflammatory response and lysosomal genes (Fig. 8F). Given that lower expression of FASN predicts a favorable survival outcome in de novo AML patients (Fig. 1C), we hypothesized that the upregulated genes upon FASN depletion or inhibition should prolong the patient survival if they are functional targets of FASN deficiency. By interrogating the event-free survival of untreated AML patients associated with the genes identified in pathways of inflammation and lysosome (fig. S13A-C), we uncovered that over 50% (37/66) genes tended to improve patient survival when highly expressed (fig. S13D). More specifically, *GRN*, *TIMP1*, *HLA-DRA*, *CFD*, and *SERPINA1* were positively associated with patient survival significantly (P<0.05) (fig. S13D).

*GRN* encodes progranulin, a glycoprotein located in cellular lysosome compartments. Emerging evidence has implicated the essential role of progranulin in neuroinflammatory responses and lysosomal functionality involved in neurodegenerative diseases ^31–33^, yet its role in leukemic progression remains unclear. The expression of GRN is suggested to be governed by both TFEB-dependent and independent mechanisms ^27,34,35^. We first sought to explore whether TFEB directs the transcriptional activation of *GRN* in the context of FASN depletion. By CUT&Tag sequencing, we revealed that the TFEB binding intensity near the transcription start sites (TSS) was broadly augmented when FASN was depleted (fig. S14A). This enhancement of TFEB binding could either stimulate (i.e., *CTSD*, *HMOX1*, *TREM1*) or suppress (i.e., *TOP2A*, *MTBP*, *KLF14*) the gene expression (fig. S14B-E). The predicted top motifs enriched upon FASN KD were for TFE3 and TFEC (fig. S14F), all the homologs of TFEB. Most importantly, we found TFEB bound to *GRN* genomic loci upon FASN ablation within two significant binding peaks (Fig. 8G). Luciferase assay demonstrated that potent TFEB-responsive cis-elements only existed in the promoter region (Peak1) of the *GRN* gene (Fig. 8H). Mutation of the conserved CLEAR motif (CAYRTG) recognized by TFEB largely abolished the luciferase activity that was induced by TFEB over-expression (Fig. 8I-J), suggesting that *GRN* served as a *bona fide* downstream target of TFEB in leukemic cells when FASN was depleted.

We next asked whether anti-leukemic effects exerted by FASN loss relied on GRN’s upregulation. To this aim, *GRN* was depleted further by shRNA in the presence of FASN ablation (Fig. 8K). Significantly, the loss of progranulin largely reverted the reduced cell growth, increased cell apoptosis, and declined clonogenicity of MOLM-13 cells lacking FASN (Fig. 8L-N). More importantly, although protein-protein interaction networks revealed IL-1B as the central node, blocking of IL-1B by shRNA had minimal rescue effects on cell growth, cell survival, and clonogenicity in FASN-depleted cells (fig. S15A-E). Similarly, depletion of either CTSD, the major lysosomal endopeptidase, or PLD3 that hydrolyzes phospholipids in lysosomes could not revert the anti-leukemic effect of FASN deficiency (fig. S16A-F). Collectively, our results highlight the specificity of Progranulin in mediating the FASN depletion-induced cell death and support that FASN-deficiency triggered activation of TFEB-GRN axis suppressed leukemia.

## Discussion

The most important finding of our work is that genetic ablation of FASN at either early or late stages of leukemia can effectively and significantly suppress the leukemic progression. We efficiently delete Fasn in *Fasn^fl/fl^* Mx1-Cre mice by polyIC administration before MLL-AF9 retroviral transduction, and pre-ablation of Fasn not only blocked the oncogene-induced cell proliferation and transformation *in vitro* but also ameliorates leukemic mortality in transplants *in vivo*. In addition, by the manners of shRNA knockdown, CRISPR-Cas9 knockout in leukemic xenografts, or conditional *Fasn* KO after MLL-AF9 viral infection and transplantation, the absence of FASN (Fasn) protein prevented the disease progression and improved the animal survival (Fig. 3). Other investigation also provides elegant evidence for supporting the FASN dependency in oncogenic activation of KRAS, HER2, AKT, and loss of PTEN-induced tumorigenesis ^8,36,37^. Most importantly, similar to the observation that the haploid insufficiency of Fasn caused much fewer of *Fasn*^+/-^ mutant progeny than Mendelian inheritance prediction ^38^, we found leukemic progression driven by MLL-AF9 also relied on *Fasn* gene dosage, as *Fasn^fl/+^* Mx1-Cre donor cells, to a less extent when comparing with *Fasn^fl/fl^* Mx1-Cre cells, significantly impeded the leukemogenesis, and prolonged the mice survival (Fig. 3K). Thus, FASN could be a key node in oncogene-induced tumorigenesis.

FASN is essential for several physiological activities, including embryogenesis ^38^, neurogenesis ^39,40^, inflammation ^41,42^, autophagy ^28^, and energy homeostasis ^43^. However, the role of FASN in hematopoiesis has not been fully understood. Previously, tamoxifen-induced global knockout of *Fasn* in mice led to a disrupted granulopoiesis, and mice exhibited leukopenia in a cell-autonomous manner ^44^. One significant finding from our study is that hematopoietic specific deletion of *Fasn* minimally affected the normal murine hematopoiesis. *Fasn^-/-^* Mx1-Cre or *Fasn^+/-^* Mx1-Cre mice had comparable blood cell counts with control mice after polyIC treatment (Fig. 4A). Similarly, Fasn-deficient hematopoietic stem progenitor cells remained either unchanged in numbers or intact in the aspect of engraftment ability (Fig. 4F-Q). Our results thus far highlight the rationality of prospectively targeting FASN in hematopoietic disorders. Our data also reflect that global deletion of Fasn instead of tissue-specific ablation may cause extra insults from microenvironments to HSCs that confer irreversible damage to its self-renewal and differentiation.

The fatty acid biosynthesis, which depends on several core enzymes such as FASN, stearoyl-CoA desaturase (SCD), very-long-chain fatty acids elongase 6 (ELOVL6), and acetyl-CoA carboxylase 1 (ACC1), has been recently highlighted as a potential vulnerability for leukemia therapy ^2–4,17,18,20^. For instance, the sterol regulatory-element binding proteins (SREBP)-regulated lipogenic response was revealed to protect leukemic stem cells (LSCs) from intracellular nicotinamide adenine dinucleotide (NAD) depletion-induced apoptosis ^2^. Free fatty acid derived from triglyceride hydrolysis via lipophagy supports oxidative phosphorylation (OxPHOS) in AML ^4^, and upregulated fatty acid oxidation underlined the venetoclax/azacytidine resistance in acute myeloid leukemia ^20^. Genetic disruption of *Elovl6* blocked acute myeloid leukemia (AML) development in the MLL-AF9 mouse model ^18^. More recently, activation of C/EBPα and Fms-like tyrosine kinase 3 (FLT3) increased lipid anabolism primarily through SCD, and inhibition of SCD sensitized the FLT3-mutant AML cells to ferroptosis ^3^. Additionally, silencing FASN sensitizes the myeloid leukemia cells to ATRA-triggered myeloid differentiation *in vitro* ^17^. Our research provides direct evidence that inactivation of FASN *in vivo* significantly hinders the leukemic progression in AML. Noteworthy, while we and others demonstrate that antagonizing FA biosynthesis through inhibition of FASN or SCD confers therapeutic potential in leukemia, previous work demonstrated that stabilization of ACC1, a rate-limiting enzyme for fatty acid synthesis, suppressed the Trib1-COP1 complex- or MLL-AF9-driven leukemogenesis *in vivo* ^21^. These conflict findings likely reflect the diverse and non-redundant roles of different lipid metabolic enzymes in the pathogenesis of hematologic cancer with genetic variabilities and further underscore the need for caution when targeting lipid metabolism in cancer via inhibiting specific metabolic enzymes.

PTM, initially identified as a bacterial β-ketoacyl-(acyl-carrier-protein (ACP)) synthase I/II (FabF/B) inhibitor in fatty acid synthesis ^45^, was later shown to suppress mammalian FASN activity with therapeutic potential for metabolic disorders ^23^. Although FASN inhibitors are extensively studied in oncology ^5,11,12^, the limited intrinsic antitumor efficacy of PTM hindered its exploration in cancer biology ^45^. By synthetic modification, we generated PTM-derived compound libraries. PTM-6p previously exhibited antitumor activity against non-small cell lung cancer and melanoma by targeting FASN’s ketosynthase (KS) domain ^6^. Here, we identified MS-C19, another PTM derivative, as a potent inducer of leukemic cell death and differentiation, significantly reducing the tumorigenicity of leukemic cells. As expected, MS-C19 bound to the KS domain of FASN more tightly than PTM (Fig. 5T). To our knowledge, this is the first PTM-derived compound that exerts anti-leukemia effects. Whether MS-C19 has broader efficacy against other solid cancers and hematologic malignancies warrants further investigation.

FASN contains multiple catalytic domains, and dozens of FASN inhibitors have been identified for targeting diverse domains to date ^5,46–48^ (supplementary table S4). Interestingly, different FASN inhibitors exhibit differential psychopharmacological effects and cellular response, e.g., in macrophages, cerulenin or C75 that inhibits the KS domain, but not GSK2194069 or Orlistat targeting β-ketoreductase (KR) or thioesterase (TE) domain, prevented the inflammation triggered by LPS ^41^. Accordingly, inhibition of FASN by C75 blocked TLR signaling, improved neutrophil chemotaxis, and increased the survival of mice under septic shock, whereas GSK2194069 had no such effects^42^. We also demonstrated that MS-C19 had potent antileukemia activity, while TVB-3166 and Orlistat had either mild or limited effects. These inhibitory discrepancies suggest that different domains of FASN carry out diverse roles to ensure the multi-faced biological functions of FASN.

Previous work suggested that compromised FASN protein expression instead of inhibiting the enzymatic activity contributed to the differentiation-promoting therapy in leukemia cells *in vitro* ^17^. Consistently, in MOLM-13 and NB4 cells, we also observed that *in vitro* depletion of FASN by shRNA enhanced the CD11b expression upon ATRA treatment. However, in contrast to that, Orlistat treatment had no effect on AML differentiation in previous work and in this study, the compound MS-C19 boosted the myeloid differentiation of both AML cell lines and clinical AML blasts. This observation is likely attributed to the decline of FASN protein levels upon MS-C19 treatment (Fig. 5Q), which has also been verified for PTM derivatives in other cell types ^6,49^.

Increased LMP indicates the destabilization or loss of lysosomal membrane integrity ^29,50^. In our study, we observed that the marker of LMP, Galectin-3 puncta formation, was significantly increased in FASN-depleted cells (Fig. 8E). This suggests that the absence of FASN could impair lysosomal membrane integrity, likely due to its critical role in lipid biosynthesis. Indeed, lipidomic analysis of whole cells revealed an imbalance in cellular lipid content following FASN depletion, with significant increases in VLCFAs and PUFAs (Fig.7R-S). VLCFAs are mainly metabolized by β-oxidation in peroxisomes, where over 30% of hydrogen peroxide in mammalian tissues is generated ^51^. Increased PUFAs in the cell membrane are also toxic, as they form complexes with reactive oxygen species (Lipid-ROS) to initiate lipid peroxidation, leading to subsequent cell death like ferroptosis ^3^. One plausible mechanism by which FASN depletion affects lysosomal integrity is through ROS or Lipid-ROS-triggered lipid modification or peroxidation, which compromise or disrupt the lysosomal membranes (Fig.8O). On the other hand, given that lysosomes serve as the central trafficking hub for lipids and their membranes are primarily composed of lipids, including cholesterol, phospholipids, and sphingolipids, FASN depletion could disrupt the balance of these essential components, leading to impaired lysosomal membrane integrity or function ^50,52^. This hypothesis is supported by studies demonstrating that lysosomal phosphatidylserine/cholesterol and cytosolic sphingomyelin are necessary for the rapid repair of LMP ^53–55^, and loss of Fasn decreases certain membrane phosphoinositides essential for lysosome maturation ^28^. Future studies should explore the specific lipid species modulated by FASN that contribute to lysosomal stability.

*GRN* heterozygous or null mutation was revealed to cause frontotemporal lobar degeneration (FTLD) ^56,57^, and the premature stop of *GRN* by a homozygous variant resulted in a lysosomal storage disease, neuronal ceroid lipofuscinosis (NCL) ^58^. Compromised expression or dysregulation of progranulin contributed to neurodegenerative diseases by promoting inflammation and lysosomal dysfunction ^59,60^. In this study, we found *GRN* expression was aberrantly upregulated by FASN knockdown through the TFEB-orchestrated transcriptional network, and suppressing *GRN* gene activation could significantly revert the FASN-deficiency caused leukemic remission phenotype. Moreover, *GRN* expression was positively associated with AML patient overall survival. Despite the underlying mechanisms of *GRN* in oncology remaining elusive, our work emphasized the role of *GRN* in antagonizing the exacerbating effect of FASN on leukemogenesis.

In summary, our studies demonstrated the critical roles of FASN in leukemogenesis *in vitro* and *in vivo*. We identified a potent FASN inhibitor, MS-C19, and pharmaceutical targeting FASN exerted anti-leukemic effects. Our work highlighted that, in addition to the dysregulated lipid homeostasis, the FASN-deficiency-induced *GRN* expression determines the lysosome-dependent leukemic cell death (Fig. 8O). These findings will significantly benefit the anti-leukemia therapy and may also have far-reaching therapeutic implications for modulating the disease progressions of varied cancers through the inventions of FASN-governed lipogenesis and gene expression networks.

## MATERIALS AND METHODS

All the materials and methods are in the Supplementary Materials.

## Supporting information

Supplementary files including Materials and Methods, Fig. S1-S16.

## Authorship Contributions

Conceptualization, M.S., L. Z., Y. H., Y.S., and Y.M.; Methodology, M.S., Z. L., Y.L., X.X., J. O., A. Z., H. L., Q. L., X. M., Y. Y., K. L., Y. H., S. X., J. L., L. C., X. L., J. Y., C. W., L. L., L. Z., Y. S., Y. H., and Y.M.; Investigation, M.S., Z. L., Y.L., X.X., J. O., A. Z., H. L., Q. L., Y. H., L. Z., Y. S., Y. H., and Y.M.; Writing-Original Draft, Y.M.; Writing-Review & Editing, Y.H. and Y.M.; Funding Acquisition, M.S., L. Z., Y.H. and Y.M.; Supervision, Y.M.

## Data and materials availability

All data associated with this study are present in the paper or the Supplementary Materials. Available experimental resources and mouse strains generated through this study are available from Y.M. under a material transfer agreement with Hunan University upon request.

## Supplementary Materials

The PDF file includes:

Materials and Methods

Figs. S1 to S16

Tables S1 and S4

