## Supplementary files including Materials and Methods, Fig. S1-S16. for "FASN Inactivation-Induced Progranulin (GRN) Expression Promotes Lysosome-Dependent Cell Death to Suppress Leukemogenesis"

Meng Su et al.

**The PDF file includes:**

Materials and Methods

Figs. S1 to S16

Tables S1 and S4

### Materials and Methods

#### Mice

*Fasn* floxed mice and Mx1-Cre mice in C57BL/6J (CD45.2) background were purchased from Cyagen Biosciences (Suzhou, China). Immunodeficient NSGS mice were purchased from Jackson Laboratory. The CD45.1-positive congenic mice and immunodeficient NKG mice were purchased from Cyagen (Suzhou, China). Both male and female mice (8~12 weeks old) were used and randomly assigned for all experiments unless indicated elsewhere. The genotyping primer sequences were listed in the supplemental table 1. All animal experiments were conducted following the guidelines for the care and use of laboratory animals and were approved by the Institutional Animal Care and Use Committee (IACUC) at Hunan University (HNU-IACUC-2021-106).

#### Primary leukemia patient samples

AML patient samples were collected from peripheral blood or bone marrow aspirations with informed consent, and mononuclear cells (MNCs) were isolated. All experiments involving human samples were conducted in compliance with relevant ethical regulations and were approved by the ethics committees of medical research at Hunan University (HNU-SMYXLL-AP-660001). Primary AML patient-derived mononuclear cells were cultured as described previously (*1, 2*). Briefly, cells were cultured in IMDM (Gibco, Cat#C12440500BT) supplemented with 20% FBS (NEWZERUM, Cat#SE141-500), 10 ng/mL recombinant human cytokines IL3 (Stemimmune LLC., Cat #HCT-13-0010), IL-6 (Stemimmune LLC., Cat #HIM-16-0010), FLT3L (Stemimmune LLC., Cat #HHM-FT-0010), SCF (STEMCELL Technologies, Cat#78062), and 1% penicillin/streptomycin (Gibco, Cat#15140122). Leukemic patient characteristics are detailed in Supplementary Table S2.

#### Generation of PTM series compound library

The critical step for the construction of the platensimycin triazole library is the semi-synthesis of 6-azidyl substituted platensimycin **2a** from platensimycin epoxide. We obtained compound **2a** with a yield of 87%, by opening the strained epoxide ring with $NaN_3$, followed by ammonium chloride-mediated dehydration (fig. S5B). Next, the 70-membered platensimycin triazole library was prepared via click chemistry through two different routes. A typical copper(I)-catalyzed azide alkyne cycloaddition (CuAAC) reaction condition (fig. S5C, route a) was initially applied, resulting in 20 platensimycin triazoles with excellent yields as judged by LC-MS analysis, while the remaining ones had limited yields. Alternatively, phenylacetylene was used as the model terminal-alkyne substrate to improve the efficiency of the click reaction for generating **2a** (supplemental table S1). We found the yields of triazole **17** were neither improved by raising the reaction temperature to 80℃, changing the solvent to DMSO or DMF, nor running the reaction under nitrogen protection (supplemental table S1, entry 1-5). Satisfactorily, the yield of **17** was eventually improved to 95% in the presence of one equivalent of copper chelating ligand TBTA (tris(benzyltriazolemethyl)amine) regardless of nitrogen protection (supplemental table S1, entry 6-7). Similarly, the remaining fifty platensimycin triazoles were synthesized with high yields in the presence of TBTA (fig. S5C, route b).

**General experimental procedure.** The commercial reagents were used as received. All $^{1}H$ and $^{13}C$ NMR spectra were recorded on a Brucker 600, 500, or 400 MHz spectrometer. Chemical shifts were reported in ppm relative to the internal standard tetramethylsilane ($\delta$ = 0 ppm) for $^{1}H$ NMR and deuterio-chloroform ($\delta$ = 77.00 ppm) for $^{13}C$ NMR spectroscopy. The following abbreviations were used to designate chemical shift multiplicities: s = singlet, d = doublet, t = triplet, q = quartet, m = multiplet, and br = broad. High-resolution mass spectra (HRMS) were recorded on an LTQ-ORBITRAP-ETD instrument. Samples were analyzed on a Waters e2695 high-performance liquid chromatography (HPLC) system equipped with a PDA detector and a Waters Sunfire C18 column (150 × 4.6 mm). The mobile phase consisting of buffer A (ultrapure $H_2O$ containing 0.1% HCOOH and 0.1% $CH_3CN$) and buffer B (chromatographic grade $CH_3CN$ containing 0.1% HCOOH) was applied at a flow rate of 1 mL/min. All compounds used for their MIC determination and cell viability possessed a purity of at least 95%, based on HPLC and $^1$H NMR analysis.

**Synthesis of 2a**. To a solution of platensimycin epoxide in EtOH/$H_2O$ (8.8 mL/ 0.88 mL) was added $NaN_3$ (866 mg, 0.66 mmol, and $NH_4Cl$ (70 mg, 1.32 mmol). The reaction mixture was stirred under reflux conditions and 80 °C for 12.0 h. Then, the reaction was concentrated under a vacuum and neutralized with 2 M HCl (2.0 mL). The residue was dissolved in EtOAc and washed with brine and fresh water. The organic phase was dried and concentrated; the generated crude products were subjected to column chromatography (eluent: PE / EA / AcOH = 100：20：0.25 to PE / EA / AcOH = 50：50：0.25) to provide **2a** (157.0 mg, yield 87%).

**Compound library synthesis.** Compound **2a** (70 mg, 145.25 μmol) was dissolved in 3.75 mL DMF, which then was transferred to 70 vials with 50 μL of **2a** solution. Then 50 μL TBTA (81.75 mg, 145.25 μmol in 3.75 mL DMF) was similarly added to the above reaction vials. The 70 alkyne substrates (311.2 μmol) were next dissolved in DMF (5 mL), and 50 μL of each alkyne was also added. Finally, $CuSO_4$ (10.38 μmol, 20 μL), sodium ascorbate (51.88 μmol, 110 μL), and deionized $H_2O$ (50 μL) were added. The whole reaction mixtures were stirred for 12 h at room temperature, and the solvents were removed under vacuum. Appropriate amounts of DMSO were added to the crude reaction mixture to make the concentration of final products to ~5 mg/mL.

**General procedure for synthesis of CuAAC reaction products (a).** To a solution of **2a** in tertiary butanol/$H_2O$ (5.0 mL/ 5.0 mL) were added the alkyne substrates (228.25 μmol), $CuSO_4$ (1 mg, 4.15 μmol), sodium ascorbate (4.1 mg,

20.5 μmol). The mixture was stirred at room temperature for 12 h. Then, the reaction mixture was concentrated under a vacuum. The resulting residue was next dissolved in EtOAc and washed with brine and fresh water. The organic phase was dried and concentrated; the generated crude products were subjected to column chromatography (eluent: PE / EA / AcOH = 100: 20: 0.25) to provide a compound (yield＞95%).

**General procedure for synthesis of CuAAC reaction products (b).** To a solution of **2a** in DMF/$H_2O$ (5.0 mL/ 5.0 mL) were added the alkyne substrates (228.25 μmol), TBTA (109 mg, 207.5 μmol), $CuSO_4$ (200 μL,100 mg/mL$H_2O$, sodium ascorbate (1100 μL,100 mg/mL $H_2O$). The mixture was stirred at room temperature for 12 h. Then, the reaction was concentrated under a vacuum, and the residue was dissolved in EtOAc and washed with brine and fresh water. The organic phase was dried and concentrated; the generated crude products were subjected to column chromatography (eluent: PE / EA / AcOH = 100：20：0.25) to provide products (yield＞95%).

### Chemicals

TVB-3166 (MCE, Cat#HY-120394), Orlistat (MCE, Cat# HY-B0218), Bafilomycin A1 (GLPBIO, Cat#GC17597). All the inhibitors were dissolved in dimethyl sulfoxide (DMSO), which also served as vehicle control in all treatments unless otherwise mentioned.

### Cell culture

HEK293T cell line was cultured in DMEM (Gibco, Cat#C11995500BT) supplied with 10% FBS. Human leukemic cell lines MOLM-13 and Kasumi-1 were cultured in RPMI-1640 medium (Gibco, Cat#C11875500BT) with 20% FBS, and K562 and NB-4 were maintained in RPMI-1640 medium containing 10% FBS. Penicillin/Streptomycin (1%) was provided for all the cell cultures. Cell numbers were quantitated in biological triplicates using a hemocytometer to decipher the cell growth curve.

**Plasmids and virus production**

pLV3-CMV-FASN (human)-3×HA-CopGFP-Puro plasmid was purchased from Miaoling Plasmid Platform (P47948, Wuhan, China) and was verified by DNA sequencing. Short hairpin RNAs (shRNA) targeting the FASN CDS region were designed by BLOCK-iT™ RNAi Designer (Thermo Fisher Scientific), and single-guide RNA (sgRNA) against FASN was designed by Benchling (https://www.benchling.com/crispr). The oligos were synthesized and inserted into the pLKO.1-GFP-Puro vector or pLenti-CRISPRv2 by Ligation high V2 (TOYOBO, Cat#2317062). All the target sequences for shRNA and sgRNA were listed in the supplementary table S3. Lentivirus was generated in HEK293T cells by co-transfecting target constructs with lentiviral packaging plasmids pMDL, REV, and VSVG according to the previous protocol using polyethyleneimine (PEI) (Catalog #23966, Polysciences) (*3*). MSCV-MLL-AF9-IRES-YFP retroviral construct was kindly provided by Prof. Zhang, Haojian (Wuhan University). The generation of retroviruses was performed by co-transfection of pCL-Eco packaging plasmid with MLL-AF9 construct into HEK293T cells. Viral supernatants were harvested at 48 hours after transfection.

**Lentiviral infection, selection, and generation of singe-cell derived CRISPR-Cas9 FASN KO clones**

$2\times10^5$ MOLM-13, K562, or NB-4 cells were seeded in the complete medium overnight before infection. The viral supernatants containing 10μg/mL polybrene (YEASEN, Cat#40804ES86) were added to the cells and infected by centrifugation. The cells were replaced with fresh medium 24h after infection, and 1μg/mL Puromycin (Solarbio, Cat#P8230) was supplemented to establish stable infected cell lines.

To establish the sgRNA-targeted FASN knockout in single-cell-derived MOLM-13 clones, oligonucleotides encoding the sgRNA were synthesized, annealed, and inserted into Lenti-CRISPR v2. Lentiviral transduced cells were selected with 1μg/ml puromycin for two days, followed by single clone selection through limited dilution in 96-well plates. Single clones were obtained after 2-weeks of selection and expansion, and the knockout efficiency was verified by western blotting. The genomic PCR amplifications were further carried out, and PCR fragments were ligated into pMD-19T (Takara, Cat#6013) for subsequent Sanger sequencing. The targeting sites and sequences of single guide RNA were illustrated in supplementary figure S1 and table S3. The primers for genomic amplifications were 23-Fwd: 5'-AGCCAAGCTGTCAGCCCA-3'; 23-Rev: 5'-AGAGCGGAGGATGAGGAGCC-3'. 4-Fwd: 5'-AGAGCAGCCATGGAGGAGGT-3'; 4-Rev: 5'-AGTGCCTGGGTGGTGAGGA-3'.

#### Cell viability assays

MTT (3-[4,5-dimethylthiazol-2-yl]-2,5-diphenyl-tetra-zolium bromide) assay was used to evaluate the compound-induced cytotoxicity. Briefly, $0.6 \times 10^4$ cells per well were seeded into a 96-well culture plate in 100μL complete medium, and cells were then incubated with appropriate doses of compounds for 72 hours. 10μL of MTT solution (5 mg/mL, Solarbio, Cat#M8180) was loaded and incubated for 4 hours to allow the formazan crystal formation. Subsequently, the solvent (10% SDS, 5% Isobutanol, 0.012 mol/L HCl, dissolved in distilled water) was added to dissolve the purple formazan. The absorbance was measured at 570 nm using an absorbance microplate reader (BioTek, USA). The cell growth inhibition rate was calculated with GraphPad 9.0 software.

#### Cell cycle analysis

The cells were collected and washed once with PBS before staining. 1 mL PBS containing 2 μL RNase A (20 mg/mL) (Thermo Fisher Scientific, Cat#2301455), 10 μL propidium iodide (PI) (1 mg/mL) (Sigma, Cat#P1470), 0.5 μL 10% NP40 (Coolaber, Cat#SL9320) were incubated with samples, and stained at 37℃ for 30 min. The cell cycle profiling was directly acquired by flow cytometry analysis (CytoFlex, Beckman) in biological triplicates. Data were analyzed with FlowJo v10.6.2 software.

### Apoptosis assay

Cell apoptosis assay was performed with Annexin V/PI staining. Briefly, cells were suspended in the Annexin V Binding buffer (Biolegend, Cat#422201) and stained with Annexin V-PB450 (Biolegend, Cat# 640918) (1:200) antibody. After staining at room temperature for 15 minutes, PI dissolved in Annexin V Binding buffer (1:1000) was added before detection. Experiments were performed with three biological replicates, and data were acquired by CytoFlex flow cytometer and analyzed with FlowJo v10.6.2 software.

### Colony formation assay

The cells were seeded in 500 μL Human Methylcellulose Base Media (R&D Systems, Cat#HSC002) containing 2 μg/mL puromycin and 1% penicillin/streptomycin in 12-well plates. The cells were equally distributed by shaking the orifice plate, and colonies were photographed and scored after being cultured at 37℃ and 5% $CO_2$ for 7-10 days. Experiments were performed with four biological replicates for each group.

### Nitro blue tetrazolium (NBT) staining

NBT staining was performed to examine the granulocytic differentiation of leukemic cells, as previously reported (*4, 5*). The cells were collected and re-suspended in a solution containing 2 mg/mL NBT (Beyotime Biotechnology,

Cat#ST362) and 40 ng/mL Phorbol 12-myristate 13-acetate (PMA) (MCE, Cat#HY-18739), incubated at 37℃ for 15 min followed by washing with PBS once. Cells were then centrifuged onto glass slides by Cytospin instrument, air-dried, and re-dyed with sand yellow solution (BaSo, Cat#BA4016D) for 10s. NBT-positive cells were scored when the cell membrane was blue when examined by the microscope. The percentage of $NBT^+$ cells in each group was recorded, with >6 replicates per group.

**Leukemic xenograft mouse models**

MOLM-13 cells transfected with empty vector and FASN-targeting sgRNA or shRNA were re-suspended in PBS. The concentration was adjusted to $2\times10^5$ cells/mL, and 100μL cells per mouse were injected into the NOD. Cg-$Prkdc^{scid}$ $Il2rg^{tm1Wjl}$ Tg (NSGS) mice (8-10 weeks old) through the tail vein. To evaluate the engraftment and leukemic progression, the proportion of human CD45-positive cells in the peripheral blood was assessed by flow cytometry with anti-huCD45-PB450 (BioLegend, Cat# 368540) staining. The survival of mice was recorded, and data were analyzed by the Kaplan-Meier survival curve.

**Bioluminescence imaging**

Bioluminescence imaging was performed using an IVIS Lumina Series III system (PerkinElmer). Briefly, MOLM-13 cells stably expressing luciferase were lentivirally transduced and selected with puromycin, $1 \times 10^5$ cells per mouse were intravenously injected into NKG mice after irradiations (X-ray, 1.5 Gy) to establish the xenograft models. The disease burden of NKG transplants was quantitatively monitored once per week by bioluminescence imaging. Before imaging, the substrate luciferin potassium (Beyotime, Cat#ST196) dissolved in PBS was intraperitoneally injected into mice at a dose of 150 mg/kg. Mice were anesthetized with isoflurane, and whole-body dorsal bioluminescence imaging was performed 10-15 minutes after injection. The leukemic burden for each mouse was measured and quantitated by normalized radiation (p/sec/cm$^2$/sr).

### Murine MLL-AF9 leukemia model

Bone marrow cells were extracted from 8- to 12-week-old *Fasn*$^{fl/fl}$Mx1-cre, *Fasn*$^{fl/+}$Mx1-cre or control mice, and c-kit$^+$ HSPC cells were enriched and expanded in SFEM medium (Stem cell, Cat#09650) with 10 ng/mL IL-3, 10 ng/mL IL-6, 20 ng/mL SCF as previously performed (*6*). After infecting with MSCV-MLL-AF9-IRES-YFP retroviruses in the presence of 10mg/mL polybrene, 2~3 × 10$^5$ cells were transplanted into lethally (7.5Gy) irradiated C57BL/6J mice through retro-orbital injection. 14 days after transplantation, recipients were intraperitoneally injected with three concussive doses of polyIC (Sigma, Cat#P1350) at 6μg/kg in PBS every other day to induce *Fasn* deletion. The survival of transplants was analyzed by the Kaplan-Meier survival curve.

### Competitive Repopulation Assay

BM mononuclear cells (BMMC) (1 × 10$^6$, CD45.2$^+$) from 7- 8 weeks *Fasn*$^{fl/fl}$ Mx1-cre, *Fasn*$^{fl/+}$ Mx1-cre or WT mice plus an equal number of competitor BMMC (1×10$^6$, CD45.1$^+$) were transplanted into lethally irradiated (7.5 Gy) B6.SJL mice (CD45.1$^+$) by retro-orbital injection. Four weeks after transplantation, PB was collected by tail vein bleeding of the recipients and subjected to flow cytometric analysis with anti-CD45.1-APC (BioLegend, Cat#110714) and anti-CD45.2-FITC (BioLegend, Cat#109806) staining. The percentages of CD45.2$^+$ cells in the PB of chimeras were monitored by flow cytometric analysis every 4 weeks post-transplantation. After 16 weeks, the portions of donor-derived cells were further assessed in BM lineage-positive cells and HSPC by flow cytometry.

### Flow cytometric analysis

Flow cytometry analysis of mouse bone marrow and spleen cells was conducted as described previously(*6-9*). In brief, suspended single cells were prepared from bone marrow or spleen for hematopoietic and leukemia cell analysis. The cells were incubated with appropriate antibodies in PBS for 30 minutes on the ice at dark, then suspended in PBS prior to flow cytometric analysis. Anti-TER119-APC (BioLegend, Cat# 116212), Anti-Ly6G-PE-Cy7 (BD Pharmingen, Cat#560601), Anti-CD3e-APC-Cy7 (BD Pharmingen, Cat#557596), Anti-B220-FITC (BioLegend, Cat#103206), Anti-CD11b-PE (BioLegend, Cat#101208) antibodies were used for lineage-positive cell analysis. For analyzing HSPCs, BM $Lin^-$ cells were isolated by Biotin-Mouse Lineage Depletion Cocktail (BD Pharmingen, Cat#558451) and Streptavidin particles plus-DM (BD Pharmingen, Cat#557812) using negative magnetic selection with IMagnet (BD Pharmingen, Cat#552311) according to the manufacturer's instructions. Anti-Sca-1-PE (BioLegend, Cat#108108), anti-CD117-PE-Cy7 (BD Pharmingen, Cat# 558163), anti-CD48-APC-Cy7 (BioLegend, Cat#103432), anti-CD150-APC (BioLegend, Cat#115910), anti-CD127-PE (BioLegend, Cat#135010), anti-Sca1-Pacific Blue (BioLegend, Cat#108120), anti-CD34-FITC (eBioscience, Cat# 2518336), anti-CD135-APC (BioLegend, Cat#135310), anti-CD16/CD32- Brilliant Violet 510 (BioLegend, Cat#101333) antibodies were used for $Lin^-$ cell staining and subsequent HSPC analysis.

**Cellular Thermal Shift Assay (CETSA)**

CETSA was performed as previously described (*10*). Briefly, leukemic cells were seeded ($1\times10^6$ cells) in 10cm dishes and inoculated for 48 hours. The cells were treated with 10μM MS-C19, 10μM TVB-3166 (MCE, Cat#HY-120394), or DMSO as vehicle control for 1 hour at 37°C. Cell lysates were obtained by three repeated freeze-thaw cycles in liquid nitrogen followed by heating for 3 minutes at different temperatures (37°C, 42°C, 47°C, 52°C) using the heating block of

PCR instrument. The target protein in the soluble fraction was separated and quantified by western blotting assay.

### Molecular docking

Molecular docking of MS-C19 or PTM with the FASN-KS domain was performed by AutoDock Vina [11]. AutoDock Tools (The Scripps Research Institute, La Jolla, CA, USA) was used to prepare the ligands and receptor as pdbqt files after removing water, and adding polar hydrogen atoms and Gasteiger charges, respectively. The docking grid box size used was adjusted accordingly to encompass the MS-C19 and PTM interaction sites. Other parameters were used as default. The top docking pose was selected for binding mode comparison. The ligand-protein interaction structures were generated in PyMol (The PyMOL Molecular Graphics System, Version 3.0 Schrödinger, Inc., New York, NY, USA).

### RNA Extraction, Reverse Transcription, and Real-Time PCR.

Total RNA was isolated using Trizol reagent (Yeasen, 10606ES60) and was reverse transcribed using the RevertAid RT cDNA synthesis kit (Thermo Scientific, K1691). Quantitative PCR analyses were performed on QuantStudio 1 Real-time PCR instrument (ABI) using SYBR Green Master Mix (Yeasen, 11184ES08). Relative amounts of target transcripts were normalized to the housekeeping gene, human 18S rRNA or GAPDH, and calculated by the 2(−ΔΔCt) method. Primer sequences are listed in Supplemental Table S2.

### Western blotting

Standard western blotting was carried out as described previously (*8, 9*). Briefly, cell lysates were prepared by lysing the cells with cell lysis buffer (50 mM HEPES, pH7.6, 250 mM NaCl, 5 mM EDTA, pH8.0, 0.1% NP40) containing protease inhibitor cocktail (#M5293, AbMole Bioscience) on ice for 30 minutes followed by once freeze/thaw cycle at -80°C. The protein concentration was determined by the BCA protein Assay kit (#E-BC-K318-M, Elabsciences), and an equal amount of protein was prepared with the sample buffer. Samples were denatured by boiling for 10 min and separated on the SDS-PAGE gel. Subsequently, the proteins were transferred to a 0.45 μm PVDF membrane (#IPVH00010, Millipore). After blocking with 5% milk for 1 hour at room temperature, the membrane was incubated with appropriate primary antibodies at 4°C overnight. The blots were washed and incubated with HRP-conjugated secondary antibodies (Abiowell) for 1 hour at room temperature and the HRP signals were detected by the ECL chemiluminescence detection kit (#P1020, Applygen). The following antibodies were used: anti-FASN (#10624-2-AP, Proteintech), anti-HSC70 (sc-7298, Santa Cruz), and anti-GAPDH (#AP0066, Bioworld).

**RNA-sequencing**

MOLM-13 cells transduced with shFASN or empty vector after selection with 1μg/mL puromycin for two days or treated with either vehicle control or MS-C19 for 48 hours were collected and extracted total RNA by E.Z.N.A. Total RNA Kit I (Omega, # R6934-01). The RNA amount and purity were quantified using NanoDrop ND-1000. RNA integrity was further assessed by Bioanalyzer 2100 (Agilent) with RIN number >7.0, and confirmed by electrophoresis with denaturing agarose gel. Poly (A) RNA was purified for library construction, and the average size for the cDNA library was 300 ± 50 bp. Paired-end sequencing (PE150) was performed on an Illumina Novaseq™ 6000 (LC-Bio Technology, Hangzhou, China). The expression levels of all transcripts and mRNAs as calculated by FPKM were estimated by StringTie. The significant differentially expressed mRNAs were defined as fold change >2 with q value<0.05. GO and KEGG pathway analyses were performed using clusterProfiler and illustrated by R.

#### Luciferase assay

The dual luciferase activity assay was conducted as previously described (*6, 9*). HEK293T cells (1.05 × $10^5$ cells per well) were incubated in 48-well plates 14-18 hours before transfection. Cells around 70%-80% confluence in each well were co-transfected with pGL4.22 empty vector or constructs containing *GRN* genomic fragments (100 ng) together with pLV3 empty vector or an increased amount of human TFEB expressing construct (pLV3-hTFEB, 50 ng, 150 ng, and 300 ng) by using Neofect DNA transfection reagent. *Renilla* luciferase expression vector pRL-TK was co-transfected for internal control (1ng/well). Then, the cells were harvested by using a passive lysis buffer (200 μL/well ) 24 hours post-transfection, and luciferase activity was measured using the Dual-Luciferase Reporter Assay System (E1910, Promega) and GloMax 20/20 Luminometer (Promega). Firefly luciferase activities were normalized to Renilla luciferase activities (Firefly/Renilla) and calculated as the fold-change from the empty vector. All the luciferase experiments were performed in triplicate.

#### CUT&Tag sequencing

CUT&Tag was performed by using a commercial kit (Ruoyu Biotech., CUT-01) following the manufactory's protocol. Briefly, 0.5 million cells were collected, and incubated with activated concanavalin A-coated magnetic beads. The primary antibody employed was a mouse-derived anti-TFEB antibody (Santa Cruz, sc-166736, 1:33). The goat anti-mouse antibody (1:100) served as the secondary antibody. pAG-Tn5 was used for CUT&Tag reactions. Extracted DNA fragments were utilized for library construction followed by quantification and sequencing (Novogene).

#### Quantitative proteomics

Label-free quantitative proteomics was performed at the core facility of Xiamen University. Briefly, for mass spectrometry, samples were subjected to in-solution trypsin digestion and dried. Samples were then analyzed on an EASY-nLC 1200 (Thermo SCIENTIFIC) coupled to an Orbitrap Fusion Lumos (Thermo SCIENTIFIC) equipped with an EASY-IC ion source. Peptides were dissolved in 10 µl 0.1% formic acid and were auto-sampled directly onto a homemade C18 column (35 cm×75 µm i.d., 2.5 µm 100Å). Samples were then eluted for 120 mins with linear gradients of 3–35% acetonitrile in 0.1% formic acid at a flow rate of 300nl/min. For the data-independent acquisition, a 25 m/z precursor isolation width covering 400-1200 m/z was adopted. The DIA files were generated using DIA-NN software133 (V.1.8.1) against a UniProtKB human protein database without decoys, during which the parameters were set as: "FASTA digest for the library-free search" and "Deep learning-based spectra and RTs prediction" were enabled: Protease, "Trypsin/P"; Missed cleavage, "2"; N-termM excision, "checked"; c-carbamidomethylation, "checked"; Moxidation, "checked". The maximum mass accuracy tolerances were set to 10 ppm for both MS1 and MS2 spectra. Protein inference in DIA-NN was the protein name (fasta). The quantification mode was set to "Robust LC (high accuracy)". All other settings were set as default.

### Lipidomic analysis

**Lipid extraction.** 1 × $10^6$ cells were collected and frozen rapidly by liquid nitrogen. To extract total lipids, 400 µL MTBE and 80 µL pre-cooled methanol containing internal standards were added to each sample followed by vortex for 30 seconds and sonication for 10 minutes at 4°C. After centrifugation for 15 minutes at 3000g and 4°C, 300 µL supernatant enriched with lipids was aspirated and dried using a nitrogen blower maintained at room temperature. The dried lipids were then stored at -80°C for long-term preservation until further analysis. Preceding the LC-MS analysis, the lipids were resuspended in a 150 μL solvent mixture comprising of 20μL solubilizing agent ($CH_2Cl_2$: MeOH=2:1, v/v) and 130 μL of a diluent formulated with ACN, IPA, and $H_2O$ (65:30:5, v/v/v) containing 5 mM ammonium acetate. After vortex for 30 seconds followed by centrifugation, 70 μL supernatant was subjected to analysis. Additionally, a pooled aliquot of supernatants from each sample was combined as a quality control (QC) reference sample.

**Lipidomics data acquisition and analysis.** Lipidomics data were acquired using a Vanquish Flex UPLC system coupled with a Thermo Scientific Orbitrap Exploris 240 mass spectrometer equipped with a heated electrospray ionization (H-ESI) source. Samples were separated through a BEH C8 column (2.1 × 100 mm with a 1.7 μm particle size, Waters, Milford, MA, USA) with the column temperature maintained at 55°C. The mobile phases consisted of 2 mM ammonium formate in mobile phase A (40% water and 60% acetonitrile) and mobile phase B (90% isopropanol and 10% acetonitrile). The gradient started with 1.5 min of isocratic elution with 32% B (and 68% A). Phase B was increased to 85% over the next 15.5 min and then from 85% B to 97% B in only 0.1 min, and maintained at 97% B for 2.4 min. Rapidly, the mobile phase composition was returned to 32% B within 0.1 min and maintained for 5 min for column post-equilibration. The flow rate for the mobile phases was set at 0.26 mL/min. The injection volume was 2 μL for positive ions and 5 μL for negative ions.

The mass spectrometer was operated in positive or negative modes using a full scan/data-dependent secondary scan (Full-ddMS2) in the scan range m/z 100–1500 Da. Capillary voltage was set at 3400 V for positive and 3000 V for negative modes. Ion Transfer Tube Temp: 320 °C, Vaporizer Temp: 350 °C, sheath gas: 40 Arb, Aux gas: 10 Arb, Sweep Gas: 10 Arb. Orbitrap resolution was set to 120,000 in MS1 and 15,000 in MS2. The normalized collision energy type was selected. The internal mass calibration was maintained throughout the analysis. The MS data were captured and processed using Xcalibur software and LipidSearch 5.1 software (Thermo Fisher Scientific, San Jose, USA). The total number of lipids and the number of lipid classes were summarized using the online web tool LipidSuite (https://suite.lipidr.org/). The intensity distributions of SFA, MUFA, and PUFA in certain lipid classes across samples are displayed as histograms. The percentage of SFA, MUFA, and PUFA in certain lipid classes between samples was analyzed by the online web tool LipidSig (https://lipidsig.bioinfomics.org/). Differential analysis of lipidomic data was performed using the R package "limma". Lipids with a P value < 0.05 and fold change < -1.5 or > 1.5 were statistically considered significant.

### Statistical Analysis

All the statistical data are expressed as means ± standard error of the mean (SEM). The student's t-test was used for significance testing between the two groups. The ANOVA test was used to compare the significance between different groups. The log-rank Mantel-Cox test was used to compare survival curves generated using Kaplan-Meier algorithms. Experiments were performed in biological triplicates unless otherwise noted. All the statistical analyses were performed using GraphPad Prism 9.0. P < 0.05 were considered statistically significant. Asterisks indicate *P < 0.05, **P < 0.01, ***P < 0.001 and ****P < 0.0001.

**Data availability.** The data supporting the findings of this study are available within the article and its supplemental material. The RNA-Seq, proteomics, and lipidomics data were deposited in the National Genomics Data Center under accession number PRJCA031722. Any additional information required to reanalyze the data reported in this paper is available upon request.

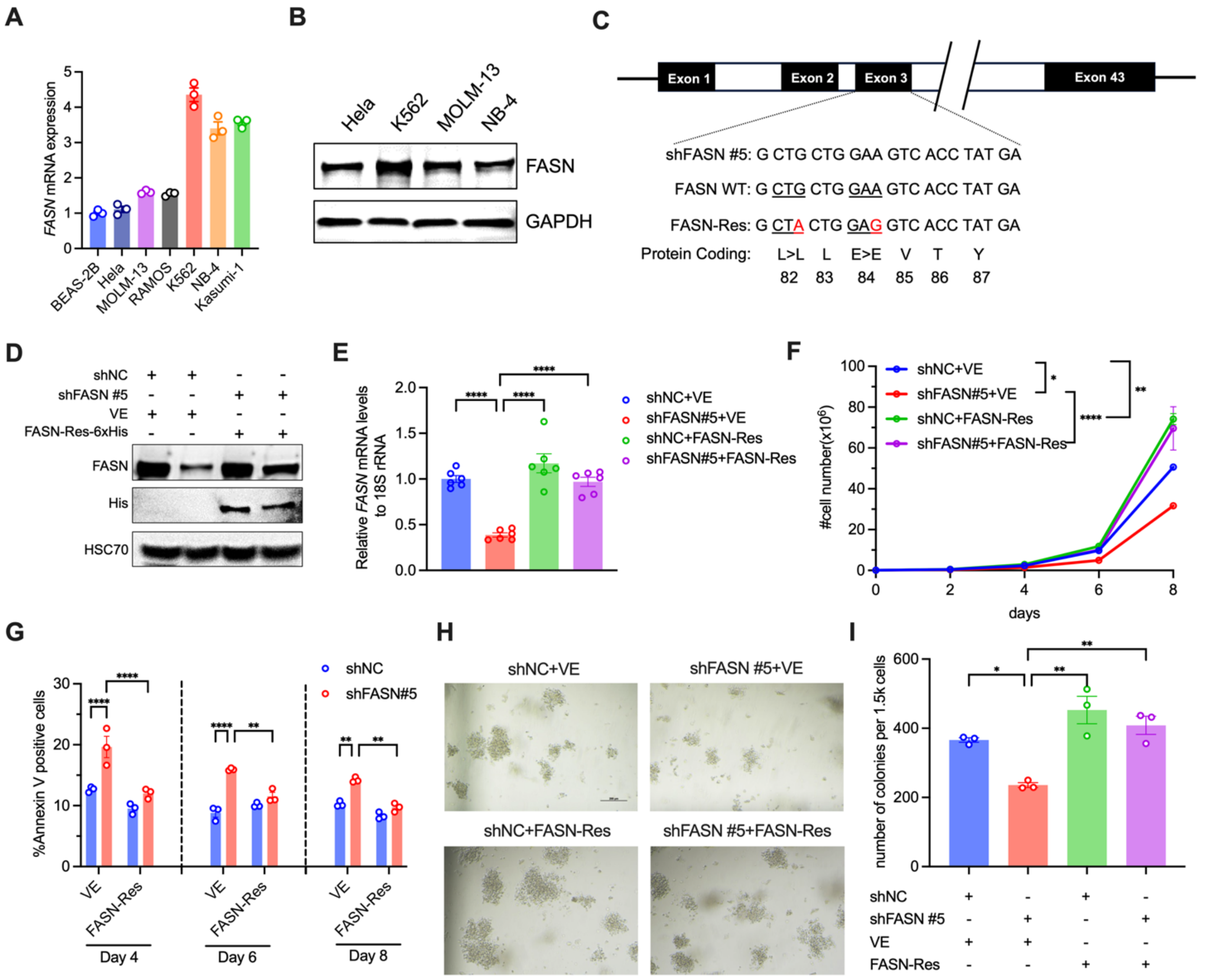

**Figure S1. shRNA-resistant FASN rescued cell growth defects due to FASN deficiency.** (**A**-**B**) FASN mRNA or protein levels from indicated leukemic cell lines were determined by quantitative PCR or Western blotting in A and B, respectively. (**C**) Schematic diagram showing the construction of a synonymous FASN mutant intended to confer resistance to shFASN #5. (**D**-**E**) Determination of FASN protein and mRNA levels of indicated MOLM-13 cells by Western blotting (D) and quantitative PCR (E). (**F**-**I**) The cells from D were examined for growth curve (F), percentages of apoptotic cells (G), and colony formation ability (H-I). The representative images of colonies from the indicated group were shown in H. Two-way ANOVA in F and G, one-way ANOVA in E and I with Tukey's corrections in all. *, P<0.05. **, P<0.01. ***, P<0.001. ****, P<0.0001.

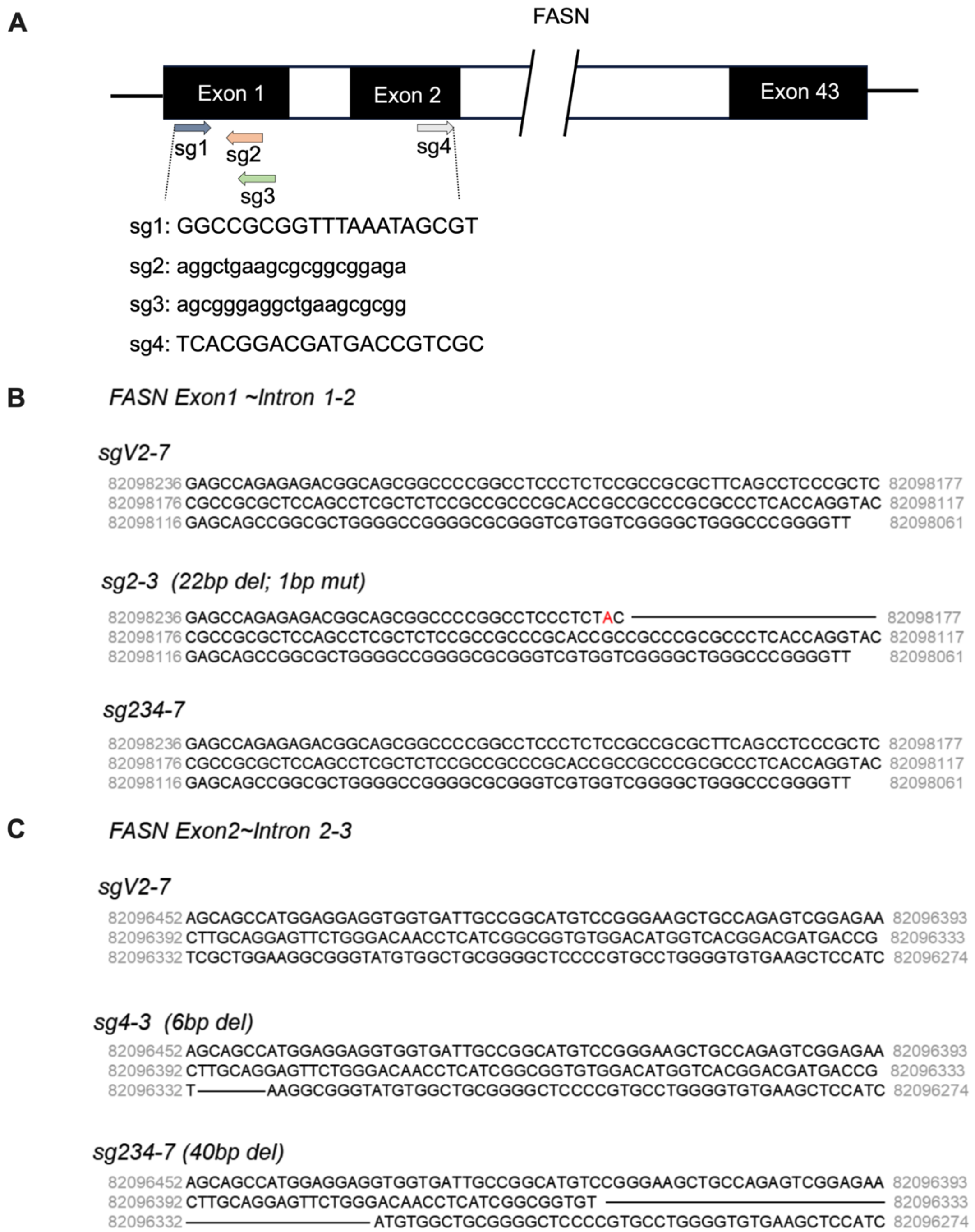

**Figure S2. CRISPR-Cas9/sgRNA-targeting in the human FASN gene.** (**A**) Schematic of the Cas9/sgRNA-targeting site in the human *FASN* genomic locus. (**B-C**) The fragments surrounding the target sites were PCR-amplified using genomic DNA extracted from indicated single-clone derived MOLM-13 cells followed by Sanger DNA sequencing for INDELs validation.

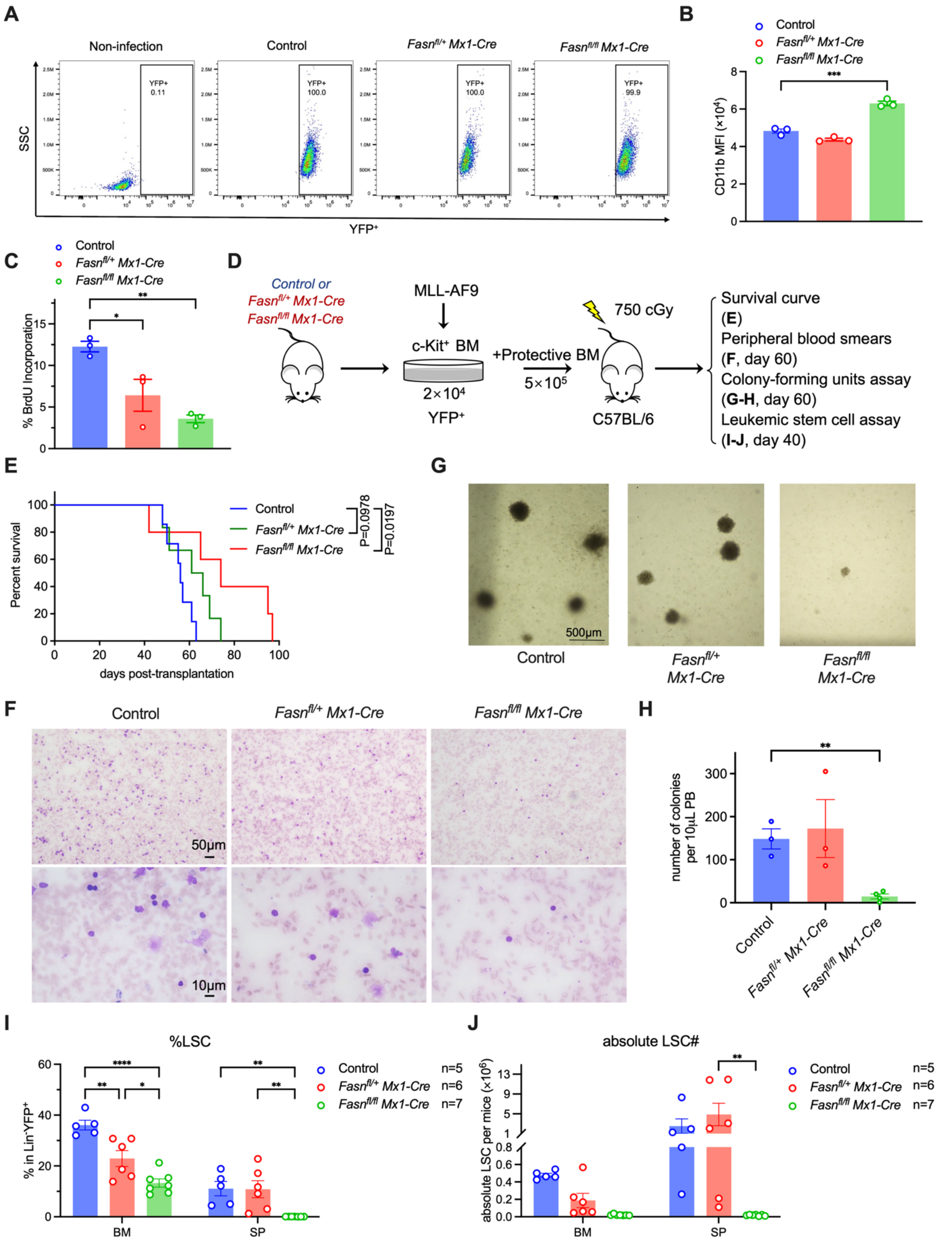

**Figure S3. Fasn deletion delayed the leukemic progression in vivo.** (**A**) Percentages of YFP-positive cells as determined by flow cytometric analysis 14

days after MLL-AF9 transduction. (**B-C**) The myeloid differentiation (B) and BrdU incorporation (C) were assessed in vitro using cells from A. (**D**) Schematic of the routines to establish leukemic mouse models using YFP+ cells. (**E**) Survival curve of indicated transplants as in D. n=7 in control, n=6 in *Fasn*$^{fl/+}$ *Mx1-Cre*, n=5 in *Fasn*$^{fl/fl}$ *Mx1-Cre*. (**F**) Peripheral blood smears were examined by Giemsa staining at day 60 after transplantation. Scale bar, 50 μM (upper) and 10 μM (lower). (**G**-**H**) Colony formation assay of peripheral blood from indicated recipient mice. The representative images were shown in G, and the numbers of colonies were scored under an inverted microscope and shown in H. Scale bar, 500 μM. n=3 in control, n=3 in *Fasn*$^{fl/+}$ *Mx1-Cre*, n=4 in *Fasn*$^{fl/fl}$ *Mx1-Cre*. (**I-J**) Frequency of leukemic stem cells (LSCs) in lineage negative and YFP positive cell population from the indicated transplants was determined by flow cytometric analyses in I. The counts of LSCs were further quantified as in J. n=5 in control, n=6 in *Fasn*$^{fl/+}$ *Mx1-Cre*, n=7 in *Fasn*$^{fl/fl}$ *Mx1-Cre*. One-way ANOVA in B and C with Tukey's corrections; Mantel-Cox (Log-Rank) statistical analysis in E; Unpaired student t-test in H; Two-way ANOVA in I and J with Tukey's corrections. *, P<0.05. **, P<0.01. ***, P<0.001.****, P<0.0001.

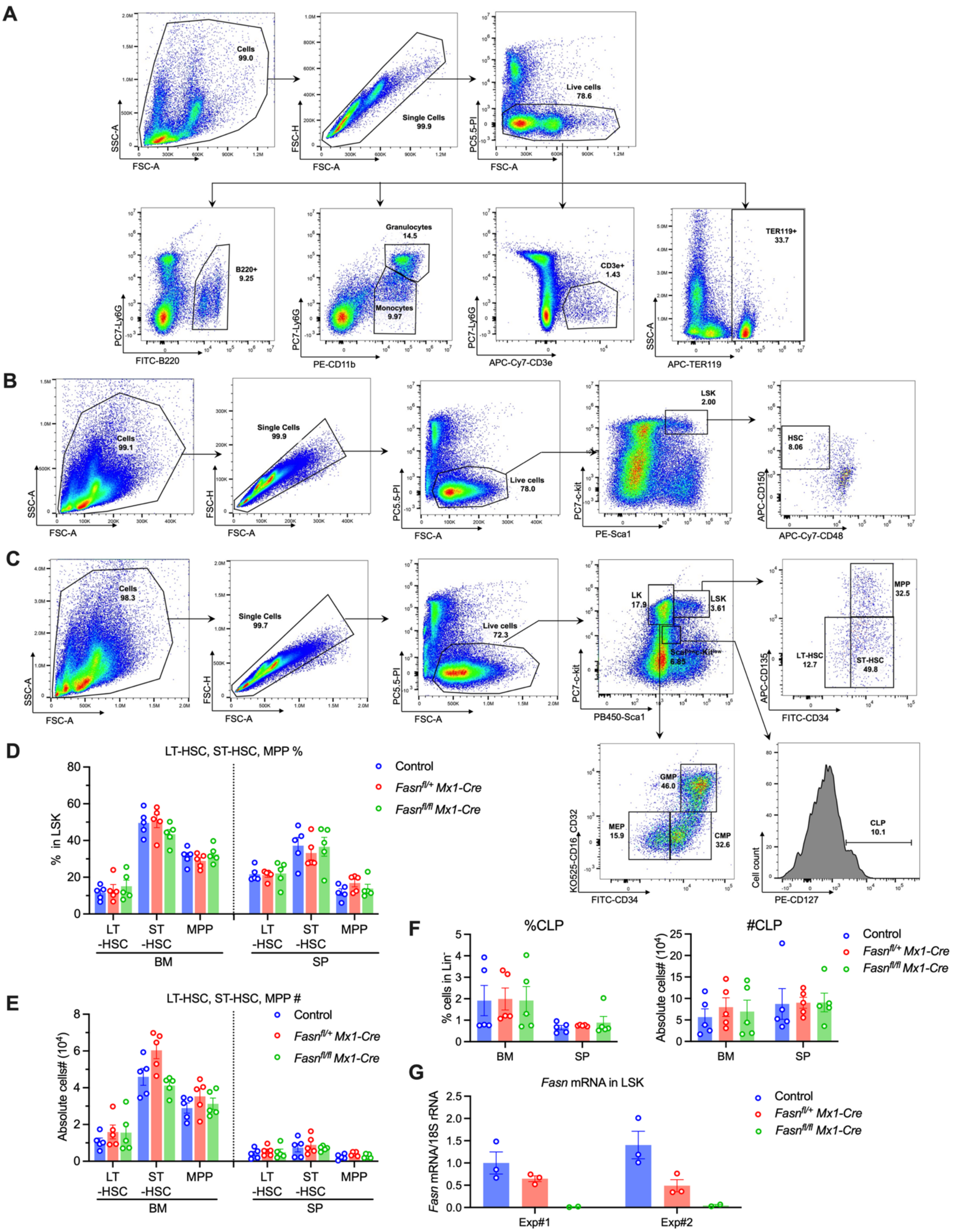

**Figure S4. HSPCs compositions in *Fasn* conditional knockout mice.** (**A-C**) Flow cytometric gating strategies for analyzing lineage-positive cell populations

(A), HSC (B) or LT-HSC, ST-HSC, and progenitor cells (MPP/CMP/GMP/MEP/CLP) (C). (**D**-**E**) Percentages and cell counts of LT-HSC, ST-HSC, and MPP in bone marrow or spleen of indicated mice determined by flow cytometric analysis were shown in D and E, respectively. LT-HSC, long-term HSC. ST-HSC, short-term HSC. MPP, multipotent progenitors. (**F**) Same as D and E, except CLP was analyzed. n=5 in each group for D, E, F. (**G**) Donor-derived LSK cells ($CD45.2^+$) from competitive transplants after polyIC administration were sorted and analyzed for *Fasn* transcripts by qPCR at week 16. 18sRNA, internal control.

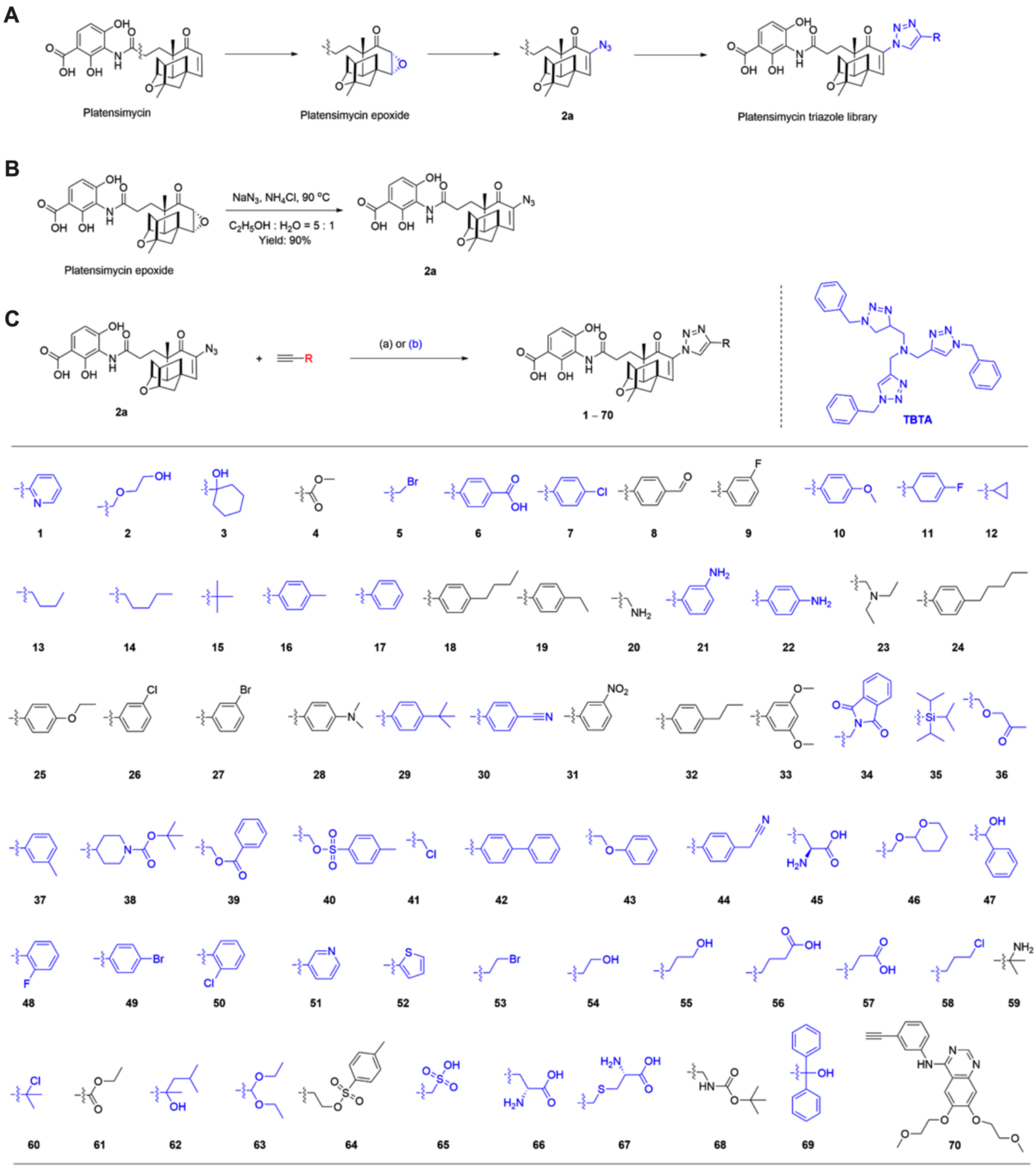

**Figure S5. Construction of PTM derivatives library.** (**A**) Synthetic strategies for constructing a focused platensimycin triazole library through click chemistry. (**B**) Synthesis routes of 6-azidyl substituted platensimycin 2a. (**C**) Preparation of a 70-membered platensimycin triazole library. (a, marked in black, 20 compounds) $CuSO_4$ (0.02 equiv), sodium ascorbate (0.1 eq.), $H_2O$: tertiary butanol (1:1), rt, 95% from **2a**. (b, marked in blue, 50 compounds) $CuSO_4$, 0.5

eq (from 100 mg/mL aqueous solution, sodium ascorbate (2.5 equiv, from 100 mg/mL $H_2O$), $H_2O$: DMF (1:1), TBTA 1 eq. yield>95% from **2a**. The yields were based on UPLC-ESI-MS analysis of the crude reaction products.

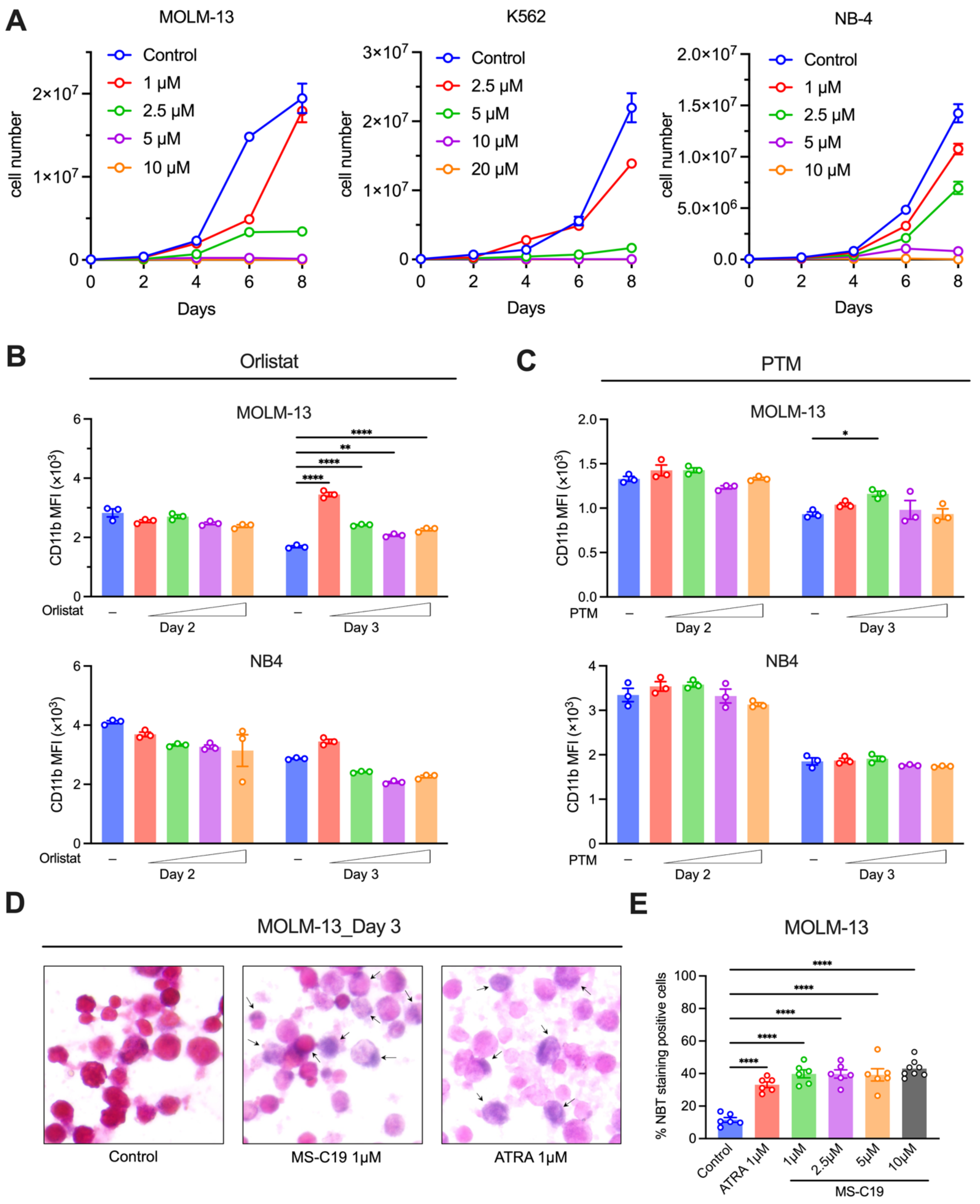

**Figure S6. Cell growth and myeloid differentiation in leukemic cells treated with FASN inhibitors.** (**A**) *In vitro* cell growth curves of MOLM-13, K562, and NB4 treated with indicated concentrations of MS-C19. (**B**-**C**) Leukemic cells treated with varied concentrations (1μM, 2.5μM, 5μM, 10μM) of either Orlistat or PTM were examined for CD11b expression by flow cytometric assay. (**D**-**E**) Differentiated MOLM-13 cells induced by MS-C19 or ATRA were assessed by NBT staining. The representative images were shown in D with arrows indicating the differentiated cells. NBT staining positive cell percentages were further quantified in E from 6-8 randomly selected fields per group. The statistical analysis was performed by two-way ANOVA in B and C, and one-way ANOVA in E with Tukey's corrections in all. *, $P<0.05$. **, $P<0.01$. ****, $P<0.0001$.

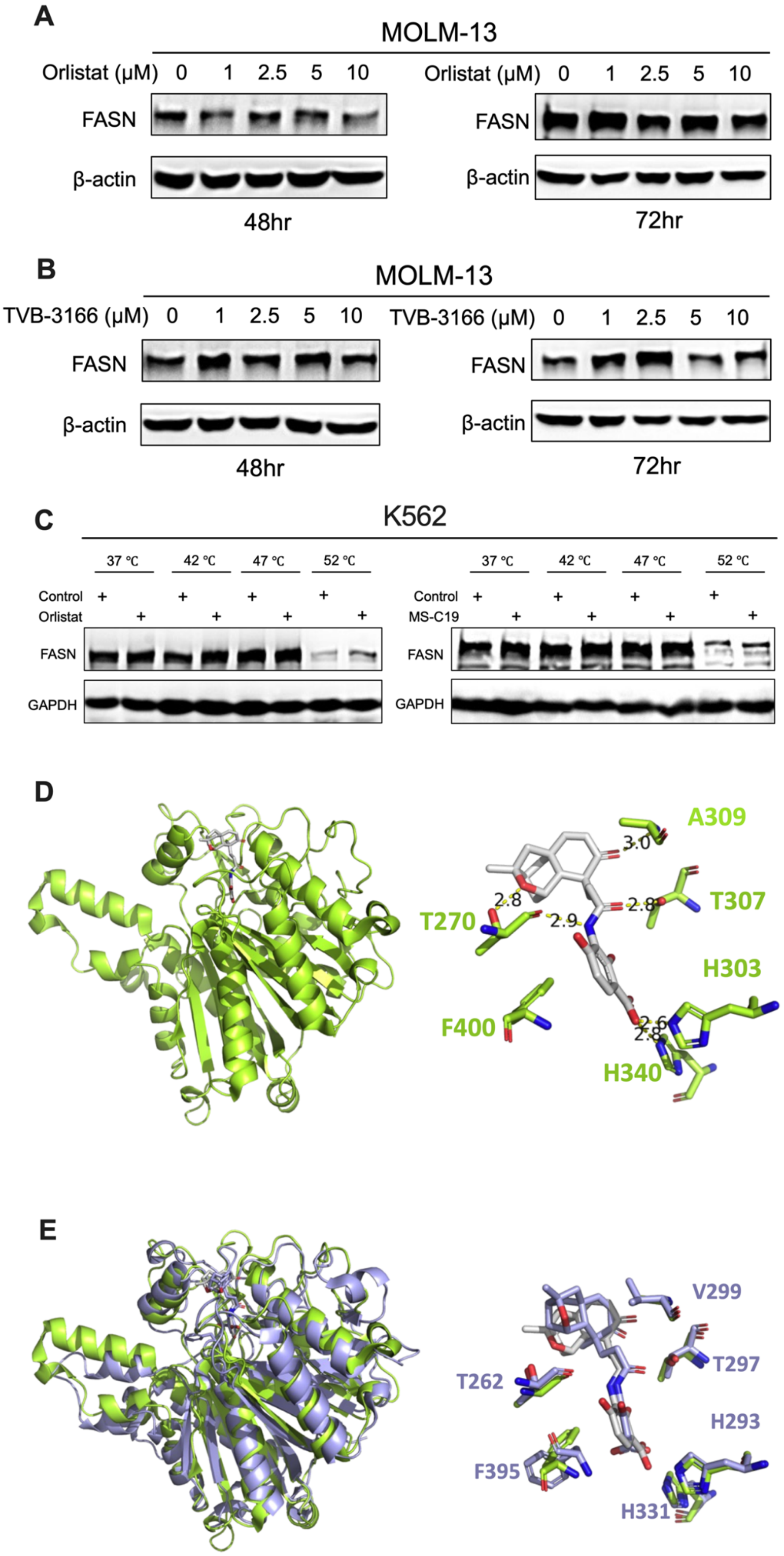
A
MOLM-13
Orlistat (μM) 0 1 2.5 5 10
FASN
β-actin
48hr
Orlistat (μM) 0 1 2.5 5 10
FASN
β-actin
72hr
B
MOLM-13
TVB-3166 (μM) 0 1 2.5 5 10
FASN
β-actin
48hr
TVB-3166 (μM) 0 1 2.5 5 10
FASN
β-actin
72hr
C
K562
37 ℃ 42 ℃ 47 ℃ 52 ℃
Control
Orlistat
FASN
GAPDH
37 ℃ 42 ℃ 47 ℃ 52 ℃
Control
MS-C19
FASN
GAPDH
D
A309
T307
T270
H303
F400
H340
3.0
2.8
2.9
2.8
2.6
2.8
E
V299
T297
T262
H293
F395
H331

557

**Figure S7. Expression of FASN protein levels upon FASN inhibitor treatments.** (**A**-**B**) MOLM-13 cells challenged with Orlistat (A), or TVB-3166 (B) at indicated concentrations for either 48 or 72hr were collected and subjected to Western blotting with indicated antibodies. β-actin, loading control. (**C**) Cellular thermal shift assay to test the binding of Orlistat or MS-C19 with FASN. K562 cells were treated with 10μM inhibitors at 37°C for 1hr followed by incubation at the indicated temperature for 3 min. Western blotting analysis was then performed for FASN. GPDH, loading control. (**D**) Crystal structure of PTM with ecFabF(C163Q), PDB code: 2GFX. The amino acid residues binding with PTM in the ecFabF active site are labeled. (**E**) Structural alignment of ecFabF(C163Q)/PTM and FASN-KS domain/PTM. The model of the FASN-KS domain/PTM is generated by the AutoDock Vina program. The structure of the FASN-KS domain is from PDB code: 3hhd. Conserved amino acid residues binding with PTM in the FASN KS domain are labeled.

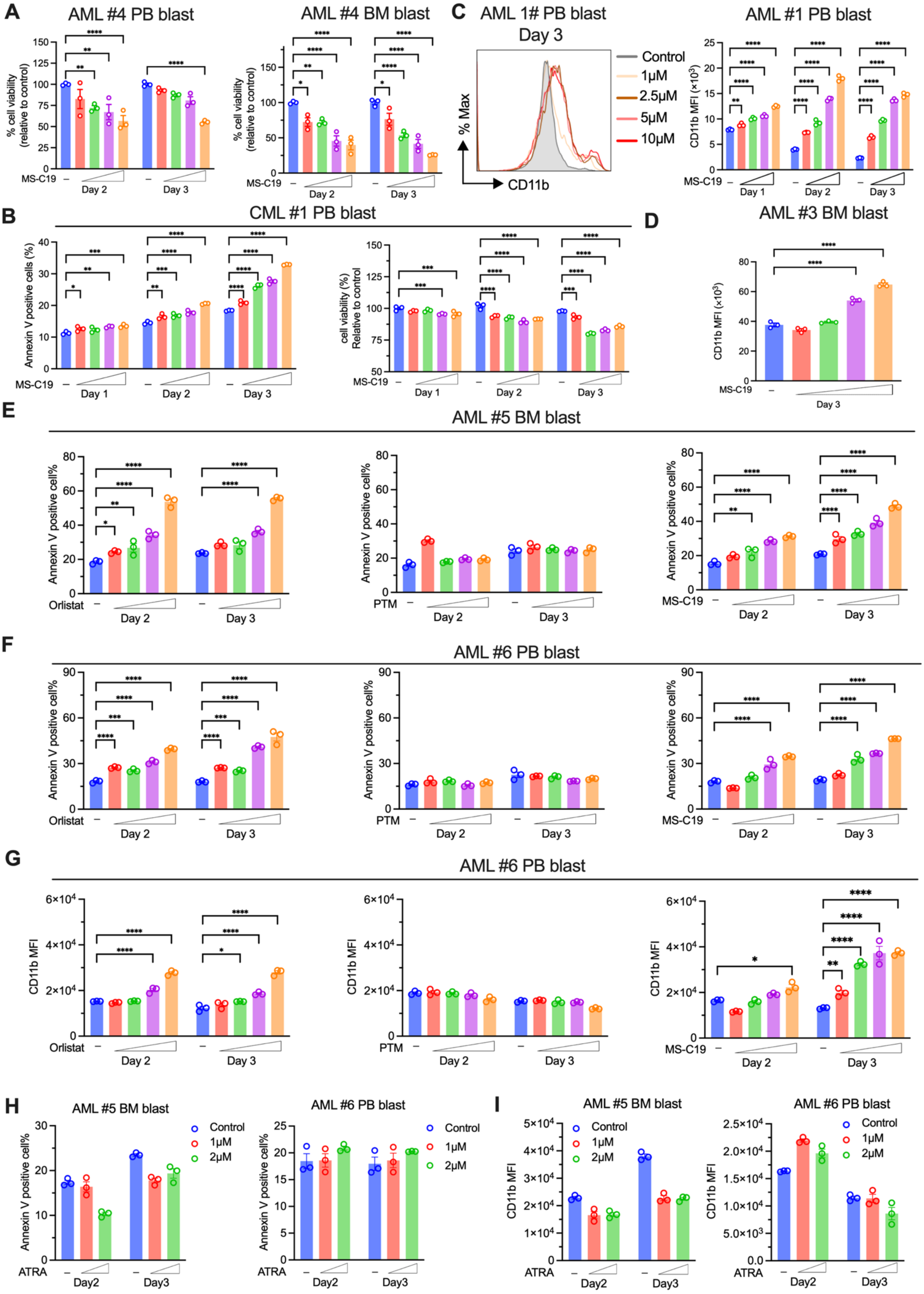
A
AML #4 PB blast
AML #4 BM blast
% cell viability (relative to control)
MS-C19
Day 2
Day 3
C
AML 1# PB blast Day 3
Control
1μM
2.5μM
5μM
10μM
% Max
CD11b
AML #1 PB blast
CD11b MFI (×10³)
Day 1
B
CML #1 PB blast
Annexin V positive cells (%)
cell viability (%) Relative to control
D
AML #3 BM blast
E
AML #5 BM blast
Annexin V positive cell%
Orlistat
PTM
F
AML #6 PB blast
G
AML #6 PB blast
CD11b MFI
H
AML #5 BM blast
AML #6 PB blast
ATRA
Day2
Day3
I
AML #5 BM blast
AML #6 PB blast

**Figure S8. Evaluations of FASN inhibitors within the clinical leukemic blast cells.** (**A**) Leukemic blast cells isolated from AML patients were untreated or treated with MS-C19 (1μM, 2.5μM, 5μM, 10μM) for 2 or 3 days, cell viabilities were determined by flow cytometric analysis of PI and Annexin V staining. (**B**) Same as A, except peripheral leukemic blast cells from CML patients were treated with MS-C19 (2.5μM, 5μM, 10μM, 20μM). (**C**-**D**) Myeloid differentiation of leukemic blast cells from either peripheral blood (C) or bone marrow (D) of AML patients was treated with MS-C19 (1μM, 2.5μM, 5μM, 10μM) at indicated timepoint, and CD11b expression was determined by flow cytometric analysis. (**E**-**G**) Leukemic blast cells from AML patients treated with Orlistat, PTM, and MS-C19 at 1μM, 2.5μM, 5μM, and 10μM were examined for apoptosis (E and F) and myeloid differentiation (G). (**H**-**I**) The same assay as in E and G, except the leukemic blast cells of AML patients were treated by ATRA at indicated concentrations.

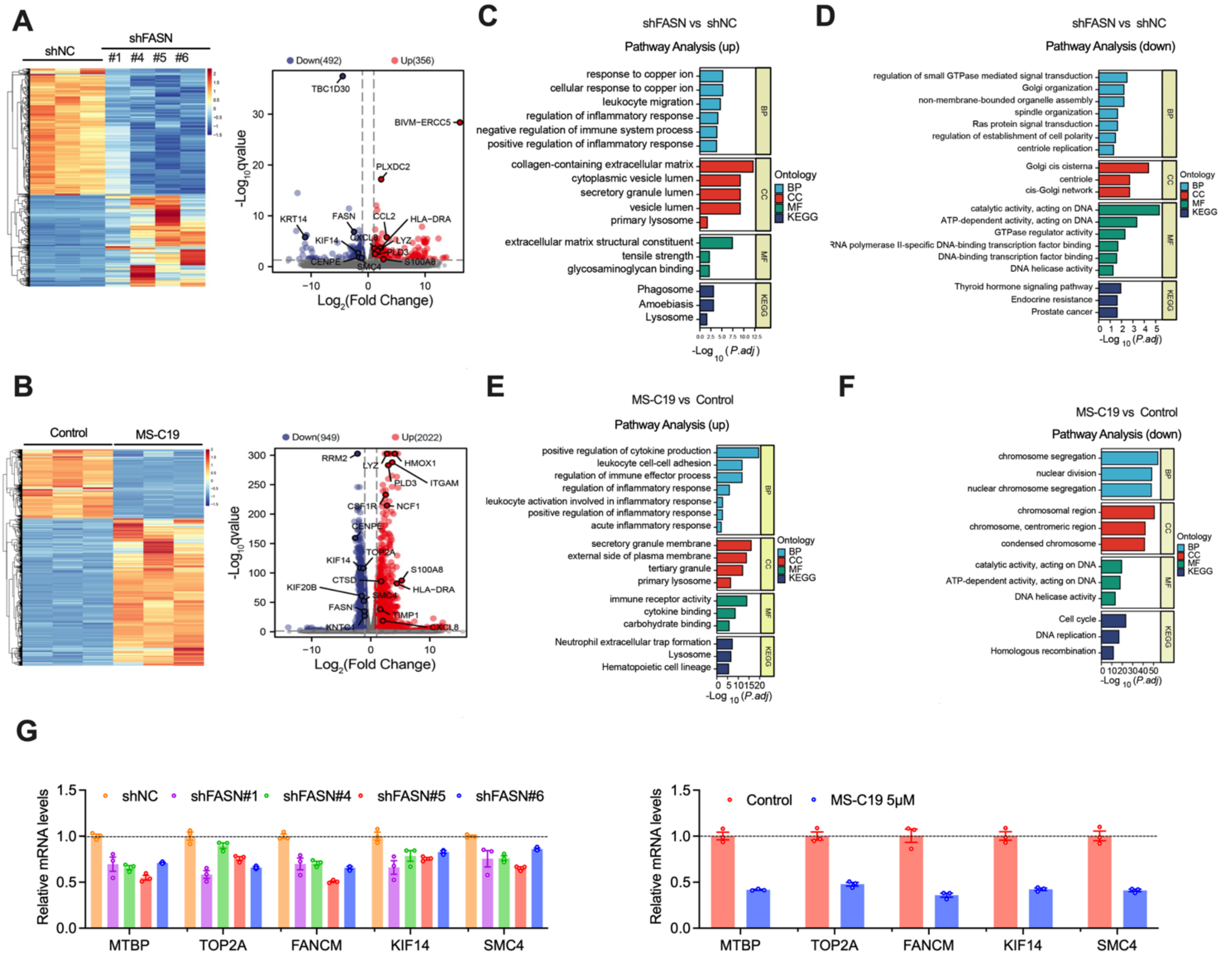

**Figure S9. GO and KEGG pathway analysis of differentially expressed genes in RNA-seq.** (**A**) Heatmap with hierarchical clustering (left) and volcano plot (right) deciphering the differential expressed genes between control and FASN depleted MOLM-13 cells at 48hr. (**B**) Same as A, except MS-C19 treated (5μM, 48hr) and untreated MOLM-13 cells were used for RNA-seq. (**C-F**) Pathway enrichment analysis of DEGs upon FASN KD (C, upregulated. D, downregulated) or MS-C19 treatment (E, upregulated. F, downregulated) in MOLM-13 cells. (**G**) Validation of downregulated DEGs in indicated cells by qPCR.

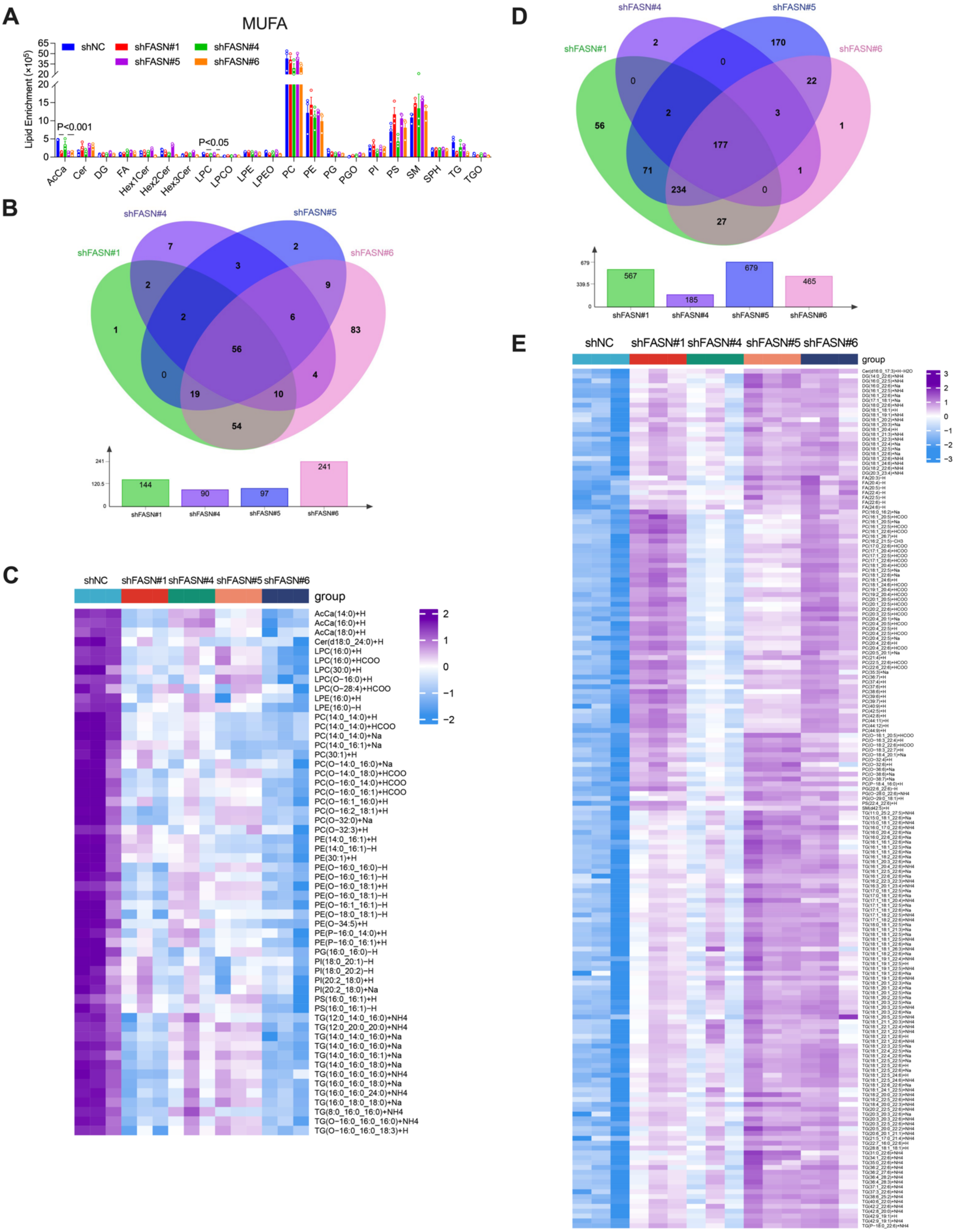

**Figure S10. Lipidomic analysis of MOLM-13 cells depleted of FASN.** (**A**) Quantification of various classes of monounsaturated fatty acids (MUFA) from the indicated cells. The P value was calculated using an unpaired student t-test between shFASN and shNC. (**B**-**E**) Venn plots and hierarchical-clustering heatmaps show significantly declined (B-C) or increased (D-E) lipids in FASN KD cells when compared with shNC. The cutoff threshold is $FC > 1.5$ or $< 0.5$, and $P < 0.05$. The number of differential lipids is shown as a histogram in the lower panel of B and D.

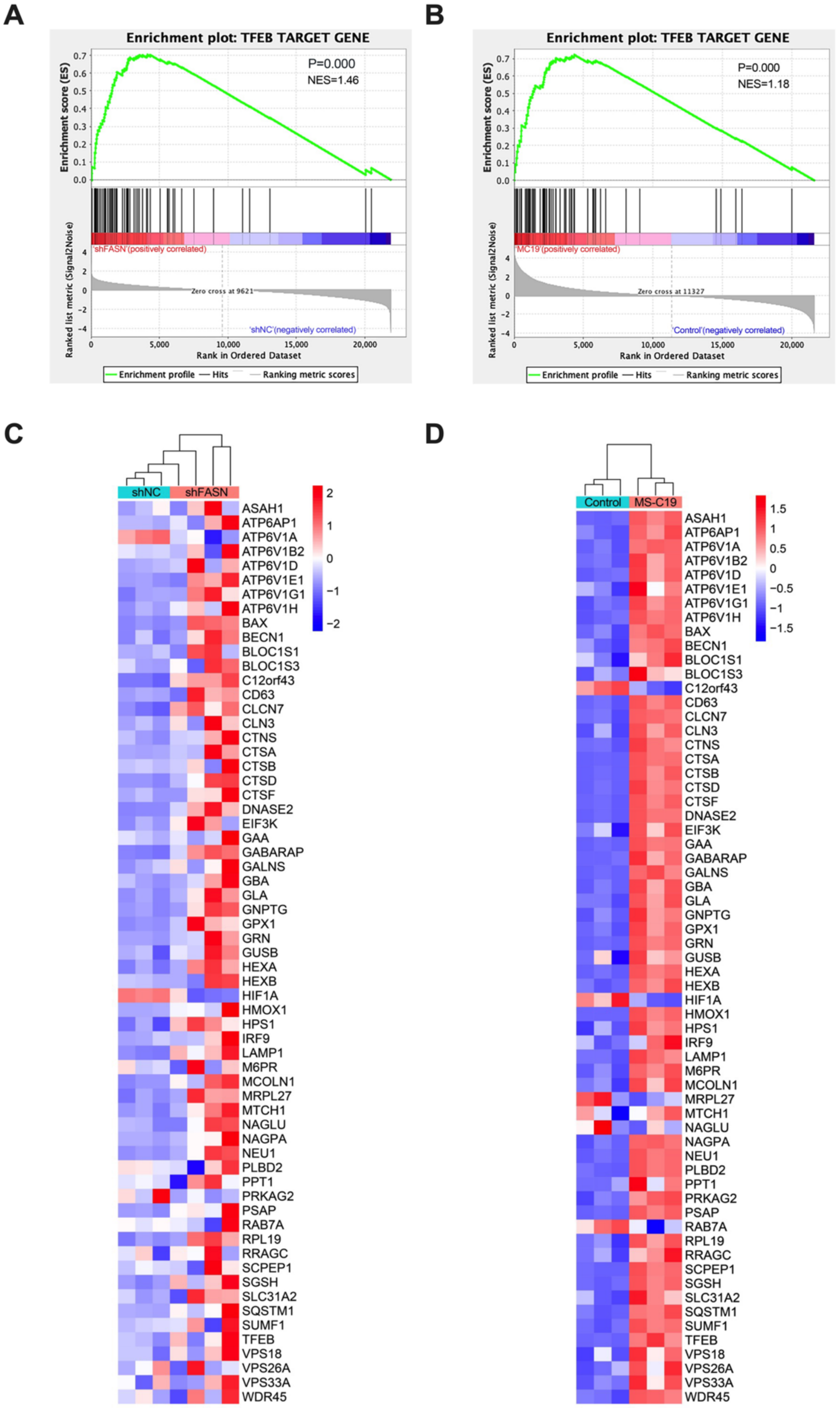

**Figure S11. Enrichment of TFEB transcription signature upon FASN inactivation.** (**A-B**) Gene Set Enrichment Analysis (GSEA) of TFEB target genes in FASN KD (A) or MS-C19 (B) treated cells compared with the control group. (**C-D**) Heatmaps illustrating the expression profiles of TFEB target genes.

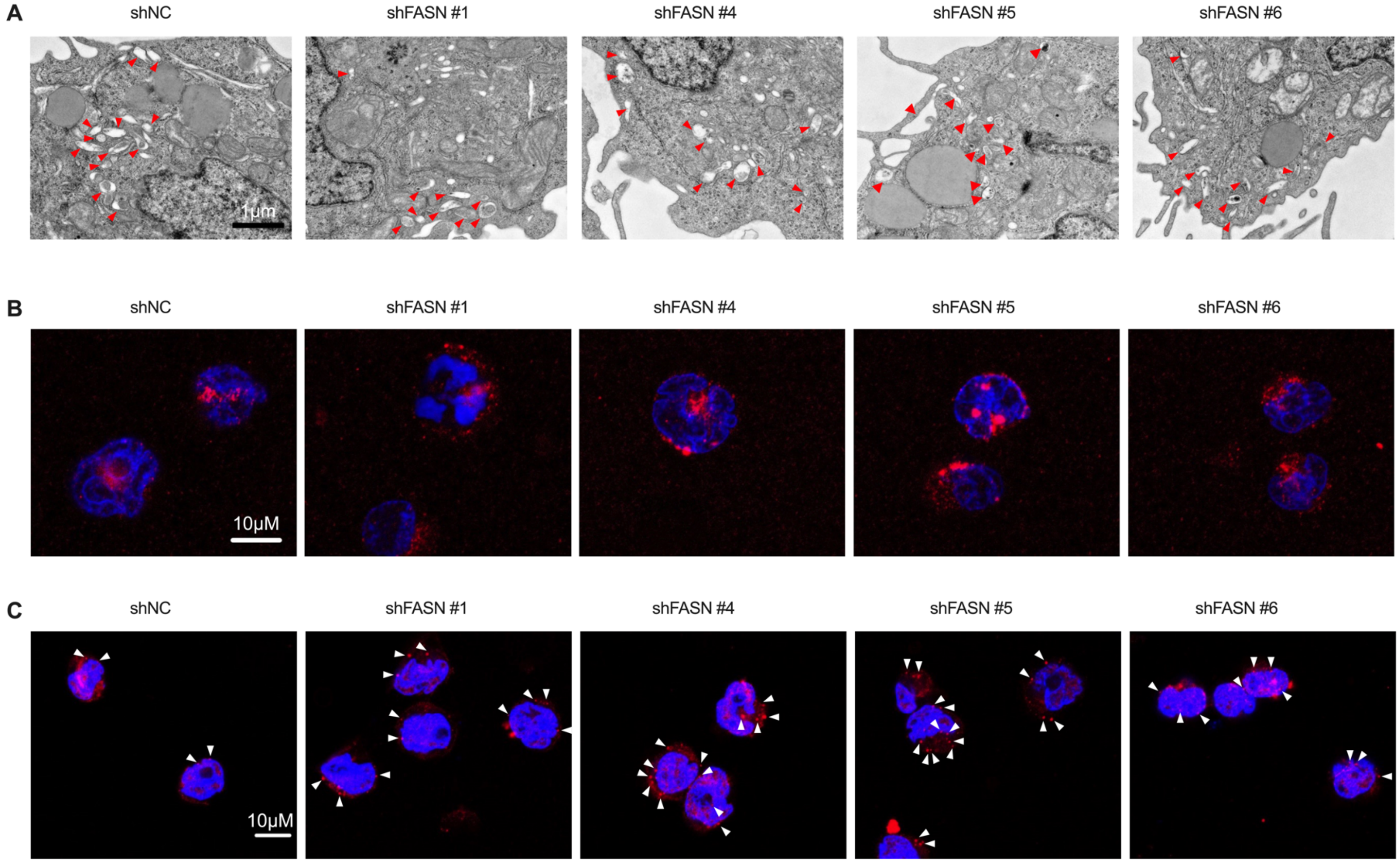

**Figure S12. Quantity and quality assessments of lysosome in FASN depleted MOLM-13.** (**A**) Visualization of lysosomes was performed by scanning electron microscope (SEM), Representative images were shown and lysosomes are indicated by arrowheads. Scale bar, 1μm. (B) Immunofluorescent staining of LAMP1 in FASN KD and control cells. Scale bar, 10µm. (**C**) Same as B, except Glactin-3 was examined. The arrowheads indicate the Glactin-3-positive puncta.

619

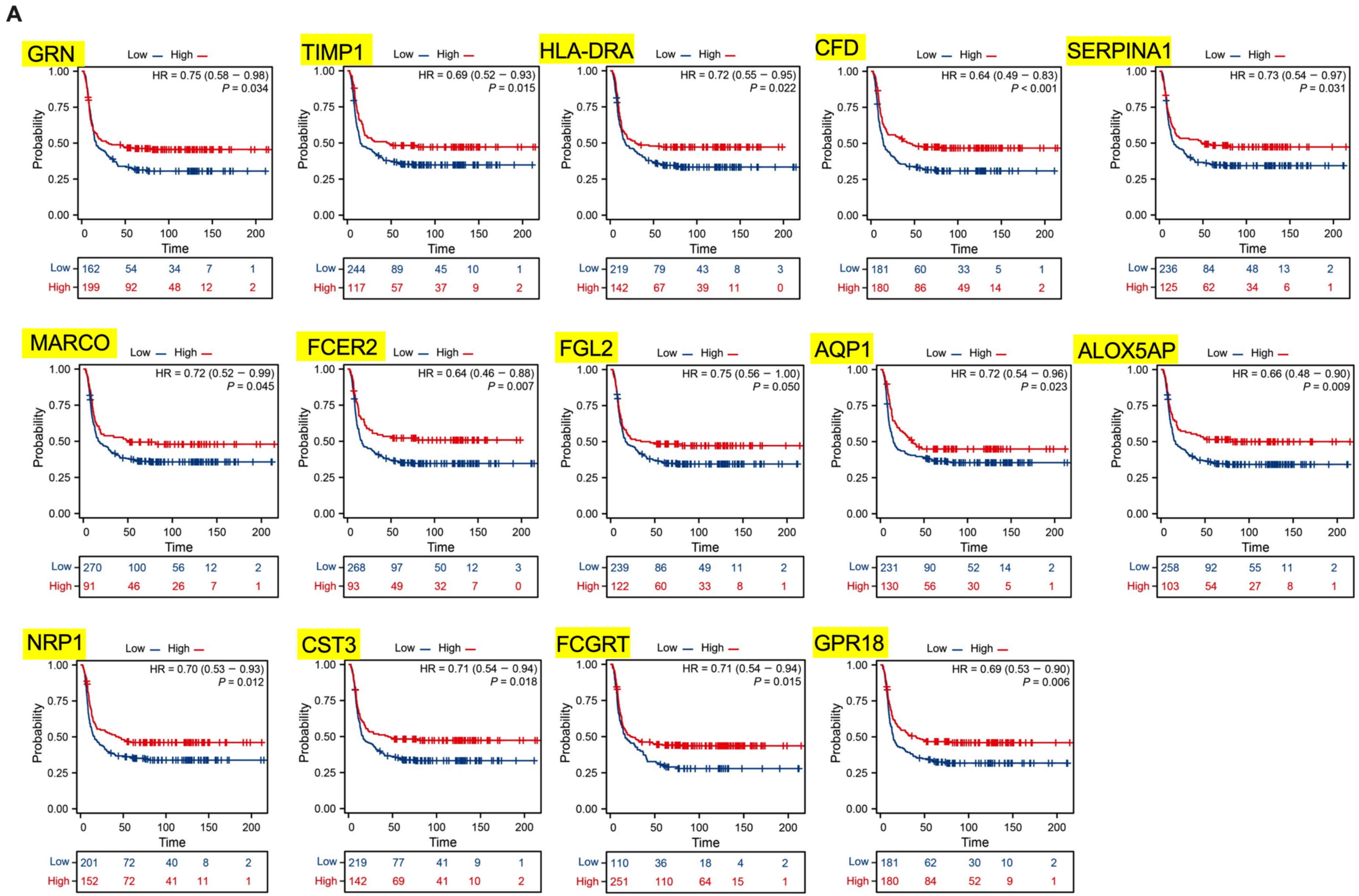
A
GRN
HR = 0.75 (0.58 − 0.98)
P = 0.034
TIMP1
HR = 0.69 (0.52 − 0.93)
P = 0.015
HLA-DRA
HR = 0.72 (0.55 − 0.95)
P = 0.022
CFD
HR = 0.64 (0.49 − 0.83)
P < 0.001
SERPINA1
HR = 0.73 (0.54 − 0.97)
P = 0.031
MARCO
HR = 0.72 (0.52 − 0.99)
P = 0.045
FCER2
HR = 0.64 (0.46 − 0.88)
P = 0.007
FGL2
HR = 0.75 (0.56 − 1.00)
P = 0.050
AQP1
HR = 0.72 (0.54 − 0.96)
P = 0.023
ALOX5AP
HR = 0.66 (0.48 − 0.90)
P = 0.009
NRP1
HR = 0.70 (0.53 − 0.93)
P = 0.012
CST3
HR = 0.71 (0.54 − 0.94)
P = 0.018
FCGRT
HR = 0.71 (0.54 − 0.94)
P = 0.015
GPR18
HR = 0.69 (0.53 − 0.90)
P = 0.006
Low — High —
Probability
Time
Low
High

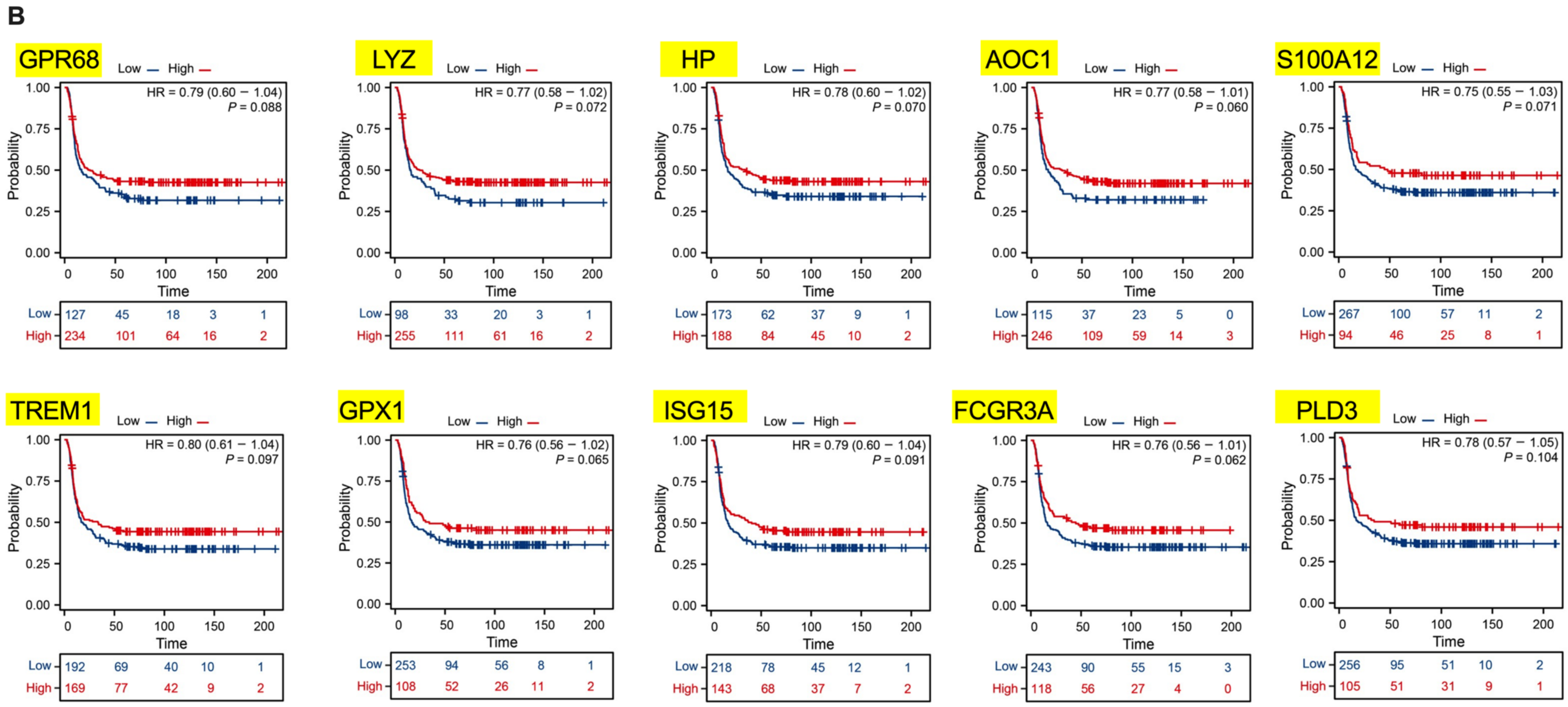
B
GPR68
HR = 0.79 (0.60 − 1.04)
P = 0.088
LYZ
HR = 0.77 (0.58 − 1.02)
P = 0.072
HP
HR = 0.78 (0.60 − 1.02)
P = 0.070
AOC1
HR = 0.77 (0.58 − 1.01)
P = 0.060
S100A12
HR = 0.75 (0.55 − 1.03)
P = 0.071
TREM1
HR = 0.80 (0.61 − 1.04)
P = 0.097
GPX1
HR = 0.76 (0.56 − 1.02)
P = 0.065
ISG15
HR = 0.79 (0.60 − 1.04)
P = 0.091
FCGR3A
HR = 0.76 (0.56 − 1.01)
P = 0.062
PLD3
HR = 0.78 (0.57 − 1.05)
P = 0.104
Low
High
Probability
Time

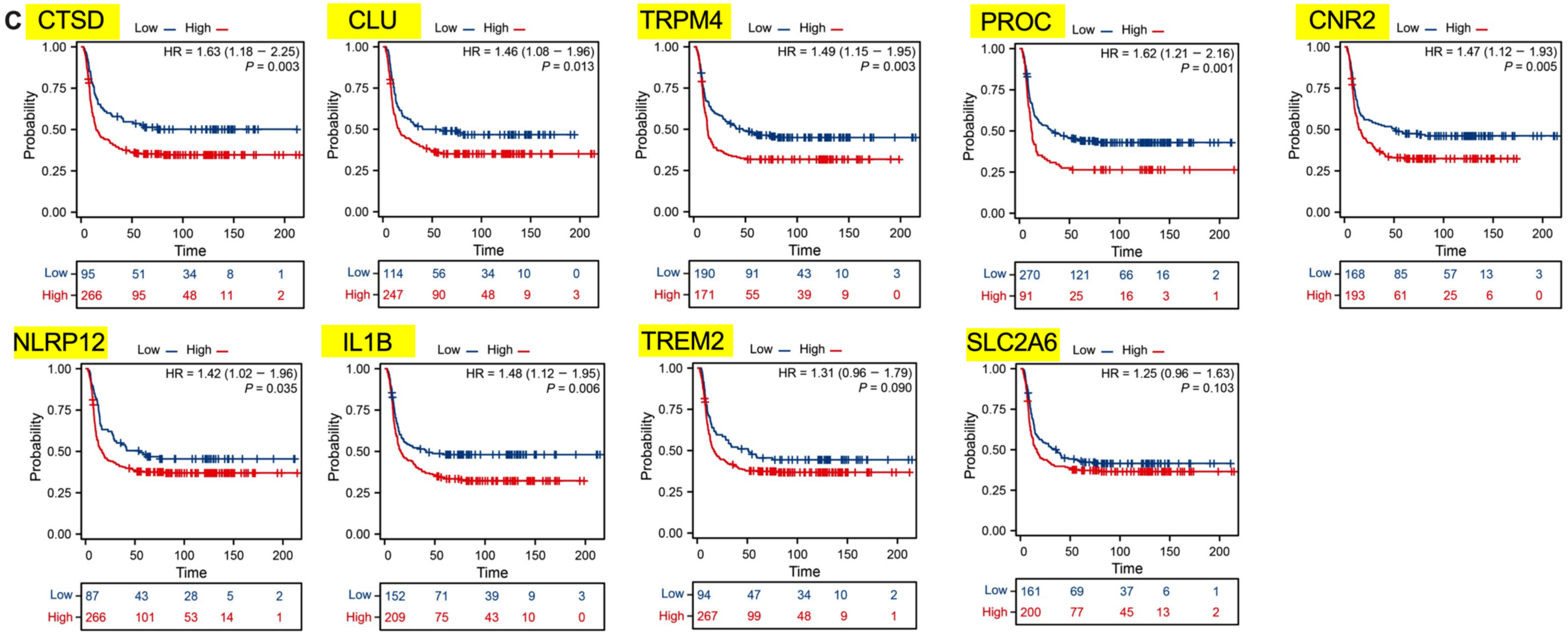

**D**

| Correlations | Gene symbol | % of Total (66) | |
|---|---|---|---|
| Positive (P<0.05) | GRN, TIMP1, HLA-DRA, CFD, SERPINA1, MARCO, FCER2, CTS3, ALOX5AP, AQP1, FGL2, NRP1, FCGRT, GPR18 | 21.21% (14) | 56.07% |
| Positive (0.05<P≤0.11) | PLD3, LYZ, HP, AOC1, S100A12, FCGR3A, TREM1, GPX1, ISG15, GPR68 | 15.16% (10) | |
| Positive (P>0.01) | S100A8, FUCA1, C3, ORM2, DNASE2, ACP5, CTSH, CHI3L1, FCAR, TYROBP, STAP1, HMOX1, FCAR | 19.70% (13) | |
| Negative (P<0.05) | CTSD, CLU, TRPM4, PROC, CNR2, NLRP12, IL1B | 10.61% (7) | 22.73% |
| Negative (0.05<P≤0.11) | TREM2, SLC2A6 | 3.03% (2) | |
| Negative (P>0.01) | OLR1, CXCL8, GPR183, IL1R1, AGT, GPR35 | 9.09% (6) | |
| Undefined | H2BC12, JAML, LILRB3, LILRA6, COL6A3, IRF9, C1QA, SGSH, GIMAP5, SLC26A11, CRISPLD2, LGALS3BP, HLA-DRB1, FN1 | (14) | 21.21% |

Note: The colored genes in red are shared by both inflammation and lysosomal activity pathways.

**Figure S13. Association of AML patient survival with genes enriched in lysosome and inflammatory pathways.** (**A**-**B**) Kaplan-Meier survival curves show the positive (A, with significance, $P<0.05$; B, $0.05<P\leq0.11$) or negative (**C**) correlations between overall survival of AML patients and indicated gene expression levels. Data were extracted from the online KMPlotter. (**D**) Table summary of the correlation analysis as in A-C.

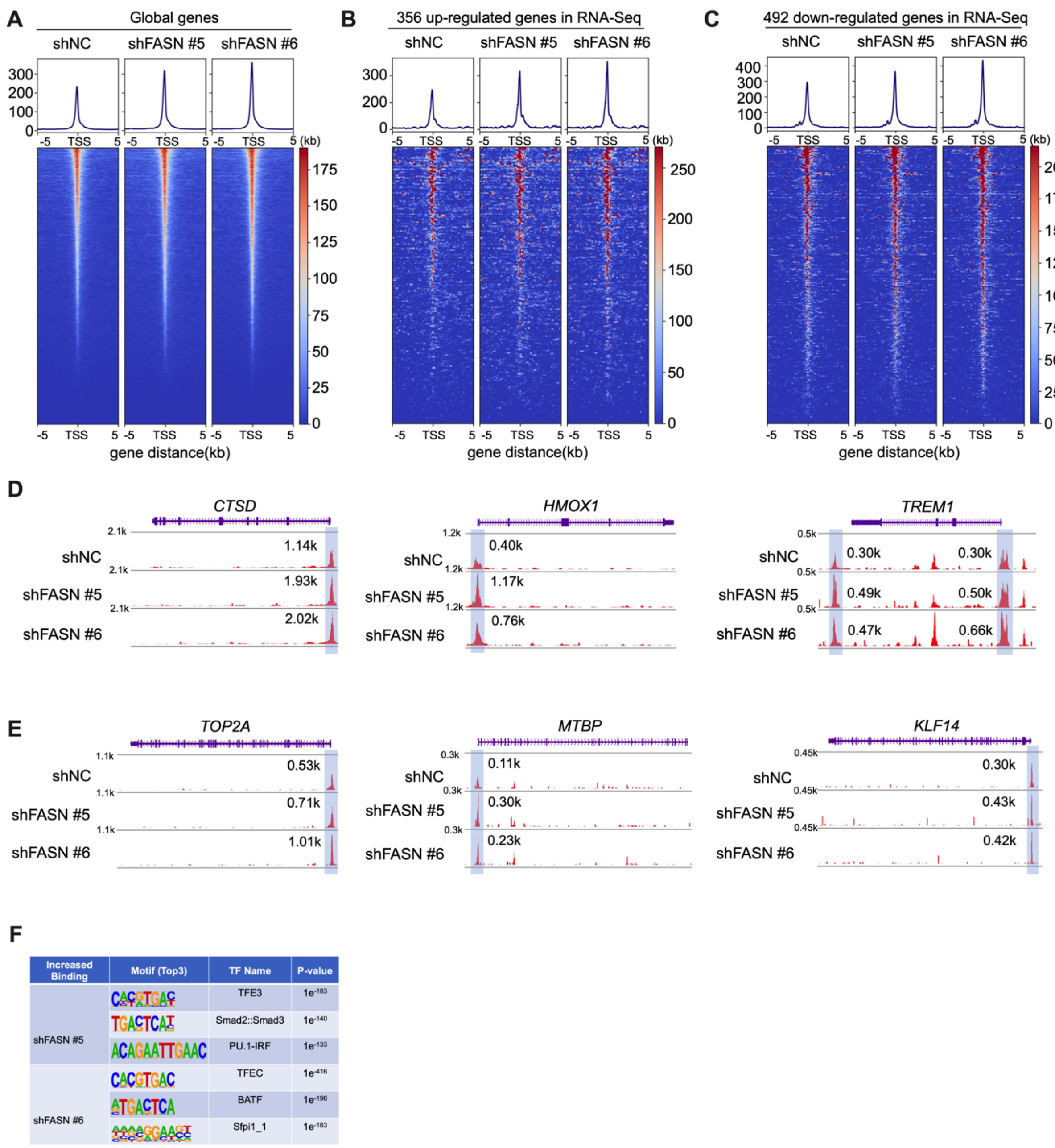

**Figure S14. Genomic landscape of TFEB occupancy revealed by CUT&Tag sequencing analysis.** (**A**-**C**) TFEB CUT&Tag profile and heat map showing TSS ± 5 kb. (**D**-**E**) Genome browser views illustrating the augmented binding intensity of TFEB on the genomic locus of both upregulated genes (*CTSD*, *HMOX1*, *TREM1*) and down-regulated genes (*TOP2A*, *MTBP*, *KLF14*) in MOLM-13 cells with FASN KD. (**F**) The top cis-regulatory elements predicated from TFEB binding peaks as recalled from FASN KD cells.

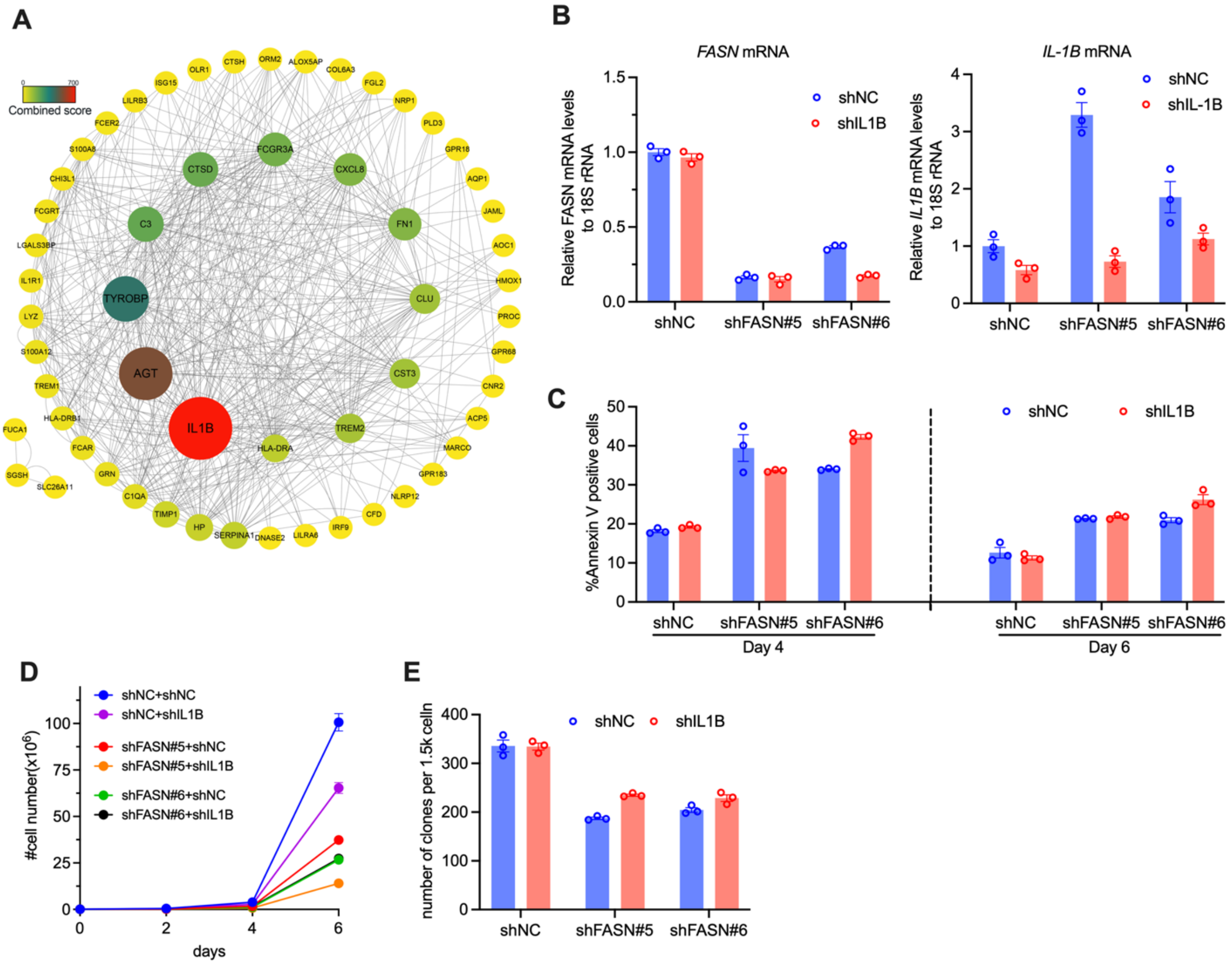

**Figure S15. IL-1B was dispensable for FASN-deficiency-induced leukemic cell death.** (**A**) Protein-protein interaction network. (**B**) *FASN* and *IL-1B* mRNA levels were determined by qPCR in control or FASN-depleted MOLM-13 cells with or without IL-1B knockdown. (**C**-**E**) The cells from B were analyzed for cell apoptosis(C), cell growth (D), and clonogenicity (E). Experiments were performed in triplicates.

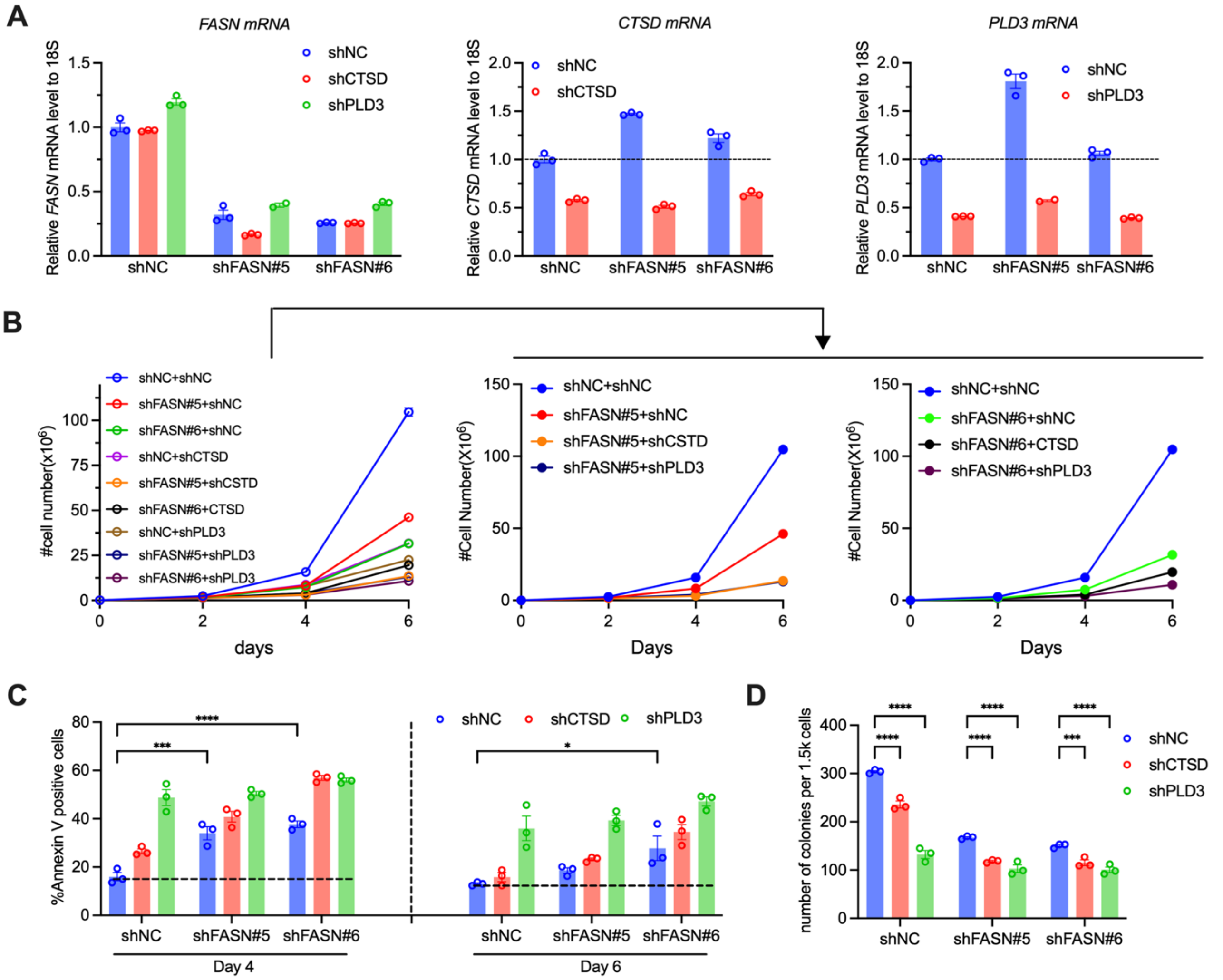

**Figure S16. CTSD or PLD3 expression was not required for FASN-depletion-induced cell death.** (**A**) *FASN*, *CTSD,* and *PLD3* mRNA levels in control or FASN-depleted MOLM-13 cells in combination with indicated shRNA. (**B-D**) The cell growth curves (B), cell apoptosis (C), and clonogenicity (D) were analyzed for the cells from A. Experiments were performed in triplicates. Two-way ANOVA in C and D with Tukey's corrections. *, P<0.05. **, P<0.01. ****, P<0.0001.

**Supplementary Table S1**. Optimization of click chemistry reaction conditions for synthesis compound 17.

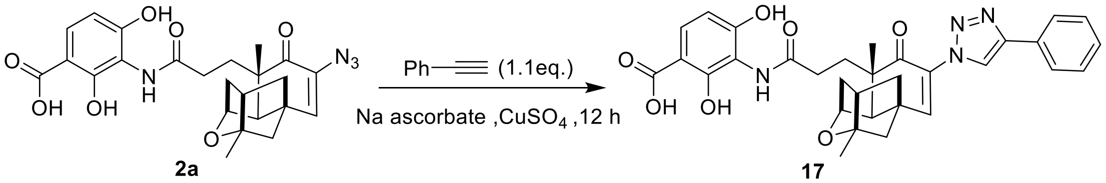

| Entry | Solvent | T(°C) | Ligand | Gas | Yield[a] (%) |
|---|---|---|---|---|---|
| 1 | t-BuOH: H2O (1:1) | 25 | / | air | <5 |
| 2 | t-BuOH: H2O (1:1) | 50 | / | air | <5 |
| 3 | t-BuOH: $H_2O$ (1:1) | 80 | / | air | <5 |
| 4 | DMSO: $H_2O$ (1:1) | 80 | / | air | <5 |
| 5 | DMF: $H_2O$ (1:1) | 80 | / | $N_2$ | <5 |
| 6 | DMF: $H_2O$ (1:1) | 25 | TBTA | $N_2$ | > 95 |
| 7 | DMF: $H_2O$ (1:1) | 25 | TBTA | air | > 95 |

[a] Yields were calculated according to HPLC analysis.

**Supplementary Table S2. Characteristics of clinical specimens.**

| Identifier | Sex | Age | Gene Mutations | Cytogenetics |
|---|---|---|---|---|
| Healthy 1 | F | 16 | n/a | n/a |
| AML remission #1 | M | 15 | n/a | complex |
| AML 1 | F | 47 | MLL-PTD | n/a |
| AML 3 | M | 63 | ASXL1, NRAS, SRSF2, STAG2, TET2 | normal |
| AML 4 | M | 49 | AML1-ETO + | n/a |
| AML 5 | F | 48 | FLT3-ITD+, IDH1 | n/a |
| AML 6 | M | 51 | NF1 | n/a |
| CML 1 | M | 32 | BCR-ABL p210 | t (9;22) (q34; q11) |
| CML 2 | M | 34 | BCR-ABL | t (9;22) (q34; q11) |
| APL 1 | F | 72 | PML-RARA | t (15;17) (q24; q21) |

656 **Supplementary Table S3.** Primer and shRNA sequences.

| Gene | Sequence (5'-3') | Comments |
|---|---|---|
| FASN | AAGGACCTGTCTAGGTTTGATGC | qPCR primers |
| | TGGCTTCATAGGTGACTTCCA | |
| GAPDH | AGGTCGGTGTGAACGGATTTG | qPCR primers |
| | TGTAGACCATGTAGTTGAGGTCA | |
| 18s rRNA | GCAATTATTCCCCATGAACG | qPCR primers |
| | GGCCTCACTAAACCATCCAA | |
| PLD3 | ACTTGGCCCCGGTTCTATGA | qPCR primers |
| | ACCACGTTGAGTAGAGCCTTC | |
| GRN | GACGTGGAGTGTGGGGAAG | qPCR primers |
| | CCGATCAGCACAACAGACG | |
| LYZ | TGCCTGTCATTTATCCTGCAG | qPCR primers |
| | AACGATTTCTCCATGCCACC | |
| TIMP1 | AGAGACACCAGAGAACCCAC | qPCR primers |
| | ACGAACTTGGCCCTGATGA | |
| S100A8 | TCGACGTCTACCACAAGTACT | qPCR primers |
| | TCCTGGAAGTTAACTGCACCA | |
| CXCL8 | AGAGTGATTGAGAGTGGACCA | qPCR primers |
| | TGCACCCAGTTTTCCTTGG | |
| HLA-DRA | AGTCCCTGTGCTAGGATTTTTCA | qPCR primers |
| | ACATAAACTCGCCTGATTGGTC | |
| CST3 | CCAGCAACGACATGTACCAC | qPCR primers |
| | TCATGGAAGGGGCAGTTGTC | |
| HMOX1 | GTCAGGCAGAGGGTGATAGAA | qPCR primers |
| | GCTCTGGTCCTTGGTGTCAT | |
| TREM2 | GGTGGCACTCTCACCATTACG | qPCR primers |
| | CTCGAAGCTCTCAGACTCCC | |

| | | |
|---|---|---|
| CTSD | TGGACATCGCTTGCTGGAT<br>TTGGCTGCGATGAAGGTGAT | qPCR primers |
| IL-1B | CGATGCACCTGTACGATCAC<br>CATGGAGAACACCACTTGTTGC | qPCR primers |
| Fasn | AGACGCTTCTGGGCTACA<br>GTGTCCAGGGCAATGCTTG | qPCR primers for mouse *Fasn* mRNA detection |
| shFASN#1<br>shFASN#4<br>shFASN#5<br>shFASN#6<br>shGRN#1<br>shGRN#2<br>shIL-1B<br>shCTSD<br>shPLD3 | CGAGAGCACCTTTGATGACAT<br>GCTACGACTACGGCCCTCATT<br>GCTGCTGGAAGTCACCTATGA<br>CATGGAGCGTATCTGTGAGAA<br>GCTTCCAAAGATCAGGTAACA<br>GCTGTGAGGACCACATACACT<br>GGAGAGTGTAGATCCCAAAT<br>GCTGCACAAGTTCACGTCCAT<br>CGAGACCTGACCAAGATCTTT | shRNA sequences for human FASN, GRN, IL-1B, CTSD, and PLD3 |
| Fasn | TATTCTGGATCCTACTGCTTGGG<br>GGCTCGCTGGTCTTATACCATAC | Genotyping primers |
| Mx1-Cre | CGTACACCAAAATTTGCCTGC<br>CTAGAGCCTGTTTTGCACGTT | |
| sgFASN-#1<br>sgFASN-#2<br>sgFASN-#3<br>sgFASN-#4 | GGCCGCGGTTTAAATAGCGT<br>AGGCTGAAGCGCGGCGGAGA<br>AGCGGGAGGCTGAAGCGCGG<br>TCACGGACGATGACCGTCGC | sgRNA sequences targeting human FASN locus |

**Supplementary Table S4. Updated summary of FASN inhibitor** (*12*)**.**

| Compound | Potential targeted domains | Potency in biochemical assays (IC50) | Indication and/or preclinical effects | Cell lines | FASN protein levels upon treatment | Dosage and (or) treated time | Other application | Clinical trial ID or references |
|---|---|---|---|---|---|---|---|---|
| Orlistat | TE | 0.1 μM | Obesity | | | | | FDA approved for treating obesity |
| Orlistat | TE | 0.1 μM | Anti-cancer: Liver | Hep3B | down | 3, 10, 30 μM for 24hr | | (*13*) |
| Orlistat | TE | 0.1 μM | Anti-cancer: Colorectal, Prostate, Ovarian, Melanoma, Breast | PC-3, LNCaP, DU-145, B16-F10 | unchanged | 0.75, 3.0, 12.5, 25, 100μM for PC-3 | *in vivo* i.p. injection (240 mg/kg/day) | (*14*),(*15*),(*16*) |
| TVB-2640 | KR | 0.052 μM | Nonalcoholic steatohepatitis (NASH) | | | | | NCT04906421 (in progress, phase II); (*17, 18*) |
| TVB-2640 | KR | 0.052 μM | | | | | | NCT03808558, NCT03179904, NCT03032484, NCT02980029 (in progress, phase I/II) |

| | | | | | | | | |
|---|---|---|---|---|---|---|---|---|
| | | | | | | | | (*19, 20*) |
| FT-4101 | KR | 0.04 μM | nonalcoholic fatty liver disease (NAFLD) | | | | | (*21*) |
| FT113 | KR | 0.213 μM | Anti-cancer：Prostate, Breast | PC3, MV-411 | | | | (*22*) |
| BI-99179 | TE | 0.079 μM | Anti-cancer: glioma | GAMG | | | | (*23, 24*) |
| Cerulenin | KS | 4.5 μM | Anti-obesity | | | | | (*25*) |
| Cerulenin | KS | 4.5 μM | Anti-cancer: Melanoma, Cervical cancer | MDA-MB-453, SK-BR-3, | down | 2,4,8 μg/ml for 48hr | | (*26-28*) |
| C75 | KS | 15.5 μM | Anti-obesity | | | | | (*29, 30*) |
| C75 | KS | 15.5 μM | Anti-colitis | | down | Mouse colon tissue | DSS-induced colitis mouse model | (*31*) |
| C75 | KS | 15.5 μM | Anti-inflammation | HL-60 cell, RAW264.7, THP-1 | | | LPS-induced sepsis model (7.5 mg/kg IP) | (*32*) |

| | | | | | | | | |
|---|---|---|---|---|---|---|---|---|
| C75 | KS | 15.5 µM | Anti-cancer: Breast, Cervical,Prostate | Hela, CaSki, PC-3, LNCap | | | | (*28, 33, 34*) |
| Fasnall | KR/ER/MAT | 3.71 µM | Anti-cancer: Breast | MCF10A, MCF7, MDA-MB-468, BT474, SKBR3 | unchanged | 10 µM, 24hr | In vivo: 5,10,15mg/kg, ip, twice weekly | (*35*) |
| GSK2194069 | KR | 7.7 nM | Anti-cancer: Breast | KATO-III, MKN45, A549, SNU-1; GAMG cell (*36*); | unchanged | 10, 100, 1000 nM, 48h,120h | | (*37*) |
| GSK837149A | KR | 30 nM (Ki value) | | | | | | (*38*) |
| IPI-9119 | TE | 0.3 nM | Anti-cancer: Prostate | LNCaP, 22RV1, LNCap-95, | up | 0.05, 0.1, 0.25, 0.5 µM for 6 Days | | (*39*) |
| MP-ML-24-N1 | TE | 1.6 µM | Anti-cancer: Prostate | CAMG, LNCap | | | | (*36*) |
| TVB-3166 | KR | | NASH | | | | | (*18*) |
| TVB-3166 | KR | 0.042 µM | Anti-cancer: Lung, Ovarian, Prostate, | PANC-1, OVCAR-8, | unchanged or up (*40*) | 0.02, 0.2, 2 µM for 96h | In vivo, oral daily, 30, | (*40, 41*) |

| | | | | | | | | |
|---|---|---|---|---|---|---|---|---|
| | | | Pancreatic, colon | CALU-6, COLO-205, 22Rv-1, A549, HT-29 etc | (unchanged: OVCAR-8 Up: CALU-6, COLO-205, 22Rv-1) | | 60, 100 mg/kg | |
| TVB-3664 | | | NASH | | | | | (*18*) |
| TVB-3664 | | 0.018 μM | Anti-cacner: Hepatic, Lung, Ovarian, Prostate, Pancreatic | MHCC97H, HLE, SNU449 | | | | (*41, 42*) |
| PTM | ACP | 0.3 μM | NAFLD/diabetes | | | | | (*43-45*) |
| PTM-6p | KS-MAT didomain | - | Anti-cacner: non-small-cell lung cancer, menaloma | A549/NCI-H1299 cells | down (NCI-H1299 cells) | 5/10/25/50 μM for 48h | | (*46*) |
| MS-C19 | | | Anti-cancer: Acute myeloid leukemia (AML), Chronic myeloid leukemia (CML) | MOLM-13, K562, NB4 | down (MOLM-13) unchanged (K562) | 1,2.5,5,10μM for 48hr or 72hr | | this study |
| EGCG | KR | 52 mM | | | | | | (*47*) |
| Resveratrol | KR | 11.1 μg/ml (FASN) and 21.9 μg/ml (KR domain) | Anti-cancer: Breast | SKBR-3 | down | 20,40,60 μM for 24h | | (*48, 49*) |

| | | | | | | | |
|---|---|---|---|---|---|---|---|
| Triclosan | ER | 5-10 μM | Anti-cancer: Prostate | LNCaP/22RV1/PC-3/C4-2B/LAPC4 | down | 10μM for 48h, 72h | (*50, 51*) |
| AZ12756122 | | ~60μM | Anti-cancer：Lung | PC9-GR4 | down | 25μM for 72h | (*52*) |
| G28UCM | | | Anti-cancer：Breast | AU565 | unchanged | 30μM for 12, 24, 48h | (*53*) |
| Icaritin | | docking data, no direct experimental evidence | Anti-cancer：melanoma | HS68, A375S, A375R, NEW0, A2058; | down | 20,40,80 μM for 24h | (*54*) |
